# Structures of aberrant spliceosome intermediates on their way to disassembly

**DOI:** 10.1101/2024.09.13.612651

**Authors:** Komal Soni, Attila Horvath, Olexandr Dybkov, Merlin Schwan, Sasanan Trakansuebkul, Dirk Flemming, Klemens Wild, Henning Urlaub, Tamás Fischer, Irmgard Sinning

**Author notes:** Correspondand requests for materials should be addressed to I.S. or T.F.

## Abstract

Intron removal during pre-mRNA splicing is of extraordinary complexity and its disruption causes a vast number of genetic diseases in humans^1^. While key steps of the canonical spliceosome cycle have been revealed by combined structure-function analyses^2,3^, structural information on an aberrant spliceosome committed to premature disassembly is not available. Here, we report two cryo-EM structures of post-B^act^ spliceosome intermediates from *S. pombe* primed for disassembly. We identify the DEAH-box helicase – G patch protein pair (Gih35-Gpl1, homologous to human DHX35-GPATCH1) and show how it maintains catalytic dormancy. In both structures, Gpl1 recognizes a remodeled active site introduced by an over-stabilization of the U5 loop I interaction with the 5’ exon leading to a single nucleotide insertion at the 5’splice site. Remodeling is communicated to the spliceosome surface and the Ntr1 complex that mediates disassembly is recruited. Our data pave the way for a targeted analysis of splicing quality control.

---

Pre-mRNA splicing is performed by the multi-subunit and highly dynamic ribonucleoprotein particle known as the spliceosome^4–6^, where two transesterification reactions, branching and exon ligation, produce the mature mRNA (reviewed in^2,7^). Initial steps of spliceosome assembly comprise recognition of the 5’splice site (ss), branch-site (BS) and 3’ss leading to the formation of an early complex (E) which transitions into the pre-spliceosome complex (A) and subsequently the fully assembled precursor spliceosome complex (pre-B). The spliceosome then transforms into the activated (B^act^) and catalytically activated (B*) forms before the first transesterification reaction, the step I (C) and step II activated complexes (C*) before the second transesterification reaction, and finally into the postcatalytic (P) and the intron lariat spliceosome (ILS), which lead to spliceosome disassembly for further rounds of processing.

This sequential remodeling of the spliceosome is driven by eight conserved DEAD/H RNA helicases^8,9^, a subset of which (namely Prp5^10^, Prp2^11,12^, Prp16^13,14^ and Prp22^15^) also guarantee splicing fidelity by actively promoting discard of aberrant RNA substrates as part of an internal splicing quality control mechanism^16^. In addition to these, the RNA helicase Prp43 is responsible for dismantling the ILS to recycle the small nuclear ribonucleoproteins (snRNPs)^17–20^. Prp43 and Prp2 work together with their respective G-patch protein co-activators Ntr1^18,21^ and Spp2^22,23^. G-patch proteins are defined by a ∼50 amino acid long glycine-rich motif domain^24^. They are conserved in many RNA-processing proteins and serve as critical cofactors of RNA helicases^24,25^. While the action of Prp2-Spp2 triggers remodeling of the spliceosome from the B^act^ to B* complex^26–28^, Prp43 together with the G-patch motif of Ntr1 can disassemble the ILS complex^29^. However, Prp43 is additionally responsible for the discard of spliceosomes stalled at stages dependent on Prp16 and Prp22^14,30–33^. To selectively discard only ILS and defective spliceosomes and prevent disassembly of properly assembled spliceosomes, Prp43 has a second associated protein Ntr2^18,34^, which together with the C-terminal domain of Ntr1 (Ntr1 CTD) act as doorkeepers for a productive disassembly of only the ILS and defective spliceosomes^35^. Prp43 and its cofactors Ntr1, Ntr2, and the stabilizing Cwc23, together form the Ntr1 complex.

After the determination of the first near-atomic cryo-electron microscopy (cryo-EM) structure of the ILS spliceosome from the fission yeast *Schizosaccharomyces pombe (sp)* in 2015^36^, all major functional states of the spliceosome from *Homo sapiens* (*hs*) and *Saccharomyces cerevisiae (sc)* have been structurally characterized (reviewed in^3^), now providing detailed mechanistic understanding of the canonical pre-mRNA splicing. However, a high-resolution structure of a defective spliceosome intermediate committed to premature disassembly providing insights into such a splicing quality control mechanism is not available to date.

## Isolation and characterization of a defective spliceosome intermediate

Recent studies have found an association of the evolutionarily conserved protein Nrl1 (homologous to *hs*NRDE-2) with components of the splicing machinery^37–39^, where it is part of a sub-complex known as the CNM (named after the proteins Ctr1-Nrl1-Mtl1) that targets unspliced pre-mRNAs to the exosome for degradation during RNA surveillance^39,40^. In fact, tandem affinity purifications (TAP) using Nrl1 as bait consistently co-purified the spliceosome disassembly factors Prp43, Ntr1 and Ntr2 in high abundance^37,38,40^. A similar protein interactome was identified using reciprocal purifications with Ntr1 and Ntr2 as bait proteins^41^.

We reasoned that a split-tag approach using affinity tags on Nrl1 and Prp43 might yield an enrichment of a spliceosome intermediate assembled on unspliced pre-mRNAs on its way to the discard pathway for subsequent cryo-EM analysis. Indeed, such an approach led to co-purification of Nrl1 associated proteins and the Ntr1 complex (**Extended Data Fig. 1**, *S. cerevisiae* nomenclature is used for *S. pombe* proteins wherever homologous proteins exist; see **Extended Data Table 1** for *S. pombe* protein names). We also identified the DEAH-box helicase Gih35 and the G-patch domain containing protein Gpl1 consistent with previous reports, where Gih35 co-purified with Gpl1 in TAP purifications^41^ and other splicing factors^36,40,42^. After judging the homogeneity of the complex using negative staining, we subjected the complex to cryo-EM. Approximately, 0.93 million particles were auto-picked from 13,000 micrographs and after 3D classification 0.25 million particles yielded a reconstruction with additional cryo-EM density near the RNaseH-like domain of Prp8 (Prp8^RH^) at the periphery of the spliceosome. Several rounds of focused classification finally yielded two reconstructions of a post-B^act^ spliceosome primed for discard (B^d^) at average resolutions of 3.2 Å and 3.1 Å (**Extended Data Fig. 2**).

## Overall architecture of a spliceosome intermediate primed for discard

The two *sp*B^d^ structures are similar with a few compositional differences which represent two distinct states -defined as here *sp*B^d^-I and *sp*B^d^-II. We first describe their similarities and later discuss differences between the two states. The overall architecture of the *sp*B^d^ complex (referring to both *sp*B^d^-I and *sp*B^d^-II) resembles a combination of B* and ILS complexes (**Fig. 1**, **Extended Data Table 2**). The *sp*B^d^ complex comprises the stable core of the spliceosome including the U5 snRNP, U6 snRNA, U2/U6 duplex, Prp19 complex (NTC) and the NTC related (NTR) proteins, which all remain largely unchanged in the B^act^ to ILS complexes^3^ as also in the *sp*B^d^ complex (**Extended Data Fig. 3a**). In addition to the core, we find the 5’-exon stabilizing factors Cwc21 and Cwc22, which are recruited into the B^act^ complex and dissociate at the P to ILS transition^43^ (**Extended Data Fig. 3b**). Moreover, Cwf11 (homologous to Aquarius in humans) is present in the *sp*B^d^ complex, a protein which is required for the transition from B^act^ to B* during catalytic activation in humans^44^. The U2 snRNP comprises a 5’-domain (SF3b complex), a 3’ domain (or core domain containing the Sm ring, Msl1 and Lea1) and the SF3a complex that bridges both the domains. While the SF3a/b complex is released during the B^act^ to B* transition, the core domain remains bound although it undergoes dramatic translocation during the B^act^ through ILS complex transitions. However, in our structures the U2 core domain remains unidentified (**Extended Data Fig. 3b**). Most importantly within the *sp*B^d^ complex, aided by crosslinking mass spectrometry (**Extended Data Fig. 4**), we have localized the helicase Gih35 and its associated G-patch containing protein Gpl1 in our structure (**Fig. 1**).

**Fig. 1:**
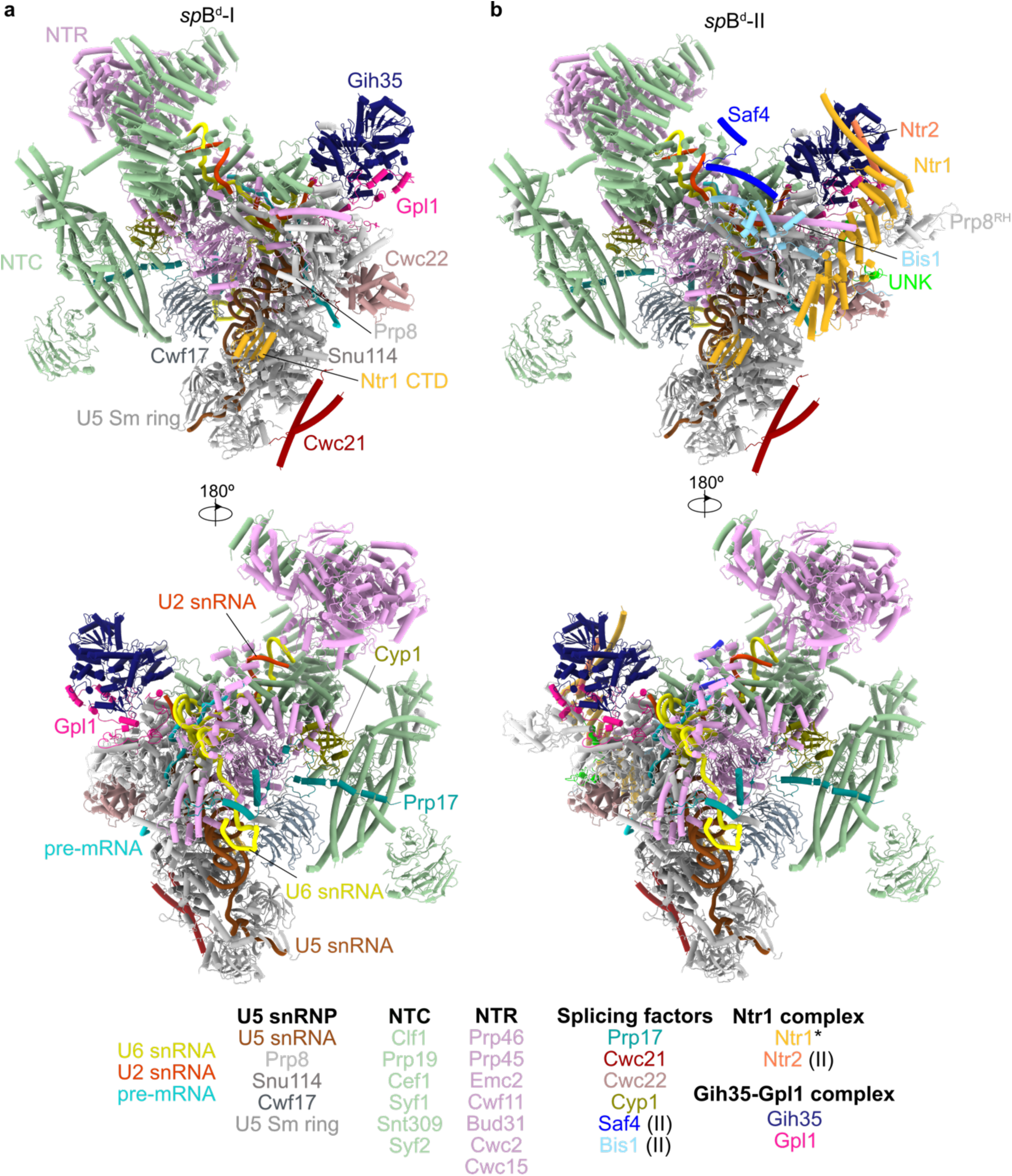
Cryo-EM structure of the *sp*B^d^ complex. **a** and **b,** Structures of the *sp*B^d^-I and *sp*B^d^-II complexes at average resolutions of 3.2 and 3.1 Å, respectively. Two views of the *sp*B^d^ complexes are shown, colored according to the different subunits which are listed below the structure. The Ntr1 protein is marked (*) and is only partially visible in the *sp*B^d^-I state while stabilized in the *sp*B^d^-II state. Protein components that are only part of the *sp*B^d^-II state are indicated.

Despite the overall similarity of the two *sp*B^d^ structures there are significant differences. In the *sp*B^d^-I state, the cryo-EM map for two components of the Ntr1 complex, namely Ntr1 and Ntr2, is very weak in comparison to *sp*B^d^-II complex and therefore they are not built in the *sp*B^d^-I structure. As an exception to this, the Ntr1 CTD which is stably bound in both states. Most significantly however, splicing factors Bis1 (homologous to human ESS2^45^) and Saf4 are only present in the *sp*B^d^-II state. Saf4 and Cwf16 both share sequence homology with the *S. cerevisiae* step I splicing factor *sc*Yju2, similar to human CCDC94 and CCDC130 (**Extended Data Fig. 5a-b**). We find the *sp*B^d^-II cryo-EM map to be consistent only with Saf4 suggesting Saf4 to be the direct homologue of human CCDC94 and *sc*Yju2 rather than Cwf16 as suggested previously (^46^ and **Extended Data Fig. 5c**). In addition, two short α-helical fragments of the NTC core components Cef1 and Syf2 are destabilized in the *sp*B^d^-II state to accommodate Saf4 and Bis1 (**Extended Data Fig. 5d-f**). The RNaseH-like domain of Prp8 (Prp8^RH^) is also only visible in the *sp*B^d^-II state (**Fig. 1**). Finally, well-defined density is found for multiple helices around Prp8 and Cwc22 in the *sp*B^d^-II cryo-EM map, the identity of which however could not be clarified (**Extended Data Fig. 6**) and is henceforth referred to as unknown (UNK).

Altogether, we have modelled 30 (*sp*B^d^-I) and 33 (*sp*B^d^-II) proteins and four RNA molecules in the *sp*B^d^ cryo-EM maps (**Extended Data Table 1**). It is noteworthy that we do not find density corresponding to the CNM complex in either of the *sp*B^d^ cryo-EM maps although intermolecular crosslinks between Ctr1 and parts of Cwc22 (not built) are observed (**Extended Data Fig. 4**), indicating that the CNM is likely located at the periphery and therefore flexibly linked.

## The Ntr1 complex is stabilized in the *sp*B^d^-II complex

Out of the four components of the Ntr1 complex, we observe defined density only for Ntr1 and Ntr2 in the *sp*B^d^-II cryo-EM map while Prp43 and Cwc23 remain unidentified. The positions of Ntr1 and Ntr2 proteins at the periphery of the spliceosome are similar to those described for the *sc*ILS complex (**Extended Data Fig. 3b and 47**). Ntr1 consists of three distinct domains, namely the G-patch motif domain, a central superhelical domain and the CTD (**Extended Data Fig. 7a**). While the Ntr1 G-patch motif domain is not found in both *sp*B^d^ states, the CTD is bound in both the *sp*B^d^ states where it is anchored on Snu114 as observed in the *sc*ILS complex (**Extended Data Fig. 3b, 7b and 47**). In the *sp*B^d^-II state, the superhelical domain contacts Ntr2, Prp45 and Bis1, and approaches Gih35 (**Extended Data Fig. 7c-d**) while in the *sp*B^d^-I state, the domain has weak cryo-EM density and was therefore not built. This flexibility of the Ntr1 superhelical domain in the *sp*B^d^-I state seems due to the absence of the two proteins Saf4 and Bis1 that stabilize the domain in the *sp*B^d^-II state (**Extended Data Fig. 7e**). For Bis1, so far only the helical bundle of its human homolog ESS2 could be localized in the *hs*C* complex (**Extended Data Fig. 3b and 45**). In addition to the helical bundle in Bis1, we now find a continuous stretch of ∼80 residues (180-262) crawling across the surface of Prp8, Prp46 and Cwc15, part of which is directly interacting with the Ntr1 superhelical domain and anchoring it to Prp8 in the *sp*B^d^-II state (**Extended Data Fig. 7e-f**).

In the *sc*ILS complex, the Ntr1 CTD is further stabilized by an α-helix of Cwc23, which is not found in the *sp*B^d^ complex (**Extended Data Fig. 7g**). However, we do observe crosslinks between Cwc23 and the linker connecting the Ntr1 superhelical domain with the CTD, suggesting that Cwc23 is flexibly bound in the *sp*B^d^ complex (**Extended Data Fig. 4**). Similarly, crosslinks between the Ntr1 G-patch domain and Prp43 suggest that Prp43 is also flexibly tethered to the *sp*B^d^ complex (**Extended Data Fig. 4**). Ntr2 has been suggested to stabilize Ntr1 on defective spliceosomes to promote disassembly by Prp43^35^. While we observe only a small α-helical fragment of Ntr2 in the *sp*B^d^-II state (**Extended Data Fig. 7c-d**), it is positioned in close proximity to Gih35 where it may play a role in stabilization of Gih35 (**Extended Data Fig. 7h**). Taken all interactions and component together, the *sp*B^d^ structures represent post-B^act^ spliceosome intermediates captured on their way to disassembly.

## Gpl1 is anchored to the spliceosome by Prp8

Gpl1 has been recently characterized as a splicing regulator in *S. pombe*, necessary for the recruitment of the Gih35 helicase^48^ and proper canonical splicing^48,49^. It is a G-patch domain (residues 137-181) containing protein comprising 534 amino acids of which about 200 residues from the N-terminal part (residues 25-188, and 199-234) can be traced in our complex (**Extended Data Fig. 8a, b;** for structural illustrations of redundant regions between the two *sp*B^d^ states of the *sp*B^d^ complex, only the higher resolution *sp*B^d^-II state is further described unless mentioned otherwise). Our data show how Gpl1 recruits the DEAH-box helicase Gih35 to the spliceosome (**Fig. 2a**) and interacts exclusively with different domains of Gih35 and Prp8 (**Fig. 2b, c**).

**Fig. 2:**
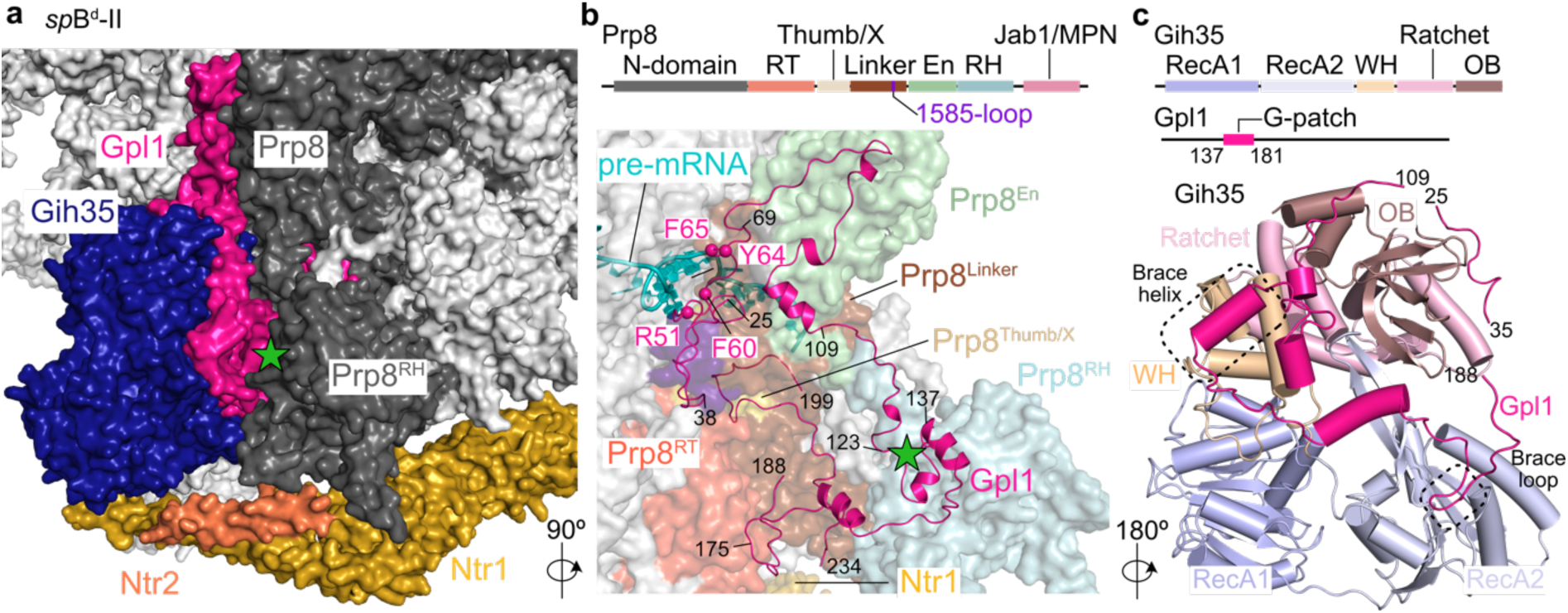
Interactions between Gpl1, Gih35 and the spliceosome. **a,** Gpl1 binds to Prp8 and Gih35 and bridges between the two proteins. **b,** Gpl1-Prp8 interaction, Top: Schematic representation of the domain architecture of Prp8. Bottom: Gpl1 contacts the Prp8 RT, Thumb/X, Linker (including the 1585-loop), En and RH domains. Residues contacting the pre-mRNA in the active site of the spliceosome are indicated. The Gpl1 knob (residues 123-137) is marked with a green star in panels (a) and (b). **c,** Gpl1-Gih35 interaction, Top: Schematic representation of the domain architectures of Gih35 and Gpl1. Bottom: Overview of the Gih35-Gpl1 interactions. The positions of the Gpl1 brace helix and brace loop are marked.

Overall, the N-terminal part of Gpl1 forms a remarkably large interface with Prp8 of 5,600 Å^2^. In detail, a region of Gpl1 (residues 38-69) which is fixed on the Linker domain of Prp8 (Prp8^Linker^, including the 1585-loop of Prp8, also known as α-finger-residues 1537-1550) also reaches into the active site of the spliceosome where it contacts the pre-mRNA with a cluster of aromatic and charged residues (region 51-65, as detailed below) (**Fig. 2b**). Next, Gpl1 residues 70-109 are anchored on the Endonuclease domain of Prp8 (Prp8^En^) where Gpl1 makes a sharp turn to cover a large surface area of interaction (**Fig. 2b**, **Extended Data Fig. 8c**). Furthermore, Gpl1 residues 123-137 form a knob that inserts into Prp8^RH^ (**Fig. 2a, b** and **Extended Data Fig. 8d**). It is known that Prp8 rearranges throughout the splicing cycle with the Prp8^RH^ and the Prp8 Jab1/MPN domain (Prp8^Jab1/MPN^) displays different positions with respect to the core domains and contributes to stabilization of the exchanging spliceosome proteins and branch helix movements^50^. While Prp8^Jab1/MPN^ remains invisible in both *sp*B^d^ complexes, Prp8^RH^ is observed only in the *sp*B^d^-II state where it adopts by far the most open conformation (**Extended Data Fig. 9a**). During the *sc*B^act^ to *sc*B* transition, the Prp8^RH^ rotates by 145° to contact and stabilize the branch helix, while it undergoes a -35° rotation in the opposite direction during the *sc*B^act^ to *sp*B^d^ transition (**Extended Data Fig. 9b**). This extended open conformation of Prp8^RH^ is fixed by its interaction with the Gpl1 knob only in the *sp*B^d^-II state (**Extended Data** Fig. 8d). Moreover, Gpl1 residues 175-188 are part of a small and rather hydrophobic interface with the Prp8 Reverse Transcriptase domain (Prp8^RT^) (**Fig. 2b**, **Extended Data Fig. 8e)**. Finally, Gpl1 residues 199-234 insert into a central Prp8 cavity created by its RT, Thumb/X and Linker domains (**Fig. 2b**, **Extended Data Fig. 8f**). This stretch of Gpl1 occupies the same general location as components of Prp45 in the *sc*B^act^ or Ntr2 in the *sc*ILS states (**Extended Data Fig. 10**).

## Gpl1 binds to Gih35 and tethers it to the spliceosome

Gih35 is a canonical DEAH-box RNA helicase with a conserved core comprising the RecA1 (residues 20-202) and RecA2 (residues 203-379) domains that is flanked by the C-terminal WH (residues 380-447), Ratchet (residues 448-557), and OB domains (residues 558-647) (**Fig. 2c**). In the *sp*B^d^ complex, Gpl1 tethers Gih35 to the periphery similar to other helicases known to remodel the spliceosome including Prp2^51–53^, Prp16^46,54,55^ and Prp22^56–61^ (**Extended Data Fig. 11**). Gpl1 shares an interface of >2,500 Å^2^ with Gih35, similar to Prp2, which is bound to the *sc*B^act^ complex via Spp2^53^. Two stretches of Gpl1 form distinct interactions with Gih35. While Gpl1 residues 25-35 insert in a groove between Gih35 WH and OB domains, residues 109-181 including the G-patch (137 to 181) traverse along the OB, WH and RecA2 domains forming an extended interaction. The N-terminus of the G-patch forms an α-helix (residues 139-148; termed as brace helix^62^) and the C-terminal loop (termed as brace loop^62^) inserts into a hydrophobic pocket on top of the RecA2 domain (**Fig. 2c**, **Extended Data Fig. 12a-d**). Superpositions of Gih35-Gpl1 with other known DEAH-helicase – G-patch co-activators, such as the Prp2-Spp2 pair from *Chaetomium thermophilum* (*ct*)^63^ and DHX15 with its G-patch co-factor NKRF^63^, show that the positions of the brace helix and brace loop are fixed while the linker region connecting the N-and C-terminal parts of the G-patch domain varies (**Extended Data Fig. 12e-f**). In accordance, the sequence alignment of Gpl1 with other G-patch proteins that are known activators of Prp2/DHX16 or Prp43/DHX15 shows a high degree of conservation only in the brace helix and brace loop regions while the connecting linker is poorly conserved (**Extended Data Fig. 12a**). Overall, apart from the eponymous glycines of the motif, only hydrophobic residues within the brace helix and brace loop are maintained. Specificity of the helicase – G-patch pair is likely to be ascribed to the non-conserved regions like the brace-linker, comprising a unique α-helix in Gpl1 (**Extended Data** Fig. 12c-d), and/or N-terminal regions, which form additional contacts outside the G-patch. To ascertain whether Gpl1 binds specifically to Gih35 or does not discriminate between other DEAH-helicases such as Prp43, we performed yeast-two-hybrid experiments (**Extended Data Fig. 12g**). While Gpl1 binds to Gih35 as also shown previously^48^, it does not bind to Prp43. Similarly, Ntr1 (the *bonafide* interaction partner of Prp43) does not bind to Gih35 but interacts with Prp43 as expected. In accordance, the Ntr1 G-patch domain co-migrates with Prp43 in size-exclusion chromatography experiments, indicative of complex formation, while the Gpl1 G-patch domain does not (**Extended Data** Fig. 12h). These data underline the specific interaction of the Gih35-Gpl1 pair.

In contrast to Gpl1, interactions of Gih35 with the spliceosome are sparse. Only the RecA2 domain of Gih35 is placed in a pocket formed by Prp8^RT^, NTC proteins Cef1 and Clf1, a helix of an unknown protein and Ntr1 and Ntr2 (in case of *sp*B^d^-II state) (**Extended Data Fig. 12i**). In summary, the Gih35-Gpl1 pair is anchored on the spliceosome in a two-pronged manner with Gpl1 interacting extensively and reaching deep into the active site, and with Gih35 recruited piggy-back by stabilizing contacts to the spliceosome only via its RecA2 domain.

## Canonical and non-canonical features of the *sp*B^d^ active site

The comparison of snRNAs from canonical spliceosomes in catalytic states (B^act^ to ILS complexes) with the aberrant *sp*B^d^ complexes shows that RNA elements at the active site superimpose well (**Extended Data Fig. 13**). The U5/5’-exon duplex, U6 internal stem loop (ISL), and U2/U6 helices Ia, Ib, II are also well defined in the *sp*B^d^ structure, while the ACAGA helix is less defined (**Extended Data Fig. 14a**). We find three structural metal ions at the active site, likely Mg^2+^ ions which are exclusively coordinated by the U6 ISL (**Extended Data Fig. 14b-c**). While the catalytic metal ions M1 and M2 have not been incorporated into the *sp*B^d^ states, the K1 (potassium) site^64^ is occupied (**Fig. 3a**, **Extended Data Fig. 14c**). Strikingly, we find a tight RNA duplex formed between the U5 snRNA and the 5’-exon with Watson-Crick base pairing between U5 loop I nucleotides U_50_U_51_U_52_ and 5’-exon nucleotides A_-2_A_-3_A_-4_ and optimized packing of A_-5_ with A_53_ (**Extended Data Fig. 15a**). The pairing is similar to a cognate codon-anticodon mRNA-tRNA interaction.

**Fig. 3:**
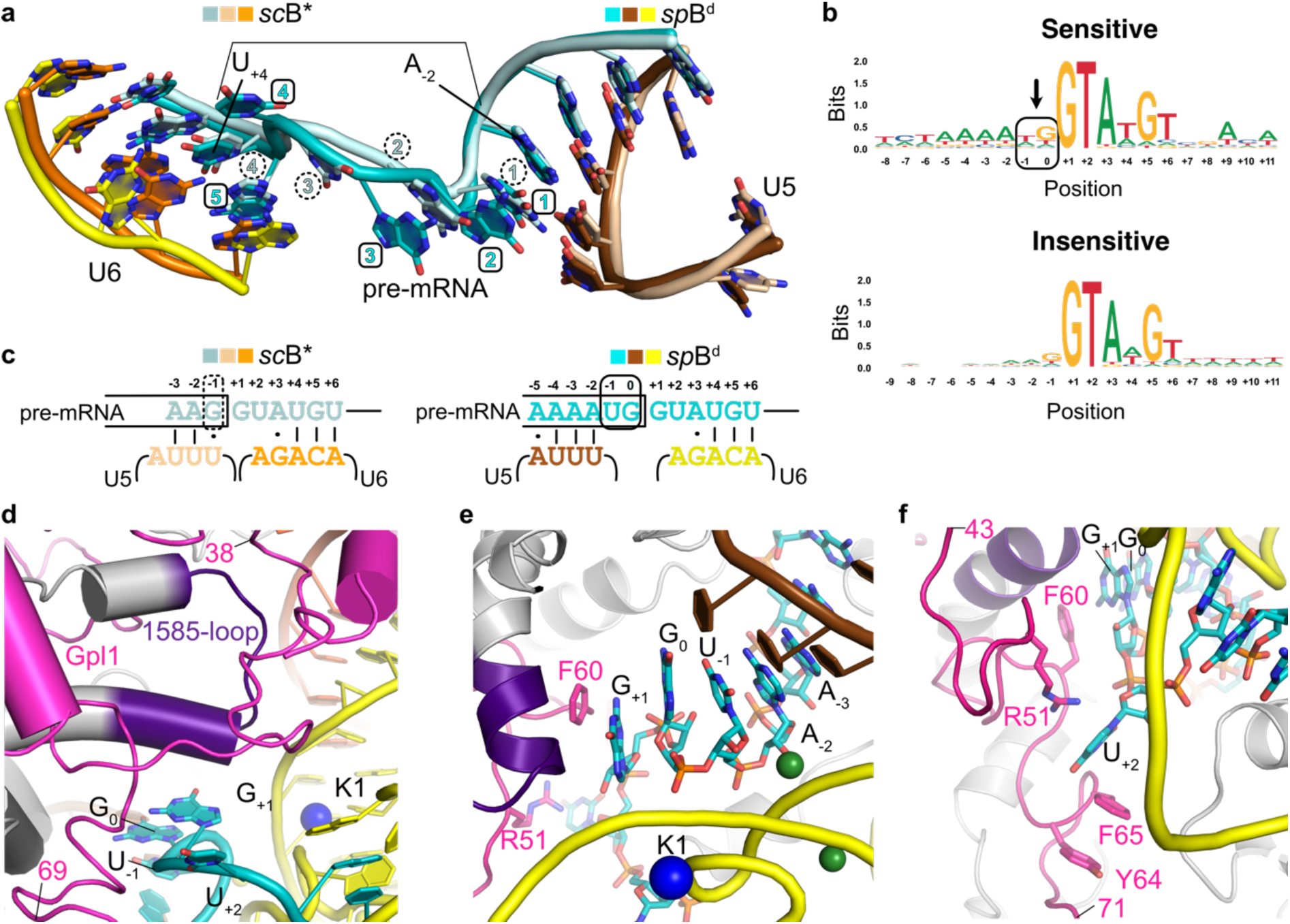
Architecture of the active site in the *sp*B^d^ complex. **a,** Superposition of RNA elements (pre-mRNA, U5 and U6) at the active site of *sp*B^d^ and canonical *sc*B* (PDB:6J6Q) complexes. The bracket marks the positions of pre-mRNA bases A_-2_ and U_+4_ which is covered by 4 nucleotides in *sc*B* (marked with dashed circles) and 5 nucleotides in *sp*B^d^ (boxed in squares). **b,** 5’ss sequence logo generated from CNM-sensitive (top) and CNM-insensitive (bottom) transcripts after normalization by gene expression. The insertion at -1/0 position is marked with an arrow. **c,** Scheme of RNA elements at the active site of *sp*B^d^ using the 5’ss sequence logo from panel b and canonical *sc*B* (PDB:6J6Q) complexes. The insertion at the -1/0 position is marked with a continuous box in *sp*B^d^ complex while the G_-1_ position in *sc*B* complex is marked with a dashed box. **d,** Gpl1 binds at the heart of the spliceosome where it interacts with the 5’ss and the 1585-loop of Prp8 (purple). Color scheme for protein and RNA elements follows Fig.1. **e,** Gpl1 residue F60 stacks with G_+1_ of the 5’ss. **f,** Rotated view showing recognition of U_+2_ of the 5’ss by Gpl1.

Most significantly we find that the 5’ss contains a single nucleotide insertion (**Fig. 3a**). It is important to mention that while the cryo-EM density for snRNAs at the active site is very well defined, that for the pre-mRNA at the 5’ss is weak in comparison causing identification of bases or the conformation beyond the -1 position inaccurate (**Extended Data Fig. 14c-d, 15a**). This is comprehensible, given that the *sp*B^d^ complex is an affinity-purified *in vivo* complex and therefore incorporates a mixture of RNA species rather than being assembled *in vitro* on a single RNA. Nevertheless, the presence of the insertion is unambiguous (**Fig. 3a**, **Extended data Fig. 14d**). Together with the assumption that the canonical base pairing in yeast spliceosomes between the U6 ACAGA box with the intron positions +4 to +6 is preserved, the strong U5 loop I interaction with the 5’-exonic Adenine quadruple (positions -2 to -5) induces a defined register constraint on the pre-mRNA. In effect, the distance between the pre-mRNA -2 and +4 positions cannot be bridged by the canonical four nucleotides but needs five nucleotides instead (**Fig. 3a**). To accommodate the insertion at the active site, the pre-mRNA is squeezed introducing a bend, which alters its overall geometry (**Fig. 3a**, **Extended Data Fig. 15b**).

## RNA sequencing provides insights into pre-mRNA targets of the *sp*B^d^ complex

To validate our structural finding of an insertion at the pre-mRNA 5’ss pre-mRNAs, we wanted to ascertain whether all pre-mRNAs bound in the *sp*B^d^ complex generally harbor the exonic quadruple Adenines (positions -2 to -5), and whether this leads to the single nucleotide insertion at the 5’ss, which could be a hallmark of transcripts targeted for discard via the CNM. We have previously reported that compared to the wild-type strain, strains harboring deletions of CNM components (*ctr1Δ, nrl1Δ*) showed accumulation of unspliced pre-mRNAs^39^. We re-analyzed the RNA-sequencing data for *ctr1Δ*, which previously showed the highest accumulation of unspliced pre-mRNAs^39^ (**Extended Data Fig. 16a-c**). Using a sensitivity-metric wherein introns with at least 20% increased accumulation in the *ctr1Δ* strain compared to wild-type were defined as introns sensitive to CNM-mediated discard (**Extended Data Fig. 16d**), we found that an exonic quadruple Adenine motif is slightly enriched in the sensitive transcripts compared to the insensitive ones (**Extended Data Fig. 16e**). Normalization of RNA species according to abundance led to the emergence of a more prevalent exonic quadruple Adenine motif (A_-2_A_-3_A_-4_A_-5_) (**Fig. 3b**, **Extended Data Fig. 16f)**. Strikingly, the motif shows the presence of five nucleotides between the A_-2_ and U_+4_ position (and therefore an insertion) in the sensitive transcripts which is in agreement with the cryo-EM map. Within the intron, strong conservation is found for nucleotides canonically interacting with the ACAGA box of U6 snRNA (A_+3_U_+4_G_+5_U_+6_) and the universal G_+1_U_+2_ motif is also strictly conserved. Therefore, to accommodate two nucleotides between the quadruple Adenine motif and the conserved G_+1_ position, we define a new position ‘0’ (**Fig. 3b-c**, **Extended Data Fig. 15c)**. The RNA-seq analysis shows that the likely gene candidate, which is highly expressed and therefore majorly contributing to the structural observation of these features, is *rpl39* encoding for the 60S ribosomal protein eL39 (**Extended Data Fig. 16g-h**).

To further validate these observations, we isolated and sequenced the co-precipitating RNA transcripts from our *sp*B^d^ spliceosome purification. These RNA immunoprecipitation (RNA-IP) experiments confirmed that our purifications strongly enrich intronic sequence coverage (**Extended Data Fig. 17a,b)**. Most intronic reads in affected introns overlap with exon-intron boundaries, indicating that they represent un-spliced and un-cleaved pre-mRNAs prior to the first catalytic step of the splicing reaction (**Extended Data Fig. 17a)**. Determination of the abundance of individual introns proved to be highly complex, due to the large amount of contaminating RNA molecules in these samples, which constituted the majority of reads in these RNA-IP experiments. Nevertheless, after filtering for enriched intronic sequences in the IP fraction and excluding mis-annotated, alternatively spliced and retained introns, the most abundant intronic sequence was the intron in the *rpl39* gene (**Extended Data Fig. 17c)**. Of note, contaminating RNA molecules that are present in our affinity purification are not incorporated into the protein complex and therefore do not appear or pose a problem in structural analysis.

Overall, these results further confirm that the pre-mRNA substrate observed in the *sp*B^d^ complex is likely *rpl39*, which is probably the most abundant substrate for the *sp*B^d^-mediated splicing quality control in *S. pombe*.

## The aberrant active site in the *sp*B^d^ complex

Following the validation of the insertion in the pre-mRNA at the active site, we proceeded with investigation of additional changes and alterations at the active site. Remarkably, we find that the Gpl1 N-terminal region (residues 38-69) closely engulfs the active site (**Fig. 3d**). The nucleotides of the 5’ss, including U_-1_, G_0_ and G_+1_, stack with each other to form a continuous ladder that is further extended by Gpl1 Phe60, which stacks on top (**Fig. 3e**, **Extended Data Fig. 15d-f**). Due to the insertion, the canonical U_+2_ is also positioned differently and is sequestered by Gpl1 in a pocket formed by residues Arg51, Try64 and Phe65 (**Fig. 3f**, **Extended Data Fig. 15g**). The presence of Gpl1 also likely occludes the docking of the U2/BS duplex at the active site, which remains invisible in the *sp*B^d^ complex.

The positioning of Gpl1 at the active site somewhat mimics that of Prp11 and Cwc24 in the B^act^ complex, where they shield the 5’ss and thus maintain catalytic dormancy (**Extended Data Fig. 18a**). The conformation of the pre-mRNA at the active site and the simultaneous binding of Gpl1 are both mutually exclusive with Prp11/Cwc24 accommodation (**Fig. 3e**, **Extended Data Fig. 18a-b**). The 1585-loop which helps to maintain catalytic dormancy at the B^act^ state by interacting with the N-terminus of Prp11, Cwc24 and the U2/U6 helix Ia^52^ (**Extended Data Fig. 18c**), is also structured in the *sp*B^d^ complex (**Fig. 3d**, **Extended Data Fig. 18d**). However, in the *sp*B^d^ complex, it is located in a position similar to that observed in the *sc*C* complex^55^ (**Extended Data Fig. 18e-f**). While in the *sc*C* complex, the 1585-loop stabilizes the active site by interacting with the U6 ISL and the lariat junction, in the *sp*B^d^ complex it is sandwiched between Gpl1 residues 38-60 on the one side and residues 204-209 on the other (**Extended Data Fig. 18e-f**). Further, it also interacts with U2/U6 helix Ia, the stacked intronic G_+1_, and Gpl1 Phe60 (**Fig. 3e**, **Extended Data Fig. 18f**). Interestingly, the N-terminus of the NTR protein Cwc15, which cradles the AGC triplex (**Extended Data Fig. 18g**) that coordinates the central Mg^2+^ ions at the active site in the *sc*B* complex but not in the *sc*B^act^ complex, is likewise disordered in the *sp*B^d^ complex. This disordering coincides with continuous stacking of U2/U6 helix Ib on the U6 ISL (U2 G_19_ on U6 C_73_), thus closing an entrance gate for Cwc15 towards the AGC triplex (**Extended Data Fig. 18g, h**).

Not all pre-mRNAs targeted by the Nrl1-containing CNM complex harbor the quadruple Adenine motif at the 3’-end of the 5’-exon, however, *rpl39* serves as an example of an aberrant splice site leading to the recruitment of Gpl1 (and Gih35). Given that the spliceosome is known to adopt different substrate-specific conformations^65^, the mechanism of Gpl1 recruitment and its conformation at the active site is likely to be substrate-specific and depends on the type of aberration. Likewise, the base pairing between 5’ exon and 5’ ss to U5 loop I and U6 snRNA for different CNM-targeted transcripts and the mRNA loop conformation in between may also differ from our proposed structural model.

Usually during the formation of the active site, the U5 loop I pairs with the 5’ exon sequence immediately upstream of the 5’ss^66,67^, holding the 5’ exon in place. The presence of four adenines at positions -2 to -5, which pair with U5 loop I, and the ACA sequence within the U6 ACAGA box, which alike base pairs with U_+4_G_+5_U_+6_ of the intron^68,69^, form strong anchor points, which cause insertion of a single nucleotide at the active site. Taken together, this constraint seems sufficient for formation of an aberrant active site preventing the canonical progression of splicing.

## Discussion

While the recent progress in our structural and functional understanding of the splicing machinery is enormous, this information is still lacking for defective spliceosome intermediates committed to premature disassembly. The necessity to close this gap is even more relevant, as mis-splicing underlies a growing number of human diseases with substantial societal consequences^1^. Our structures provide a snapshot of such a scenario, where the overstabilization of the U5 loop I interaction with the 5’ exon and the U6 interaction with the intron leads to an extra nucleotide insertion at the active site, rendering the spliceosome defective and committed to discard (**Fig. 4**). Normally, during transition from the B^act^ to the B* complex in a productive splicing cycle, the U2 snRNP SF3a/b complex, RES complex, and the splicing factors Cwc24 and Cwc27 dissociate to release the 5’ss and the branch helix as a result of Prp2 action in yeast^65^ or the combined action of Prp2 and Cwf11/Aquarius in humans^70^. At an intermediate step, before the recruitment of the branching factors in the B* complex, the branch helix is highly flexible and does not dock the BP adenosine into the active site^65^. The Prp8 1585-loop, which directly interacts with U2/U6 helix I, Cwc24, and Prp11, and protects the G_+1_ nucleotide at the 5’ss in the B^act^ complex^52^, is likewise flexible at this intermediate step.

**Fig. 4:**
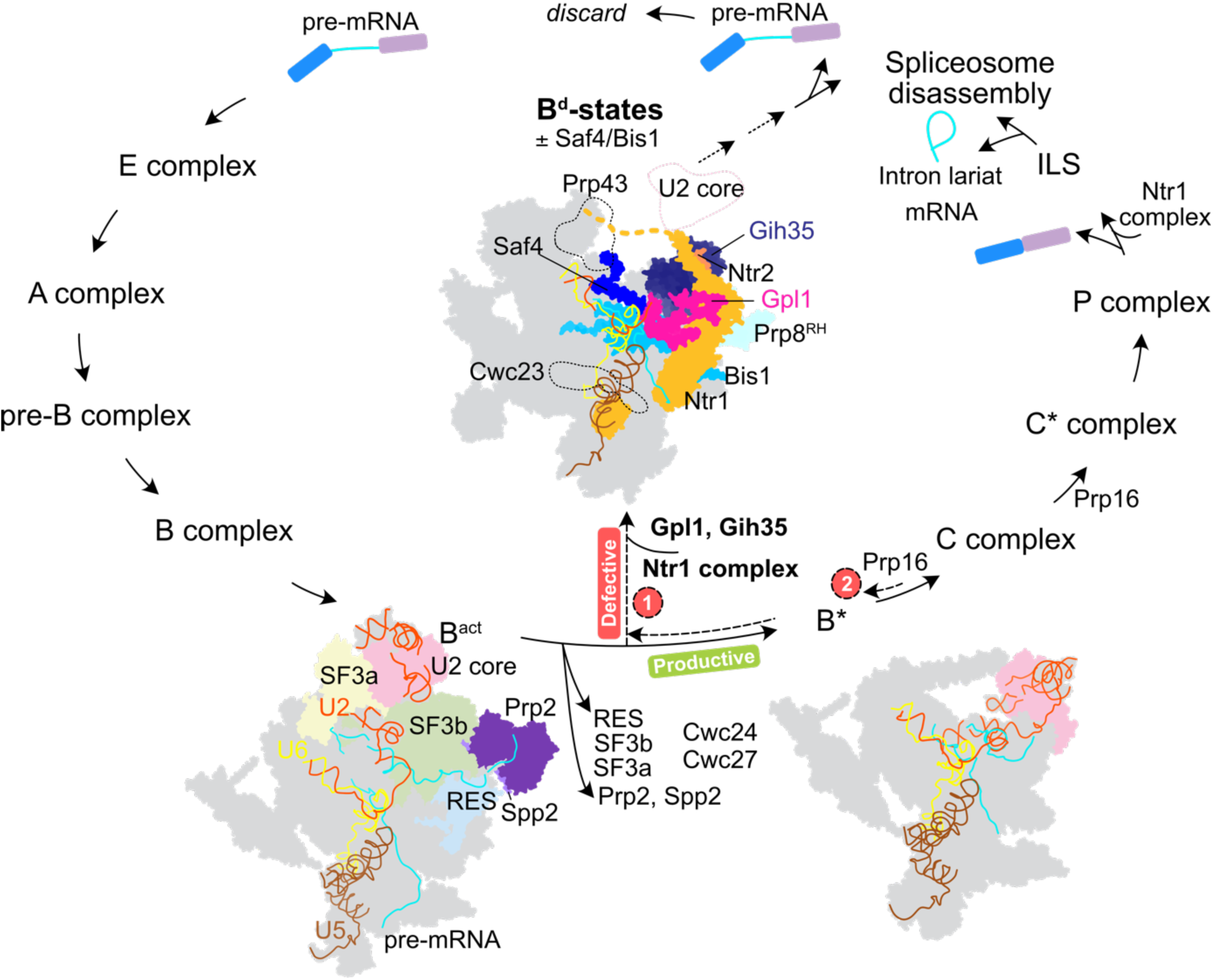
Scheme for Gpl1-mediated discard of the aberrant B^d^ complex. Following the action of the Prp2-Spp2 complex, the RES, SF3a, SF3b complexes and splicing factors Cwc24 and Cwc27 are released. Sensing an aberration, Gpl1 either alone or in complex with Gih35 binds to the active site blocking splicing progression. Alternatively, the spliceosome is targeted for disassembly via Prp16-mediated proofreading before branching, triggering Gpl1-Gih35 binding at the B* state. Concomitantly, the Ntr1 complex is recruited and Saf4 and Bis1 help stabilize the Ntr1 complex in the B^d^ complex. Subsequently, disassembly of the defective spliceosome can be initiated by Prp43. The two discard models are marked by dashed lines.

We propose that changes in the conformation of the active site, as observed in this study, might lead to inefficient binding of Prp11 and Cwc24 to the pre-mRNA in the B^act^ complex. Cwc24 is required for both proper interaction of U5 and U6 with the 5’ss and 5’ss selection and stable association of Prp2 with the spliceosome, and its deletion results in aberrant cleavage at the 5’ss^71,72^. Whether it is the propensity for the altered U5 loop I-5’ exon interaction at positions -5 to -2 of the pre-mRNA (due to the exonic quadruple Adenine motif) that leads to a compromised Cwc24 binding capacity or the other way around, in both cases this over-stabilized ‘improper’ interaction leads to the non-canonical active site conformation as observed in the B^d^ complex. Subsequently, Prp2 and/or Cwf11^70^ mediated remodeling of the spliceosome might be faster, creating a time window for recruitment and binding of Gpl1 to the modified active site. Sensing of the 5’ss aberration latest at this stage involves Gpl1 binding at the active site, either as a pre-formed complex with Gih35 or with Gih35 being recruited sequentially. Gpl1 binding occludes the docking of the branch helix at the active site and thus maintains catalytic dormancy of the spliceosome as found in our B^d^ complex. The 1585-loop becomes ordered due to its interaction with Gpl1. Taken together, binding of Gpl1 blocks the progression of productive splicing at the B^d^ intermediate between the B^act^ and B* states. Alternatively, the spliceosome could reach the B* state transiently and then be targeted by proofreading activity of Prp16. In the B*/C complexes, branching is promoted by engagement of N-termini of step I splicing factors Cwc25, Isy1 and Yju2 with the branch helix and branching occurs spontaneously upon Cwc25 binding^54,64,65^. However, removal of Cwc25 by Prp16 prior to branching reverts the spliceosome back to B* state which is vulnerable to disassembly^73^. The changes in the 5’ss conformation as observed in the B^d^ complex could hinder canonical interactions with the branch helix^54,64^, likely weakening the association of Cwc25 with the spliceosome, delaying branching and therefore promoting Prp16-mediated discard and recruitment of Gih35-Gpl1.

Once Gpl1 is loaded into the active site and folds on the surface of Prp8, Gih35 is positioned right at the beginning of the branch helix. Weak cryo-EM density attributable to two-three nucleotides in the RNA binding channel of Gih35 is observed (**Extended Data Fig. 19a**). Although in the absence of continuous cryo-EM density into the active site of Gih35, the identity of its RNA substrate remains elusive, structural comparisons with *sc*B^act^ and *sc*B* complexes show its close proximity to the branch helix and indicate that pre-mRNA or U2 snRNA are plausible candidates (**Extended Data Fig. 19b-e**). In any case, due to steric exclusion the branch helix must be dissolved in order to allow, or as a consequence of, Gih35 binding.

As a consequence of Gih35-Gpl1 binding, catalysis is blocked and the spliceosome is licensed to be disassembled by the Ntr1 complex. While the binding site for Prp43 on Syf1 is available in all spliceosomes containing the NTC, the position of the Ntr1 complex is incompatible with SF3b in the B^act^ complex, Prp16 in the B* and C complexes, and Prp22 in the C* and P complexes preventing premature disassembly by Prp43^3^. In the *sp*B^d^ complexes, we find that only the Ntr1 CTD is fixed in the *sp*B^d^-I state, while the Ntr1-superhelical domain and Ntr2 are more stabilized in the *sp*B^d^-II complex, and Cwc23 and Prp43 remain flexibly attached throughout.

For the maintenance of splicing fidelity, several RNA helicases proofread and discriminate against suboptimal substrates triggering Prp43-mediated disassembly of defective spliceosomes in an internal splicing quality control mechanism^11,16,31,32^. Structural or compositional changes in defective spliceosomes such as a compromised conformation of the branch helix may additionally (or instead) trigger disassembly via the Ntr1 complex^35^. Therefore, the Ntr1 complex could either be recruited to the *sp*B^d^ complex after proofreading by an RNA helicase (such as Prp16, suggested above), or spontaneously as a consequence of Gpl1 blocking the progression of productive splicing. We find that binding of Saf4 and Bis1 help in stabilization of the Ntr1 complex on the spliceosome (**Extended Data Fig. 7e**). A similar role for ESS2 has been suggested in humans, where its binding to SKIP (homologous to Prp45) helps stabilize the connection from PRP22 to the spliceosome core^45^. However, we find that Bis1 does not cause stabilization of Gih35 on the spliceosome. It is intriguing that two factors usually recruited to the spliceosome at different stages, namely step I factor Yju2 at B* in *S. cerevisiae*^65^ and ESS2 at C* in humans^45^, are recruited together in the *sp*B^d^-II complex. In addition, Gih35-Gpl1 also usually recruited at the C* state in humans^45^ are already recruited in the B^d^ complex. Taken together, our data suggest that the aforementioned proteins are recruited before branching in the B^d^ complex, when an aberration is detected and the spliceosome needs to be discarded.

As mentioned above, Gih35-Gpl1 are found in catalytically active spliceosomes in humans and also in *C. neoformans* where they have been suggested to play a role in maintenance of splicing fidelity by a proofreading mechanism^74^. Studies in *S. pombe* also report that Gih35-Gpl1 are required for proper canonical splicing^48,49^. Together, this suggests that Gih35-Gpl1 might not only be recruited to aberrant or defective spliceosomes undergoing discard via the CNM complex as seen in our purifications but also to productive spliceosomes. Indeed Gih35-Gpl1 were also detected in purifications from *S. pombe* using an affinity-tagged NTC component (Cef1) while components of the CNM complex were not reported^36^. Taken together, these observations indicate that Gih35-Gpl1 might have other functions apart from their central role in the discard of aberrant spliceosome complexes proposed here.

How would the conformation of Gih35-Gpl1 bound to a C* complex compare to the B^d^ complex? To answer this question, we performed structural superpositions of our *sp*B^d^-II complex with the human C* complex. We find that Gpl1 cannot enter the active site in *hs*C* due to steric clashes with Prp8^RH^, PRKRIP1 and FAM32A binding at the position of the 5’ss G_+1_U_+2_ nucleotides in the B^d^ complex (**Extended Data Fig. 20a-b**). While all crosslinks between GPATCH1, DHX35 and CDC5L observed in the *hs*C* are satisfied in the *sp*B^d^ complex, those between GPATCH1 and PRP8^RH^ remain unsatisfied due to substantial differences in location of Prp8^RH^/PRP8^RH^ with respect to Gpl1-Gih35 in the *sp*B^d^ complex (**Extended Data Fig. 20c-f**). Taken together, the interaction between GPATCH1 and DHX35 remains likely conserved from the *sp*B^d^ complex to *hs*C*. However, Gpl1 might not be bound to the active site, being associated only at the periphery, where it could help in tethering Gih35 to the spliceosome. Moreover, Gih35-Gpl1 could function during the transition from C to C* complex where it might associate transiently in a pre-C* complex as proposed before^45^. Further biochemical, structural and functional studies need to be performed to understand the specific role of Gih35-Gpl1 as part of catalytically active spliceosomes.

Our RNA-seq experiments show that *rpl39* is an example of a gene prone to discard via the CNM complex. *Rpl39* encodes the ribosomal protein eL39 important for maturation of the nascent polypeptide exit tunnel^75^. Curiously the first exon of *rpl39* itself encodes a stop codon and therefore does not in principle require proper splicing for the production of a functional transcript. It is important to note that *rpl39* mRNA is not always discarded and majority of the *rpl39* transcripts go through productive splicing. So why is a productive spliceosome sometimes assembled on this substrate or results in a defective spliceosome? We propose that depending on the register of interactions between the 5’ss exon and U5/U6 snRNAs, the erroneous conformation of the active site is an aberrant complex of similar binding energy, which is then discarded via the B^d^-state route. Recent reports show that fine tuning the strength of interactions between the 5’ss and U5 loop I, 5’ss and U6 ACAGA box together determine the efficiency of splicing^76,77^. In *S. pombe*, the m^6^A modification status of the central adenosine of the ACAGA box, which pairs with the +4 position in the intron, changes the strength of the U6-intron interaction^76^. For *rpl39* mRNA, the U_+4_ could form an energetically favorable Watson-Crick base pair with unmethylated U6 AC**A**GA^78^, which together with strong base pairing of U5 loop I-5’ exon could be a detrimental combination, compared to a decreased stability of the U6-intron duplex due to the m^6^A modification. In addition, trans-acting factors that recognize cis-regulatory RNA elements to modulate spliceosome recognition of certain splice sites^79^ might also contribute to context-specific splicing efficiency of *rpl39* mRNA. The propensity of *rpl39* mRNA to be discarded might actually have a regulatory role itself, with changes in its pre-mRNA splicing efficiency modulating the amount of the protein produced. The influence on ribosome biosynthesis by varying the amounts of different ribosomal proteins is a well-known phenomenon^80^.

Taken together, the *sp*B^d^ complexes describe a new route for the recognition and discard of aberrant splicing intermediates. This route is part of a splicing quality control mechanism characterized by a shortcut of the splice cycle involving the Gih35-Gpl1 DEAH-box helicase – G-patch protein pair. As splicing errors are an increasing threat of a growing intron complexity, more strategies and components of splicing quality control mechanisms await to be deciphered.

## Methods

### TAP tagging in *S. pombe*

Using Cre-recombinase-mediated cassette exchange (RMCE)^81^ system, the respective genes were tagged at the C-termini with a loxP/M flanked ura4+ cassette, which was further replaced by a Myc-tag fused to a Tobacco etch virus (TEV) protease cleavage site and a protein A tag in case of Nrl1 and 6x-Flag tag in case of Prp43. The transformants were replica plated on SDC+FOA, SDC-Ura or SDC-Leu plates and selected for those only growing on SDC+FOA plates. Finally, the genomic integration of the TAP-tagged Nrl1 and Prp43 were confirmed by PCR and western blot^40^.

### Split-tag TAP and RNA-IP from *S. pombe*

The split-tagged RNP complex was harvested from ∼10-12 L YEA cultures of the yeast strain grown to OD_600_ of 1.8-2.2. Cell pellets were snap-frozen in liquid nitrogen and subsequently mechanically lysed by a cryogenic cell mill (Retsch MM400). The cell lysate was resuspended in lysis buffer containing 20 mM HEPES pH 7.5, 100 mM NaCl, 1.5 mM MgCl_2_ supplemented with 0.05 % NP-40 (Sigma-Aldrich), 1 mM DTT, 1 mM phenylmethylsulfonyl fluoride, protease inhibitor cocktail (EDTA-free, Roche) and RiboLock RNase inhibitor (Thermo Fischer Scientific). The cell lysate was cleared by centrifugation and the supernatant was loaded onto IgG Sepharose 6 Fast Flow beads (Cytiva) for overnight (ON) incubation at 4 °C. Non-specifically bound proteins were removed by washing with lysis buffer supplemented with 0.01 % NP-40, followed by TEV cleavage for 5 h at 4 °C. For the second affinity purification, the eluate from IgG beads was loaded onto Flag beads (Anti-Flag M2 Affinity Gel, Sigma-Aldrich) for ON incubation at 4 °C. The beads were washed and eluted with buffer containing 3X Flag peptide at a final concentration of 0.4 mg/mL. The elution buffer for cryo-EM sample preparation contained 20 mM HEPES pH 7.5, 100 mM NaCl, 1.5 mM MgCl_2_, 0.01 % NP-40 and 1 mM TCEP. In RNA-IP experiments, the RNA was isolated from the Flag-eluate, using RNA Clean and Concentrator Kit (Zymo Research), treated with 20 U Turbo DNase (Thermo Fischer) at 37C for 30 minutes. RNA Clean and Concentration Kit was used once again to remove DNase. RNA sequencing libraries were prepared using NEBNext® Ultra II Directional RNA Library Prep Kit for Illumina® (NEB) following the manufacturer’s instructions.

### Electron microscopy sample preparation and data acquisition

Negative staining was performed as a quality control before cryo-EM grid preparation and after crosslinking with BS3 for mass spectrometric analysis to control for sample aggregation. For cryo-EM grid preparation, a 5 μL aliquot of the sample was applied to Quantifoil R 2/1 grids coated with a thin layer of homemade carbon film, glow discharged for 45 sec using PELCO easiGlow. After ∼8-10 min incubation of the sample on grids, blotting was performed for 5 sec using a blot force of 0 at 100% humidity using Vitrobot Mark IV (FEI) operated at 4 °C and the grids were immediately plunge-frozen in liquid ethane cooled with liquid nitrogen. Cryo-electron microscopy data were acquired at the cryo-electron microscopy platform of the European Molecular Biology Laboratory (EMBL) in Heidelberg on 300 kV FEI Titan Krios electron microscope equipped with a K3 detector (Gatan) in counting mode at 0.822 Å/pixel with 1.235 e^-^/pixel/frame over 40 frames and 1.491 sec total exposure. In total, 13,096 images were collected.

### Image processing

The images were processed using Relion 3.1 software package^82^. Movie stacks were motion corrected using MotionCor2 with 5x5 as the number of patches^83^ and estimation of contrast transfer function was performed with Gctf v1.06^84^ on the motion corrected micrographs. After manual curation of particles picked with Warp v1.0.9^85^ on 10 micrographs, 929,930 particles were automatically picked from all micrographs. Motion correction and CTF estimation was simultaneously performed in cryoSPARC v3.1^86,87^. Using particle coordinates from Warp, particles were extracted in cryoSPARC and subjected to two-dimensional classification and initial *ab initio* 3D construction, heterogenous refinement which was subsequently low-pass filtered to 60 Å and used as the initial model for 3D classification of all particles (929,930) in Relion. Two best classes were merged (256,572 particles) and subjected to 3D auto-refinement. These particles were then imported in cisTEM v1.0.0^88^ where they were first subjected to 3D auto-refinement (6 cycles) and subsequently to focused classification using manual refinement (3 classes, 40 cycles) by applying a soft circular mask around Gih35. This yielded a single class average with cryo-EM density for Gih35 (153,834 particles, 59.2%) which was further subjected to another round of focused classification with a soft mask around the Ntr1 complex (3 classes, 40 cycles). Two good classes were obtained of which one showed decreased signal for Ntr1 (*sp*B^d^-I; 40.1%; 61,423 particles) compared to the other (*sp*B^d^-II; 47.1%; 72,631 particles). These particle stacks were then imported into Relion and subjected to 3D auto-refinement, CTF-refinement and Bayesian polishing to give final reconstructions with overall resolutions of 3.2 Å (*sp*B^d^-I) and 3.1 Å (*sp*B^d^-II). The cryo-EM maps were sharpened in Phenix (version 1.19.2)^89^ using the Autosharpen map tool and local resolution estimation in was performed in Relion.

For the *sp*B^d^-II state, further rounds of 3D classification without image alignment with separate masks around the Prp8^RH^, Ntr1 complex and Cwf11 were performed, which invariably led to better local reconstructions in the flexible regions. After 3D-autorefinement of the subset of particles for Prp8^RH^ (20,173), Ntr1 complex (21,292) and Cwf11 (9,361), the cryo-EM maps were low-pass filtered to 7 Å, 8 Å, and 10 Å, respectively, to obtain more continuous cryo-EM density in these regions. For Gih35, another round of 3D auto-refinement was performed in Relion, which served as a basis for multibody refinement. For this, the bulk of the spliceosome was treated as one component and the region encompassing Prp8^En^, Prp8^RH^ and Gih35 was treated as the second component, moving independently of each other. The final cryo-EM densities after multi-body refinement were subjected to post-processing including automatic B-factor sharpening and local resolution estimation in Relion. The particle stack for *sp*B^d^-II was imported in cryoSPARC where local refinement was performed with a mask around the NTC leading to an improved resolution of 4.3 Å in this region. For Ntr1 CTD, focused 3D classification (in cryoSPARC) with masks around Snu114 and Ntr1 CTD followed by local refinement led to a local resolution of ∼6-8 Å in the *sp*B^d^-I (6,500 particles) and *sp*B^d^-II (8,560 particles) states. A detailed summary of the image processing workflow is illustrated in **Extended Data Fig. 2**.

### Structural modelling, refinement and analysis

We used a combination of *de novo* model building, rigid body docking of known structures, homology models obtained using Swissmodel^90^, and models deposited in the AlphaFold protein structure database^91^ for different components of the *sp*B^d^ complexes depending on the local resolution. Identification and docking of different components of the *sp*B^d^ complexes was facilitated by the *sp*ILS complex (PDB ID: 3JB9)^36^, *sc*ILS complex (PDB ID: 5Y88)^47^, *sc*C complex (PDB ID: 5LJ3)^54^, *sc*B* complex (PDB ID: 6J6Q)^65^ and *hs*C* complex (PDB ID: 5MQF)^92^. Identification of Saf4 and Bis1 in the *sp*B^d^-II cryo-EM map was done using ModelAngelo^93^. The structures of the individual components were first fitted into cryo-EM density using UCSF Chimera v1.12^94^ and then manually adjusted in Coot v0.8.9.3-pre^95^.

Gpl1 was built *de novo* in the cryo-EM map. The Gpl1 fragment contacting Gih35 suffers from limited local resolution. Here, high resolution structures of other DEAH-helicase – G-patch protein pairs (PDB ID: 6HS6^62^, 7DCP^53^), together with Clustal Omega^96^ based sequence alignments and secondary structure prediction using PSIPRED^97^, aided in model building and identification of the sequence register. For Gih35, the AlphaFold model was rigid body docked in the cryo-EM map in Chimera and then individual domains were manually adjusted to best fit the cryo-EM density in Coot. All components were first built in the *sp*B^d^-II cryo-EM map and were then adjusted for differences in the *sp*B^d^-I cryo-EM map in Coot. The pre-mRNA sequence was built according to the consensus sequence logo from CNM-sensitive genes between positions -5 to +6 and uracil or adenine bases were placed where a pyrimidine or purine satisfied the cryo-EM map better.

While the models of individual components of the *sp*B^d^ complex were built in and refined against focused maps (only for *sp*B^d^-II), the final model was real space refined against the overall map in Phenix v1.19.2^89^ with secondary structure and geometry restraints. The model qualities were assessed using MolProbity^98^ within Phenix. A detailed summary of modelling building for *sp*B^d^-I and *sp*B^d^-II complexes are presented in **Extended Data Table 2.**

### Mass spectrometry analysis of the *sp*B^d^ complex

Approximately 5 pmol of the *sp*B^d^ complex was purified from ∼24 L YEA culture as described above, crosslinked with 1 mM BS3 (Thermo Scientific) for 30 min at 30 °C, subsequently quenched with 50 mM ammonium bicarbonate and pelleted by ultracentrifugation. The pelleted complex was solubilized in 50 mM ammonium bicarbonate (pH 8.0) supplemented with 4M Urea, reduced with dithiothreitol and alkylated with iodoacetamide. After dilution to 1 M urea with 50 mM ammonium bicarbonate, crosslinked complexes were digested with trypsin (Promega) in a 1:20 enzyme-to-protein ratio (w/w) at 37 °C overnight. Peptides were reverse-phase extracted using SepPak Vac tC18 1cc/50mg (Waters) and eluted with 50% acetonitrile (ACN) / 0.1 % TFA. The eluate was lyophilized. Dried peptides were dissolved in 40 µl 2% ACN / 20 mM ammonium hydroxide and reverse-phase fractionated at basic pH using a Vanquish HPLC system (Thermo Scientific) with an xBridge C18 3.5µm 1x150mm column (Waters) applying a 4-36% ACN gradient over 45 min at a flow rate of 60 µl/min. One-minute fractions of 60 µl were collected, pooled in a step of 13 minutes (resulting in 13 pooled fractions total), vacuum dried and dissolved in 5% ACN / 0.1% TFA for subsequent uHPLC-ESI-MS/MS analysis.

Fractionated peptides were measured in triplicate on an Orbitrap Exploris 480 (Thermo Scientific). The mass spectrometer was coupled to a Dionex UltiMate 3000 uHPLC system (Thermo Scientific) with a custom 35 cm C18 column (75 µm inner diameter packed with ReproSil-Pur 120 C18-AQ beads, 3 µm pore size, Dr. Maisch GmbH). MS1 and MS2 resolution were set to 120,000 and 30,000, respectively. Only precursors with a charge state of 3-8 were selected for MS2. Protein composition was determined by MaxQuant v.1.6.17.0. The entire S. pombe proteome or a set of fourty-five abundant spliceosomal proteins identified by MaxQuant were used as a database to search for crosslinked peptides. Protein-protein crosslinks were searched using pLink v.2.3.9 (pfind.org/software/pLink) according to the recommendations of the developer^99^. A list of the top 200 proteins identified in the sample, ranked according to label-free quantitation (LFQ) intensity as well as inter-protein and intra-protein crosslinks of these proteins are provided in **Supplementary Table 1**.

### RNA-seq data analysis and assessing intron retention in the *ctr1**Δ*** mutant and in RNA-IP experiments

Paired-end RNA-seq reads were aligned to the *S. pombe* reference genome (ASM294v2) using HISAT2^100^ allowing a maximum intron length of 2,000 bps. Alignment files were sorted and indexed using SAMtools^101^. Un-normalized bigwig files were generated with bamCoverage from deepTools^102^ with 1 bp resolution. To avoid any bias from rRNAs transcribed on chromosome III, raw signals were normalized by the sum of the coverage of chromosome I and II using in-house bash and R scripts. Intron annotation was adopted from^103^ and only introns longer than 10 bps were used. For RNA expression analysis, introns with normalized scores ≥30 in both biological replicates were included in downstream analyses. *Sensitivity score* in the *ctr1Δ* mutant was calculated by subtracting the normalized RNA-seq signal over the introns of the WT strain from the *ctr1Δ* strain, divided by the averaged signal of the neighboring exons. The 5’ss and 3’ss with an upstream and downstream flanking regions of 9 and 10 bases, respectively, were inferred from the reference genome using the *seqinr*^104^ and *bedr* R packages^105^. The occurrences of 5’ss and 3’ss sequences were then scaled to be proportional to the average signal of the upstream exon, measured in the wildtype strain. Introns were considered as *Sensitive* with at least 20% increase in abundance in the *ctr1Δ* strains compared to wildtype. The probability weight matrices of the 5’ss and 3’ss of the sensitive introns were plotted as motif logos using the gglogo R package. Lists of all CNM-sensitive and CNM-insensitive sites are provided in **Supplementary Table 2**. Absolute intron coverage in RNA-IP experiments were determined by the average sequence coverage at the 5’ end (5% of the total intron length) of all annotated introns. Intron enrichments were calculated in *sp*B^d^ RNA-IP sample over no-tag RNA-IP control.

### Cloning, protein expression and *in vitro* complex assembly

The DNA sequence encoding Prp43 (residues 1-735), Ntr1 (residues 107-170) and Gpl1 (residues 109-181) were cloned and ligated into a modified pET24d vector containing an N-terminal His-Thioredoxin tag upstream of a Tobacco-Etch-Virus cleavage site before the corresponding protein. The plasmids were transformed in BL21(DE3) Rosetta electrocompetent cells, grown at 37 °C up to an OD of 1.4-1.6 in auto-induction media^106^ and subsequently expressed at 20 °C for ∼16 hrs. Cells were lysed in buffer containing 20 mM Tris pH 7.5, 500 mM NaCl, 1M Urea, 20 mM imidazole, and 2 mM β-mercaptoethanol. The proteins were purified over 2 ml His-Trap FF column (GE Healthcare) with elution buffer containing 250 mM imidazole. Next, overnight tag-cleavage using TEV protease and simultaneous dialysis of the proteins into buffer containing 20 mM HEPES pH 7.5, 150 mM NaCl, 2 mM β-mercaptoethanol was carried out. The proteins were further purified over His-Trap FF columns to remove uncleaved proteins and fusion tags from the cleaved proteins and the flow through samples were collected. Finally, the proteins were polished using gel filtration (Superdex 200 16/60) columns (GE Healthcare) equilibrated with SEC buffer containing 20 mM HEPES pH 7.5, 150 mM NaCl and 1 mM DTT. For monitoring complex formation, Ntr1/Gpl1 co-factors were added in 2-fold molar excess over Prp43, incubated at 4 °C for 30 min and purified using gel filtration (Superdex 200 Increase 10/300 GL) columns (GE Healthcare).

### Yeast two hybrid assays

The respective sequences of full-length proteins were cloned into low copy plasmids-pG4BDN22 (Gpl1, Ntr1) and pG4ADAHAN111 (Gih35, Prp43) which harbor either the DNA binding domain or activation domain at the N-termini, respectively. The interaction pairs were analyzed by co-transformation into PJ69-4A strain. After a 10-fold serial dilution, colonies were spotted on SDC (SDC-Leu-Trp), SDC-His (SDC-Leu-Trp-His) and SDC-Ade (SDC-Leu-Trp-Ade) plates, incubated at 30 °C and analyzed after 3 days. The strength of the interaction was assessed by growth achieved on SDC-His and SDC-Ade as weak and strong, respectively.

## Data availability

The structures of *sp*B^d^-I and -II states are deposited in the PDB under accession codes 9ESH and 9ESI and cryo-EM maps are available at the EMDB under the accession codes EMD-19941 and EMD-19942, respectively. The RNA-seq data have been deposited to NCBI GEO under the reference number: GSE235589 with the following reviewer access token: mroximscttixpat.

## Code availability

All in-house R and bash scripts are available on the following GitHub repository: https://github.com/ahorvath/Soni-et-al.-2023.git

## Acknowledgements

We are thankful to EMBL Heidelberg for seamless cryo-EM data collection via the iNEXT Discovery program, to HDCryoNet for access to our inhouse cryo-EM setup for screening, and Nikolay Dobrev and Melanie McDowell for their inputs on the project. We are grateful to Janosch Hennig for supporting completion of final experiments by K.S. in his lab. We acknowledge the data storage service SDS@hd and bwHPC supported by the Ministry of Science, Research and the Arts Baden-Württemberg (MWK) and the German Research Foundation (DFG) through grants INST 35/1314-1 FUGG, INST 35/1503-1 FUGG and INST 35/1597-1 FUGG. This work was supported by the Australian Research Council’s Discovery Projects (projects DP190100423, DP240102611) to T.F and A.H.; the DFG through the Leibniz program (SI 586/6-1), and TRR 319 (Project-ID 439669440, TP B03) to I.S.

## Author contributions

K.S. purified the spB^d^ complex, prepared cryo-EM samples, collected, processed EM data and performed *in vitro* protein expression and complex assembly. O.D. performed proteome analysis and crosslinking mass spectrometry. O.D. and H.U. interpreted the mass spectrometry results. M.S. performed yeast-two-hybrid experiments. S.T. prepared samples for RNA-seq. A.H. and T.F. performed the RNA-sequencing analysis. D.F. performed negative staining and helped with cryo-EM grid screening. K.S. and K.W. built the structural models. K.S., K.W., T.F. and I.S. interpreted the data. K.S., K.W. and I.S. wrote the manuscript. K.S., T.F. and I.S. planned the study and designed the experiments. All authors contributed to the final version of this manuscript.

## Additional Information

Supplementary Information and Source Data are available for this paper.

## References

1 Scotti, M. M. & Swanson, M. S. RNA mis-splicing in disease. Nat Rev Genet 17, 19–32 (2016). 10.1038/nrg.2015.3

2 Will, C. L. & Luhrmann, R. Spliceosome structure and function. Cold Spring Harb Perspect Biol 3 (2011). 10.1101/cshperspect.a003707

3 Wilkinson, M. E., Charenton, C. & Nagai, K. RNA Splicing by the Spliceosome. Annu Rev Biochem 89, 359–388 (2020). 10.1146/annurev-biochem-091719-064225

4 Brody, E. & Abelson, J. The "spliceosome": yeast pre-messenger RNA associates with a 40S complex in a splicing-dependent reaction. Science 228, 963–967 (1985). 10.1126/science.3890181

5 Frendewey, D. & Keller, W. Stepwise assembly of a pre-mRNA splicing complex requires U-snRNPs and specific intron sequences. Cell 42, 355–367 (1985). 10.1016/s0092-8674(85)80131-8

6 Grabowski, P. J., Seiler, S. R. & Sharp, P. A. A multicomponent complex is involved in the splicing of messenger RNA precursors. Cell 42, 345–353 (1985). 10.1016/s0092-8674(85)80130-6

7 Shi, Y. Mechanistic insights into precursor messenger RNA splicing by the spliceosome. Nat Rev Mol Cell Biol 18, 655–670 (2017). 10.1038/nrm.2017.86

8 Cordin, O. & Beggs, J. D. RNA helicases in splicing. RNA Biol 10, 83–95 (2013). 10.4161/rna.22547

9 Cordin, O., Hahn, D. & Beggs, J. D. Structure, function and regulation of spliceosomal RNA helicases. Curr Opin Cell Biol 24, 431–438 (2012). 10.1016/j.ceb.2012.03.004

10 Xu, Y. Z. & Query, C. C. Competition between the ATPase Prp5 and branch region-U2 snRNA pairing modulates the fidelity of spliceosome assembly. Mol Cell 28, 838–849 (2007). 10.1016/j.molcel.2007.09.022

11 Chen, H. C., Tseng, C. K., Tsai, R. T., Chung, C. S. & Cheng, S. C. Link of NTR-mediated spliceosome disassembly with DEAH-box ATPases Prp2, Prp16, and Prp22. Mol Cell Biol 33, 514–525 (2013). 10.1128/MCB.01093-12

12 Wlodaver, A. M. & Staley, J. P. The DExD/H-box ATPase Prp2p destabilizes and proofreads the catalytic RNA core of the spliceosome. RNA 20, 282–294 (2014). 10.1261/rna.042598.113

13 Burgess, S. M. & Guthrie, C. A mechanism to enhance mRNA splicing fidelity: the RNA-dependent ATPase Prp16 governs usage of a discard pathway for aberrant lariat intermediates. Cell 73, 1377–1391 (1993). 10.1016/0092-8674(93)90363-u

14 Koodathingal, P., Novak, T., Piccirilli, J. A. & Staley, J. P. The DEAH box ATPases Prp16 and Prp43 cooperate to proofread 5’ splice site cleavage during pre-mRNA splicing. Mol Cell 39, 385–395 (2010). 10.1016/j.molcel.2010.07.014

15 Mayas, R. M., Maita, H. & Staley, J. P. Exon ligation is proofread by the DExD/H-box ATPase Prp22p. Nat Struct Mol Biol 13, 482–490 (2006). 10.1038/nsmb1093

16 Egecioglu, D. E. & Chanfreau, G. Proofreading and spellchecking: a two-tier strategy for pre-mRNA splicing quality control. RNA 17, 383–389 (2011). 10.1261/rna.2454711

17 Fourmann, J. B. et al. Dissection of the factor requirements for spliceosome disassembly and the elucidation of its dissociation products using a purified splicing system. Genes Dev 27, 413–428 (2013). 10.1101/gad.207779.112

18 Tsai, R. T. et al. Spliceosome disassembly catalyzed by Prp43 and its associated components Ntr1 and Ntr2. Genes Dev 19, 2991–3003 (2005). 10.1101/gad.1377405

19 Martin, A., Schneider, S. & Schwer, B. Prp43 is an essential RNA-dependent ATPase required for release of lariat-intron from the spliceosome. J Biol Chem 277, 17743–17750 (2002). 10.1074/jbc.M200762200

20 Arenas, J. E. & Abelson, J. N. Prp43: An RNA helicase-like factor involved in spliceosome disassembly. Proc Natl Acad Sci U S A 94, 11798–11802 (1997). 10.1073/pnas.94.22.11798

21 Boon, K. L. et al. Yeast ntr1/spp382 mediates prp43 function in postspliceosomes. Mol Cell Biol 26, 6016–6023 (2006). 10.1128/MCB.02347-05

22 Warkocki, Z. et al. The G-patch protein Spp2 couples the spliceosome-stimulated ATPase activity of the DEAH-box protein Prp2 to catalytic activation of the spliceosome. Genes Dev 29, 94–107 (2015). 10.1101/gad.253070.114

23 Roy, J., Kim, K., Maddock, J. R., Anthony, J. G. & Woolford, J. L., Jr. The final stages of spliceosome maturation require Spp2p that can interact with the DEAH box protein Prp2p and promote step 1 of splicing. RNA 1, 375–390 (1995).

24 Aravind, L. & Koonin, E. V. G-patch: a new conserved domain in eukaryotic RNA-processing proteins and type D retroviral polyproteins. Trends Biochem Sci 24, 342–344 (1999). 10.1016/s0968-0004(99)01437-1

25 Bohnsack, K. E., Ficner, R., Bohnsack, M. T. & Jonas, S. Regulation of DEAH-box RNA helicases by G-patch proteins. Biol Chem 402, 561–579 (2021). 10.1515/hsz-2020-0338

26 Lardelli, R. M., Thompson, J. X., Yates, J. R., 3rd & Stevens, S. W. Release of SF3 from the intron branchpoint activates the first step of pre-mRNA splicing. RNA 16, 516–528 (2010). 10.1261/rna.2030510

27 Bao, P., Hobartner, C., Hartmuth, K. & Luhrmann, R. Yeast Prp2 liberates the 5’ splice site and the branch site adenosine for catalysis of pre-mRNA splicing. RNA 23, 1770–1779 (2017). 10.1261/rna.063115.117

28 Ohrt, T. et al. Prp2-mediated protein rearrangements at the catalytic core of the spliceosome as revealed by dcFCCS. RNA 18, 1244–1256 (2012). 10.1261/rna.033316.112

29 Fourmann, J. B. et al. The target of the DEAH-box NTP triphosphatase Prp43 in Saccharomyces cerevisiae spliceosomes is the U2 snRNP-intron interaction. Elife 5 (2016). 10.7554/eLife.15564

30 Pandit, S., Lynn, B. & Rymond, B. C. Inhibition of a spliceosome turnover pathway suppresses splicing defects. Proc Natl Acad Sci U S A 103, 13700–13705 (2006). 10.1073/pnas.0603188103

31 Koodathingal, P. & Staley, J. P. Splicing fidelity: DEAD/H-box ATPases as molecular clocks. RNA Biol 10, 1073–1079 (2013). 10.4161/rna.25245

32 Semlow, D. R. & Staley, J. P. Staying on message: ensuring fidelity in pre-mRNA splicing. Trends Biochem Sci 37, 263–273 (2012). 10.1016/j.tibs.2012.04.001

33 Mayas, R. M., Maita, H., Semlow, D. R. & Staley, J. P. Spliceosome discards intermediates via the DEAH box ATPase Prp43p. Proc Natl Acad Sci U S A 107, 10020–10025 (2010). 10.1073/pnas.0906022107

34 Tsai, R. T. et al. Dynamic interactions of Ntr1-Ntr2 with Prp43 and with U5 govern the recruitment of Prp43 to mediate spliceosome disassembly. Mol Cell Biol 27, 8027–8037 (2007). 10.1128/MCB.01213-07

35 Fourmann, J. B., Tauchert, M. J., Ficner, R., Fabrizio, P. & Luhrmann, R. Regulation of Prp43-mediated disassembly of spliceosomes by its cofactors Ntr1 and Ntr2. Nucleic Acids Res 45, 4068–4080 (2017). 10.1093/nar/gkw1225

36 Yan, C. et al. Structure of a yeast spliceosome at 3.6-angstrom resolution. Science 349, 1182–1191 (2015). 10.1126/science.aac7629

37 Lee, N. N. et al. Mtr4-like protein coordinates nuclear RNA processing for heterochromatin assembly and for telomere maintenance. Cell 155, 1061–1074 (2013). 10.1016/j.cell.2013.10.027

38 Aronica, L. et al. The spliceosome-associated protein Nrl1 suppresses homologous recombination-dependent R-loop formation in fission yeast. Nucleic Acids Res 44, 1703–1717 (2016). 10.1093/nar/gkv1473

39 Zhou, Y. et al. The fission yeast MTREC complex targets CUTs and unspliced pre-mRNAs to the nuclear exosome. Nat Commun 6, 7050 (2015). 10.1038/ncomms8050

40 Zhu, J. Identification and characterization of the fission yeast exosome targeting complexes: the MTREC and CNM complexes. Doctoral dissertation, Ruperto-Carola University of Heidelberg, Germany (2017).

41 Cipakova, I. et al. Identification of proteins associated with splicing factors Ntr1, Ntr2, Brr2 and Gpl1 in the fission yeast Schizosaccharomyces pombe. Cell Cycle 18, 1532–1536 (2019). 10.1080/15384101.2019.1632126

42 Ren, L. et al. Systematic two-hybrid and comparative proteomic analyses reveal novel yeast pre-mRNA splicing factors connected to Prp19. PLoS One 6, e16719 (2011). 10.1371/journal.pone.0016719

43 Yan, C., Wan, R. & Shi, Y. Molecular Mechanisms of pre-mRNA Splicing through Structural Biology of the Spliceosome. Cold Spring Harb Perspect Biol 11 (2019). 10.1101/cshperspect.a032409

44 De, I. et al. The RNA helicase Aquarius exhibits structural adaptations mediating its recruitment to spliceosomes. Nat Struct Mol Biol 22, 138–144 (2015). 10.1038/nsmb.2951

45 Dybkov, O. et al. Regulation of 3’ splice site selection after step 1 of splicing by spliceosomal C* proteins. Sci Adv 9, eadf1785 (2023). 10.1126/sciadv.adf1785

46 Zhan, X., Yan, C., Zhang, X., Lei, J. & Shi, Y. Structure of a human catalytic step I spliceosome. Science 359, 537–545 (2018). 10.1126/science.aar6401

47 Wan, R., Yan, C., Bai, R., Lei, J. & Shi, Y. Structure of an Intron Lariat Spliceosome from Saccharomyces cerevisiae. Cell 171, 120–132 e112 (2017). 10.1016/j.cell.2017.08.029

48 Selicky, T. et al. Defining the Functional Interactome of Spliceosome-Associated G-Patch Protein Gpl1 in the Fission Yeast Schizosaccharomyces pombe. Int J Mol Sci 23 (2022). 10.3390/ijms232112800

49 Larson, A., Fair, B. J. & Pleiss, J. A. Interconnections Between RNA-Processing Pathways Revealed by a Sequencing-Based Genetic Screen for Pre-mRNA Splicing Mutants in Fission Yeast. G3 (Bethesda) 6, 1513–1523 (2016). 10.1534/g3.116.027508

50 Plaschka, C., Newman, A. J. & Nagai, K. Structural Basis of Nuclear pre-mRNA Splicing: Lessons from Yeast. Cold Spring Harb Perspect Biol 11 (2019). 10.1101/cshperspect.a032391

51 Rauhut, R. et al. Molecular architecture of the Saccharomyces cerevisiae activated spliceosome. Science 353, 1399–1405 (2016). 10.1126/science.aag1906

52 Yan, C., Wan, R., Bai, R., Huang, G. & Shi, Y. Structure of a yeast activated spliceosome at 3.5 A resolution. Science 353, 904–911 (2016). 10.1126/science.aag0291

53 Bai, R. et al. Mechanism of spliceosome remodeling by the ATPase/helicase Prp2 and its coactivator Spp2. Science 371 (2021). 10.1126/science.abe8863

54 Galej, W. P. et al. Cryo-EM structure of the spliceosome immediately after branching. Nature 537, 197–201 (2016). 10.1038/nature19316

55 Yan, C., Wan, R., Bai, R., Huang, G. & Shi, Y. Structure of a yeast step II catalytically activated spliceosome. Science 355, 149–155 (2017). 10.1126/science.aak9979

56 Bai, R., Yan, C., Wan, R., Lei, J. & Shi, Y. Structure of the Post-catalytic Spliceosome from Saccharomyces cerevisiae. Cell 171, 1589–1598 e1588 (2017). 10.1016/j.cell.2017.10.038

57 Fica, S. M. et al. Structure of a spliceosome remodelled for exon ligation. Nature 542, 377–380 (2017). 10.1038/nature21078

58 Liu, S. et al. Structure of the yeast spliceosomal postcatalytic P complex. Science 358, 1278–1283 (2017). 10.1126/science.aar3462

59 Wilkinson, M. E. et al. Postcatalytic spliceosome structure reveals mechanism of 3’-splice site selection. Science 358, 1283–1288 (2017). 10.1126/science.aar3729

60 Fica, S. M., Oubridge, C., Wilkinson, M. E., Newman, A. J. & Nagai, K. A human postcatalytic spliceosome structure reveals essential roles of metazoan factors for exon ligation. Science 363, 710–714 (2019). 10.1126/science.aaw5569

61 Zhang, X. et al. Structures of the human spliceosomes before and after release of the ligated exon. Cell Res 29, 274–285 (2019). 10.1038/s41422-019-0143-x

62 Studer, M. K., Ivanovic, L., Weber, M. E., Marti, S. & Jonas, S. Structural basis for DEAH-helicase activation by G-patch proteins. Proc Natl Acad Sci U S A 117, 7159–7170 (2020). 10.1073/pnas.1913880117

63 Hamann, F. et al. Structural analysis of the intrinsically disordered splicing factor Spp2 and its binding to the DEAH-box ATPase Prp2. Proc Natl Acad Sci U S A 117, 2948–2956 (2020). 10.1073/pnas.1907960117

64 Wilkinson, M. E., Fica, S. M., Galej, W. P. & Nagai, K. Structural basis for conformational equilibrium of the catalytic spliceosome. Mol Cell 81, 1439–1452 e1439 (2021). 10.1016/j.molcel.2021.02.021

65 Wan, R., Bai, R., Yan, C., Lei, J. & Shi, Y. Structures of the Catalytically Activated Yeast Spliceosome Reveal the Mechanism of Branching. Cell 177, 339–351 e313 (2019). 10.1016/j.cell.2019.02.006

66 Newman, A. J. & Norman, C. U5 snRNA interacts with exon sequences at 5’ and 3’ splice sites. Cell 68, 743–754 (1992). 10.1016/0092-8674(92)90149-7

67 Sontheimer, E. J. & Steitz, J. A. The U5 and U6 small nuclear RNAs as active site components of the spliceosome. Science 262, 1989–1996 (1993). 10.1126/science.8266094

68 Kandels-Lewis, S. & Seraphin, B. Involvement of U6 snRNA in 5’ splice site selection. Science 262, 2035–2039 (1993). 10.1126/science.8266100

69 Lesser, C. F. & Guthrie, C. Mutations in U6 snRNA that alter splice site specificity: implications for the active site. Science 262, 1982–1988 (1993). 10.1126/science.8266093

70 Schmitzova, J., Cretu, C., Dienemann, C., Urlaub, H. & Pena, V. Structural basis of catalytic activation in human splicing. Nature (2023). 10.1038/s41586-023-06049-w

71 Wu, N. Y. & Cheng, S. C. Functional analysis of Cwc24 ZF-domain in 5’ splice site selection. Nucleic Acids Res 47, 10327–10339 (2019). 10.1093/nar/gkz733

72 Wu, N. Y., Chung, C. S. & Cheng, S. C. Role of Cwc24 in the First Catalytic Step of Splicing and Fidelity of 5’ Splice Site Selection. Mol Cell Biol 37 (2017). 10.1128/MCB.00580-16

73 Chung, C. S., Wai, H. L., Kao, C. Y. & Cheng, S. C. An ATP-independent role for Prp16 in promoting aberrant splicing. Nucleic Acids Res 51, 10815–10828 (2023). 10.1093/nar/gkad861

74 Sales-Lee, J. et al. Coupling of spliceosome complexity to intron diversity. Curr Biol 31, 4898–4910 e4894 (2021). 10.1016/j.cub.2021.09.004

75 Micic, J. et al. Ribosomal protein eL39 is important for maturation of the nascent polypeptide exit tunnel and proper protein folding during translation. Nucleic Acids Res 50, 6453–6473 (2022). 10.1093/nar/gkac366

76 Ishigami, Y., Ohira, T., Isokawa, Y., Suzuki, Y. & Suzuki, T. A single m(6)A modification in U6 snRNA diversifies exon sequence at the 5’ splice site. Nat Commun 12, 3244 (2021). 10.1038/s41467-021-23457-6

77 Parker, M. T. et al. m(6)A modification of U6 snRNA modulates usage of two major classes of pre-mRNA 5’ splice site. Elife 11 (2022). 10.7554/eLife.78808

78 Roost, C. et al. Structure and thermodynamics of N6-methyladenosine in RNA: a spring-loaded base modification. J Am Chem Soc 137, 2107–2115 (2015). 10.1021/ja513080v

79 Coltri, P. P., Dos Santos, M. G. P. & da Silva, G. H. G. Splicing and cancer: Challenges and opportunities. Wiley Interdiscip Rev RNA 10, e1527 (2019). 10.1002/wrna.1527

80 Petibon, C., Malik Ghulam, M., Catala, M. & Abou Elela, S. Regulation of ribosomal protein genes: An ordered anarchy. Wiley Interdiscip Rev RNA 12, e1632 (2021). 10.1002/wrna.1632

81 Watson, A. T., Garcia, V., Bone, N., Carr, A. M. & Armstrong, J. Gene tagging and gene replacement using recombinase-mediated cassette exchange in Schizosaccharomyces pombe. Gene 407, 63–74 (2008). 10.1016/j.gene.2007.09.024

82 Zivanov, J. et al. New tools for automated high-resolution cryo-EM structure determination in RELION-3. Elife 7 (2018). 10.7554/eLife.42166

83 Zheng, S. Q. et al. MotionCor2: anisotropic correction of beam-induced motion for improved cryo-electron microscopy. Nat Methods 14, 331–332 (2017). 10.1038/nmeth.4193

84 Zhang, K. Gctf: Real-time CTF determination and correction. J Struct Biol 193, 1–12 (2016). 10.1016/j.jsb.2015.11.003

85 Tegunov, D. & Cramer, P. Real-time cryo-electron microscopy data preprocessing with Warp. Nat Methods 16, 1146–1152 (2019). 10.1038/s41592-019-0580-y

86 Punjani, A., Rubinstein, J. L., Fleet, D. J. & Brubaker, M. A. cryoSPARC: algorithms for rapid unsupervised cryo-EM structure determination. Nat Methods 14, 290–296 (2017). 10.1038/nmeth.4169

87 Punjani, A., Brubaker, M. A. & Fleet, D. J. Building Proteins in a Day: Efficient 3D Molecular Structure Estimation with Electron Cryomicroscopy. IEEE Trans Pattern Anal Mach Intell 39, 706–718 (2017). 10.1109/TPAMI.2016.2627573

88 Grant, T., Rohou, A. & Grigorieff, N. cisTEM, user-friendly software for single-particle image processing. Elife 7 (2018). 10.7554/eLife.35383

89 Liebschner, D. et al. Macromolecular structure determination using X-rays, neutrons and electrons: recent developments in Phenix. Acta Crystallogr D Struct Biol 75, 861–877 (2019). 10.1107/S2059798319011471

90 Waterhouse, A. et al. SWISS-MODEL: homology modelling of protein structures and complexes. Nucleic Acids Res 46, W296–W303 (2018). 10.1093/nar/gky427

91 Jumper, J. et al. Highly accurate protein structure prediction with AlphaFold. Nature 596, 583–589 (2021). 10.1038/s41586-021-03819-2

92 Bertram, K. et al. Cryo-EM structure of a human spliceosome activated for step 2 of splicing. Nature 542, 318–323 (2017). 10.1038/nature21079

93 Jamali, K. et al. Automated model building and protein identification in cryo-EM maps. Nature 628, 450–457 (2024). 10.1038/s41586-024-07215-4

94 Pettersen, E. F. et al. UCSF Chimera--a visualization system for exploratory research and analysis. J Comput Chem 25, 1605–1612 (2004). 10.1002/jcc.20084

95 Emsley, P., Lohkamp, B., Scott, W. G. & Cowtan, K. Features and development of Coot. Acta Crystallogr D Biol Crystallogr 66, 486–501 (2010). 10.1107/S0907444910007493

96 Sievers, F. et al. Fast, scalable generation of high-quality protein multiple sequence alignments using Clustal Omega. Mol Syst Biol 7, 539 (2011). 10.1038/msb.2011.75

97 Buchan, D. W. A. & Jones, D. T. The PSIPRED Protein Analysis Workbench: 20 years on. Nucleic Acids Res 47, W402–W407 (2019). 10.1093/nar/gkz297

98 Chen, V. B. et al. MolProbity: all-atom structure validation for macromolecular crystallography. Acta Crystallogr D Biol Crystallogr 66, 12–21 (2010). 10.1107/S0907444909042073

99 Chen, Z. L. et al. A high-speed search engine pLink 2 with systematic evaluation for proteome-scale identification of cross-linked peptides. Nat Commun 10, 3404 (2019). 10.1038/s41467-019-11337-z

100 Kim, D., Paggi, J. M., Park, C., Bennett, C. & Salzberg, S. L. Graph-based genome alignment and genotyping with HISAT2 and HISAT-genotype. Nat Biotechnol 37, 907–915 (2019). 10.1038/s41587-019-0201-4

101 Li, H. et al. The Sequence Alignment/Map format and SAMtools. Bioinformatics 25, 2078–2079 (2009). 10.1093/bioinformatics/btp352

102 Ramirez, F., Dundar, F., Diehl, S., Gruning, B. A. & Manke, T. deepTools: a flexible platform for exploring deep-sequencing data. Nucleic Acids Res 42, W187–191 (2014). 10.1093/nar/gku365

103 Burke, J. E. et al. Spliceosome Profiling Visualizes Operations of a Dynamic RNP at Nucleotide Resolution. Cell 173, 1014–1030 e1017 (2018). 10.1016/j.cell.2018.03.020

104 Charif, D. & Lobry, J. R. SeqinR 1.0-2: a contributed package to the R project for statistical computing devoted to biological sequences retrieval and analysis. Structural approaches to sequence evolution: Molecules, networks, populations, 207–232 (2007).

105 Haider, S. et al. A bedr way of genomic interval processing. Source code for biology and medicine 11, 1–7 (2016).

106 Studier, F. W. Protein production by auto-induction in high density shaking cultures. Protein Expr Purif 41, 207–234 (2005). 10.1016/j.pep.2005.01.016

