## Supplementary Information for "Structures of aberrant spliceosome intermediates on their way to disassembly"

### Extended Data

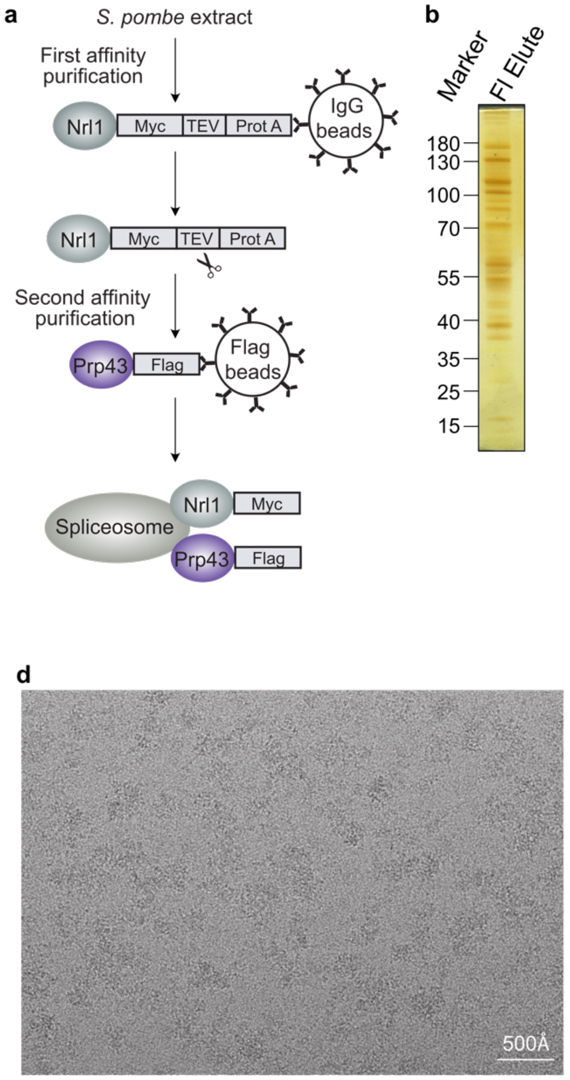


c

Mass spectrometry analysis

| Sub-complex | Accession ID | Protein name | Mol. Wt. (kDa) | PSMs 1 | PSMs 2 | PSMs 3 | Modeled($sp$B$^d$) I | Modeled($sp$B$^d$) II |
|---|---|---|---|---|---|---|---|---|
| U5 snRNP | O14187 | Prp8 | 275 | 247 | 257 | 247 | ✓ | ✓ |
| | O94316 | Snu114 | 111 | 138 | 142 | 143 | ✓ | ✓ |
| | Q9UT24 | Brr2 | 249 | 164 | 169 | 150 | x | x |
| | O94620 | Cwf17 | 37 | 48 | 43 | 43 | ✓ | ✓ |
| | Q9UUC6 | SmD3 | 11 | 12 | 15 | 15 | ✓ | ✓ |
| | Q10163 | SmB | 15 | 8 | 9 | 10 | ✓ | ✓ |
| | O42661 | SmD1 | 13 | 7 | 5 | 7 | ✓ | ✓ |
| | O14036 | SmD2 | 13 | 10 | 8 | 12 | ✓ | ✓ |
| | Q9USZ3 | SmE | 10 | 4 | 5 | 4 | ✓ | ✓ |
| | O59734 | SmF | 9 | 9 | 9 | 10 | ✓ | ✓ |
| | O74966 | SmG | 9 | 3 | 3 | 3 | ✓ | ✓ |
| U2 snRNP core | Q9UUC6 | SmD3 | 11 | 12 | 15 | 15 | x | x |
| | Q10163 | SmB | 15 | 8 | 9 | 10 | x | x |
| | O42661 | SmD1 | 13 | 7 | 5 | 7 | x | x |
| | O14036 | SmD2 | 13 | 10 | 8 | 12 | x | x |
| | Q9USZ3 | SmE | 10 | 4 | 5 | 4 | x | x |
| | O59734 | SmF | 9 | 9 | 9 | 10 | x | x |
| | O74966 | SmG | 9 | 3 | 3 | 3 | x | x |
| | Q9USX8 | Lea1 | 27 | 22 | 21 | 20 | x | x |
| | Q7LL14 | Msl1 | 13 | 7 | 7 | 7 | x | x |
| NTC core | P87312 | Clf1 | 81 | 81 | 80 | 84 | ✓ | ✓ |
| | O14011 | Prp19 | 54 | 62 | 68 | 72 | ✓ | ✓ |
| | P39964 | Cef1 | 87 | 71 | 68 | 63 | ✓ | ✓ |
| | Q9P7R9 | Syf1 | 93 | 57 | 49 | 46 | ✓ | ✓ |
| | Q9USV3 | Snt309 | 21 | 24 | 25 | 25 | ✓ | ✓ |
| | O59733 | Syf2 | 28 | 34 | 33 | 35 | ✓ | ✓ |
| | O74370 | Isy1 | 26 | 25 | 28 | 29 | x | x |
| NTC related | O13615 | Prp46 | 52 | 69 | 69 | 65 | ✓ | ✓ |
| | Q09882 | Prp45 | 63 | 67 | 71 | 62 | ✓ | ✓ |
| | O59800 | Emc2 | 40 | 35 | 38 | 31 | ✓ | ✓ |
| | O94508 | Cwf11 | 148 | 75 | 74 | 78 | ✓ | ✓ |
| | O74772 | Bud31 | 17 | 23 | 19 | 25 | ✓ | ✓ |
| | P87126 | Cwc2 | 44 | 22 | 23 | 26 | ✓ | ✓ |
| | P78794 | Cwc15 | 30 | 22 | 21 | 19 | ✓ | ✓ |
| Splicing factors | O43071 | Prp17 | 63 | 49 | 56 | 53 | ✓ | ✓ |
| | O14161 | Cwc21 | 35 | 16 | 16 | 15 | ✓ | ✓ |
| | Q9P6R9 | Cwc22 | 103 | 57 | 53 | 58 | ✓ | ✓ |
| | P87051 | Cyp1 | 17 | 15 | 16 | 14 | ✓ | ✓ |
| | O60141 | Saf4 | 35 | 22 | 22 | 28 | x | ✓ |
| | O59793 | Bis1 | 43 | 9 | 12 | 10 | x | ✓ |
| Ntr1 complex | O42945 | Prp43 | 89 | 174 | 185 | 184 | x | x |
| | Q9UTK6 | Ntr1 | 92 | 47 | 49 | 45 | * | ✓ |
| | O13758 | Ntr2 | 42 | 18 | 19 | 19 | x | ✓ |
| | Q9P7C6 | Cwc23 | 35 | 18 | 18 | 16 | x | x |
| CNM complex | O42975 | Nrl1 | 135 | 52 | 48 | 48 | x | x |
| | O13799 | Mtl1 | 118 | 60 | 57 | 65 | x | x |
| | Q9P7B7 | Ctr1 | 35 | 24 | 27 | 28 | x | x |
| Gih35-Gpl1 | Q9HE06 | Gih35 | 73 | 39 | 42 | 35 | ✓ | ✓ |
| | Q9HE07 | Gpl1 | 61 | 40 | 43 | 40 | ✓ | ✓ |

**Extended Data Fig. 1: Purification and characterization of the *sp*B$^d$ complex.**

**a,** Purification scheme for the *sp*B$^d$ complex using a split-tag approach. **b,** After elution of the *sp*B$^d$ complex from the anti-FLAG beads, the sample was analyzed using SDS-PAGE and visualized by silver staining. **c,** The purified *sp*B$^d$ complex was crosslinked with BS3 and subjected to mass spectrometric analysis. A list of the components of the U5 snRNP, U2 snRNP core, NTC core, NTC related proteins, other splicing factors, Ntr1 complex, CNM complex and the associated Gih35-Gpl1 proteins identified in the *sp*B$^d$ complex is provided. The PSM (Peptide-spectrum match) values obtained from three technical replicates are indicated. A list of the top 200 proteins identified in the sample, ranked according to label-free quantitation (LFQ) intensity, are provided in Supplementary Table 1. The proteome analysis of the *sp*B$^d$ complex was also performed on non-crosslinked samples which yielded a similar protein composition as shown here. Proteins modelled in the *sp*B$^d$-I and II complexes are marked. * represents that only the C-terminal domain of Ntr1 was modelled in the *sp*B$^d$-I state. **d,** A representative cryo-EM micrograph. Scale bar represents 500 Å.

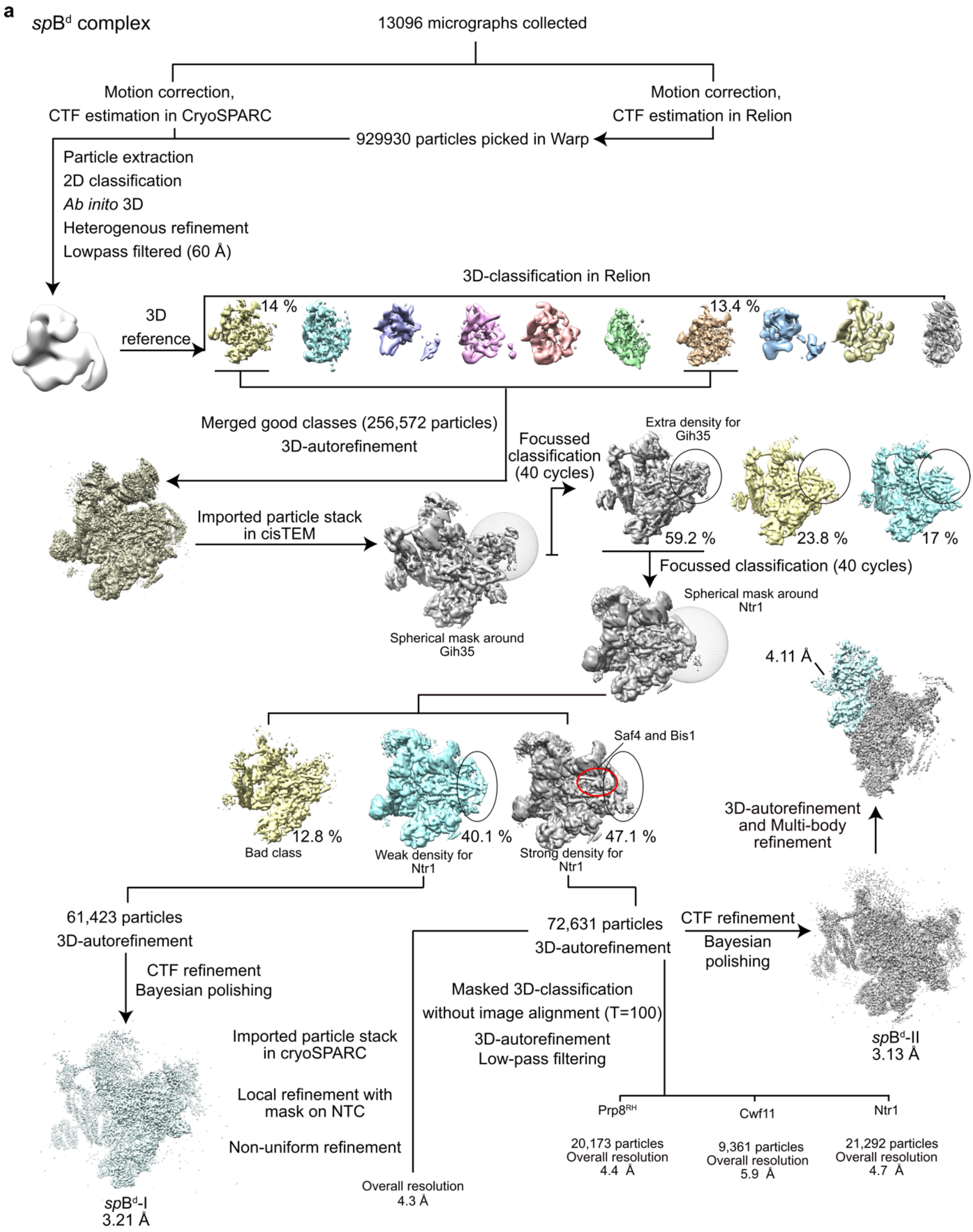
a
spB$^{d}$ complex
13096 micrographs collected
Motion correction,
CTF estimation in CryoSPARC
Motion correction,
CTF estimation in Relion
929930 particles picked in Warp
Particle extraction
2D classification
Ab inito 3D
Heterogenous refinement
Lowpass filtered (60 Å)
3D-classification in Relion
3D reference
14 %
13.4 %
Merged good classes (256,572 particles)
3D-autorefinement
Focussed classification (40 cycles)
Extra density for Gih35
59.2 %
23.8 %
17 %
Imported particle stack in cisTEM
Spherical mask around Gih35
Focussed classification (40 cycles)
Spherical mask around Ntr1
4.11 Å
Saf4 and Bis1
12.8 %
Bad class
40.1 %
Weak density for Ntr1
47.1 %
Strong density for Ntr1
3D-autorefinement and Multi-body refinement
61,423 particles
3D-autorefinement
CTF refinement
Bayesian polishing
72,631 particles
3D-autorefinement
CTF refinement
Bayesian polishing
Masked 3D-classification
without image alignment (T=100)
3D-autorefinement
Low-pass filtering
Imported particle stack in cryoSPARC
Local refinement with mask on NTC
Non-uniform refinement
Overall resolution 4.3 Å
spB$^{d}$-II
3.13 Å
Prp8$^{RH}$
20,173 particles
Overall resolution 4.4 Å
Cwf11
9,361 particles
Overall resolution 5.9 Å
Ntr1
21,292 particles
Overall resolution 4.7 Å
spB$^{d}$-I
3.21 Å

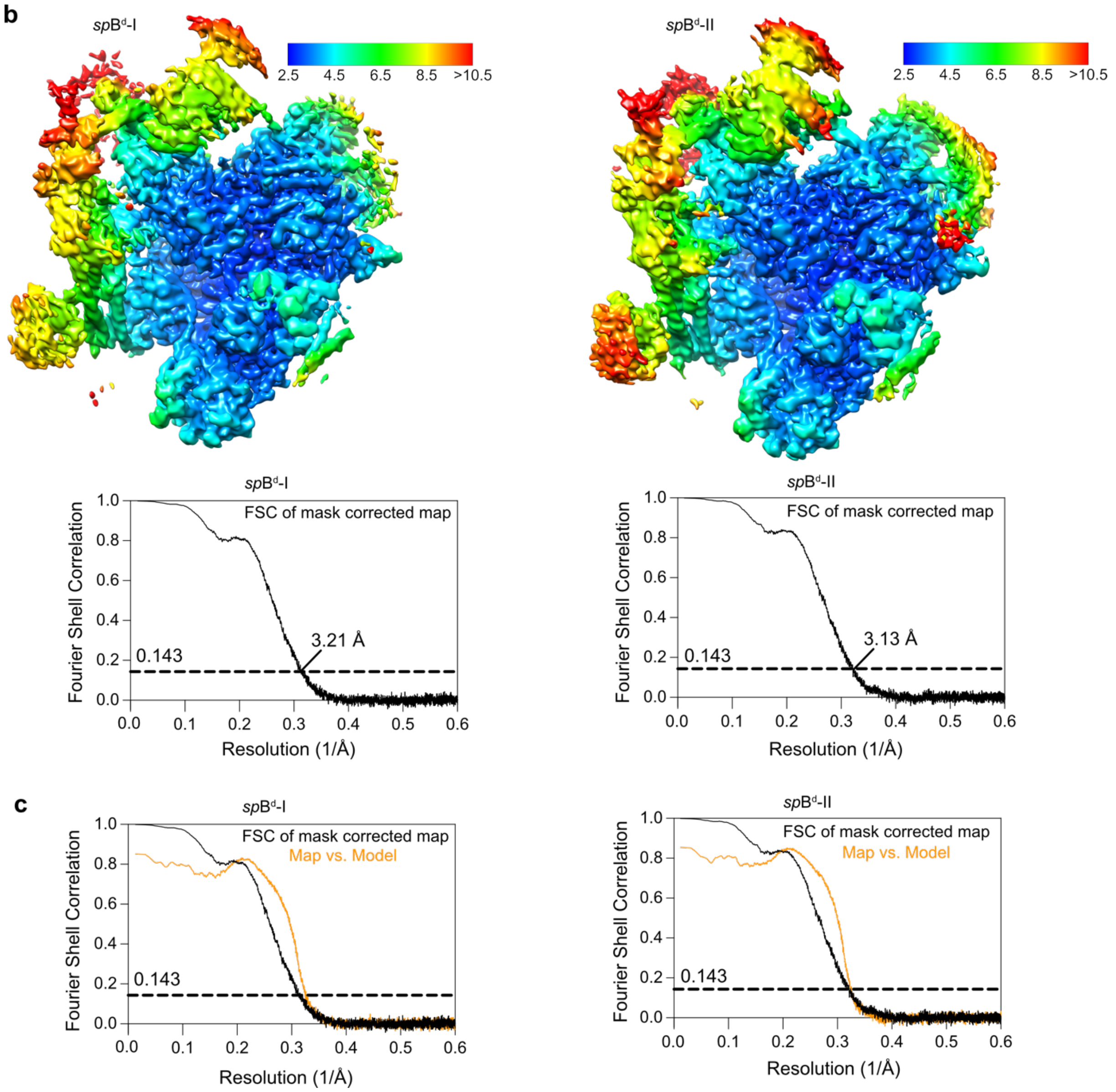
b
spBd-I
spBd-II
2.5 4.5 6.5 8.5 >10.5
FSC of mask corrected map
3.21 Å
3.13 Å
0.143
Fourier Shell Correlation
Resolution (1/Å)
c
Map vs. Model

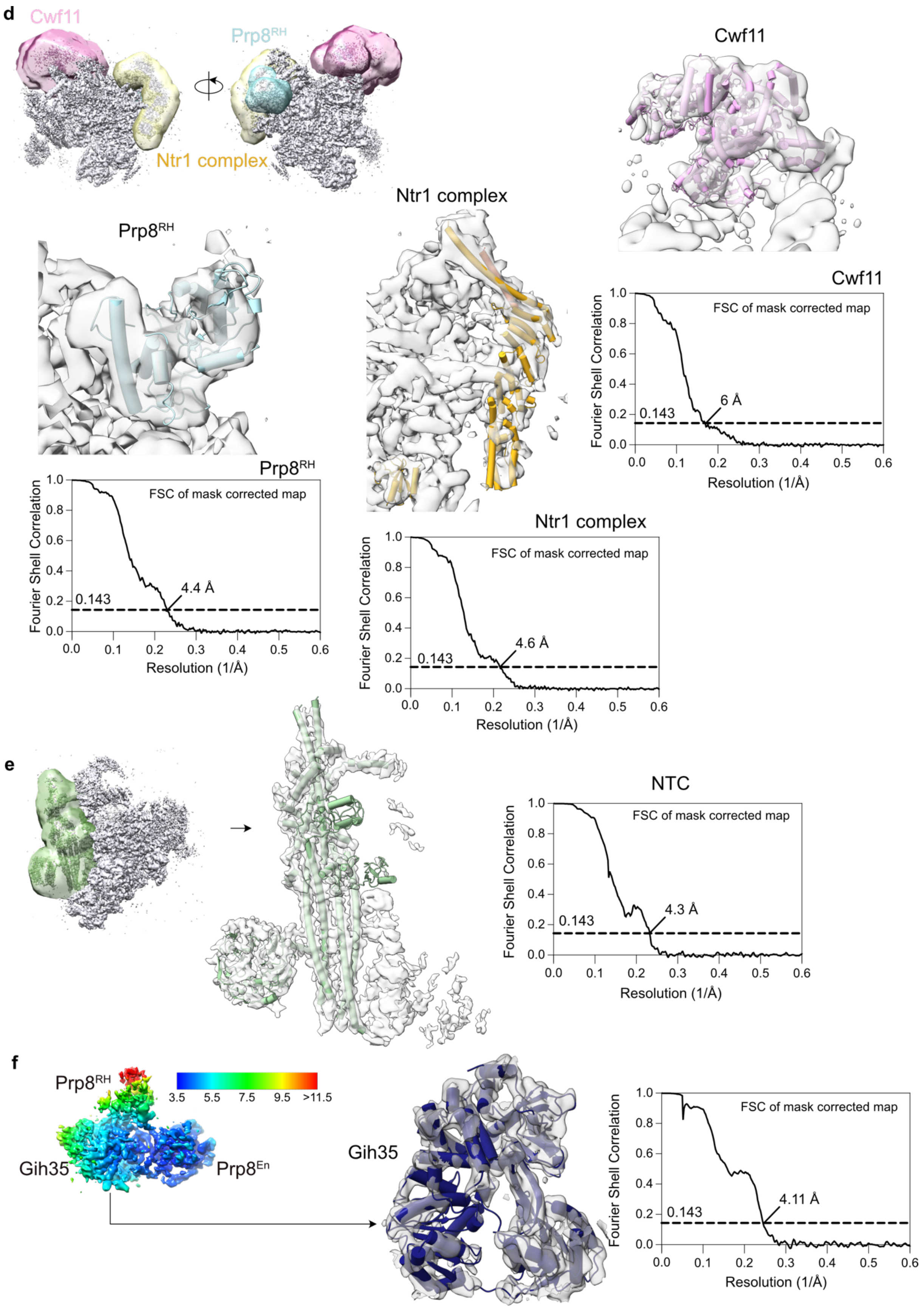
d
Cwf11
Prp8RH
Ntr1 complex
Cwf11
Ntr1 complex
Prp8RH
Cwf11
FSC of mask corrected map
Fourier Shell Correlation
Resolution (1/Å)
0.143
6 Å
Prp8RH
FSC of mask corrected map
Fourier Shell Correlation
Resolution (1/Å)
0.143
4.4 Å
Ntr1 complex
FSC of mask corrected map
Fourier Shell Correlation
Resolution (1/Å)
0.143
4.6 Å
e
NTC
FSC of mask corrected map
Fourier Shell Correlation
Resolution (1/Å)
0.143
4.3 Å
f
Prp8RH
3.5 5.5 7.5 9.5 >11.5
Gih35
Prp8En
Gih35
FSC of mask corrected map
Fourier Shell Correlation
Resolution (1/Å)
0.143
4.11 Å

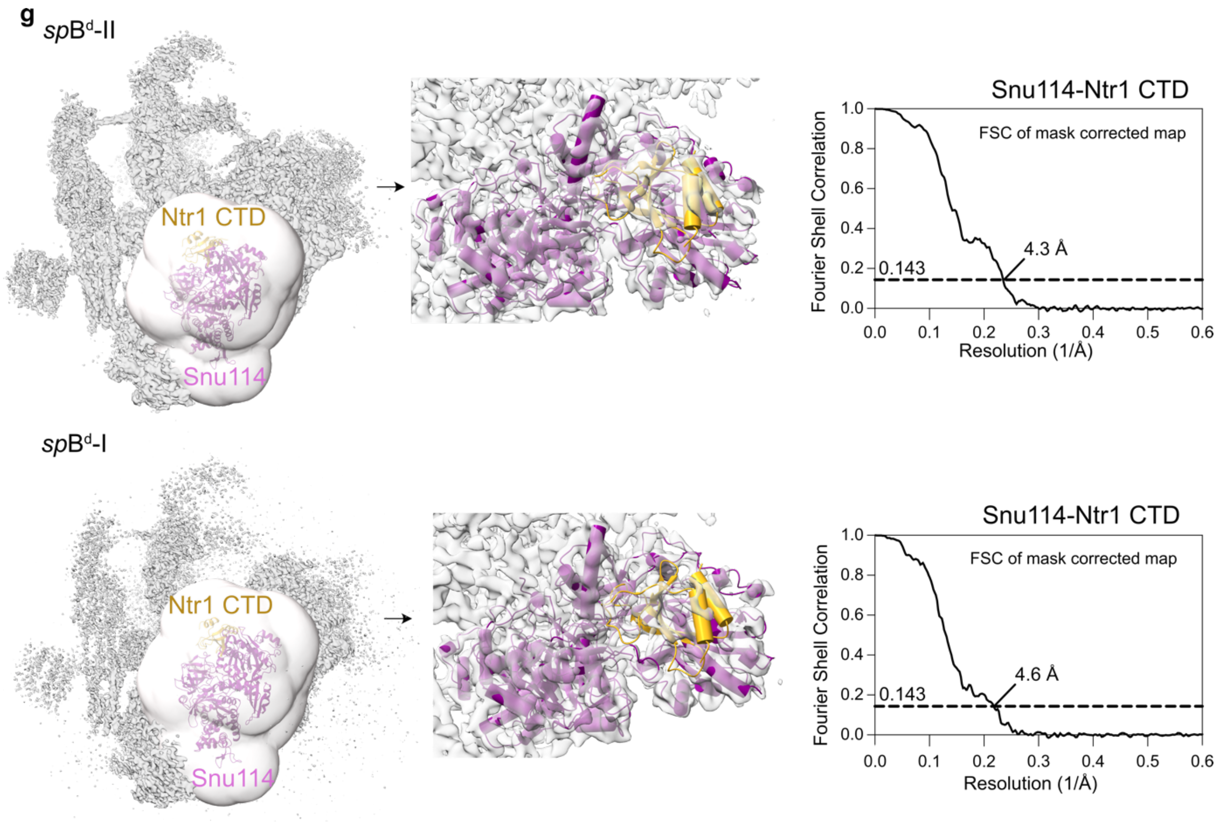


**Extended Data Fig. 2: Cryo-EM processing scheme.**
**a,** Cryo-EM processing pipeline for the *sp*B$^{d}$ complex. **b,** Local resolution and Fourier shell correlation curves (FSC) (0.143 cut-off) along with resolutions are reported for the consensus refinements of *sp*B$^{d}$-I and *sp*B$^{d}$-II states. **c,** A superposition of the FSC curves of the consensus cryo-EM map (black) and the atomic model (orange) is shown for *sp*B$^{d}$-I and *sp*B$^{d}$-II states. **d,** Masks on the consensus refinement of *sp*B$^{d}$-II complex used for masked 3D-classification shown in panel (a) for Prp8$^{RH}$, Ntr1 complex and Cwf11 are provided. The corresponding superposition of the models with their respective cryo-EM maps obtained after unmasked 3D refinement is shown along with FSC curves (0.143 cut-off) and resolution estimates. **e,** Local refinement with a mask around the NTC led to an improved resolution. Superposition of NTC with the cryo-EM maps is shown along with FSC curve (0.143 cut-off) and resolution estimate. **f,** Local resolution and Fourier shell correlation curves (FSC) (0.143 cut-off) along with resolutions are reported for multibody refinement performed on the *sp*B$^{d}$-II state. A superposition of the Gih35 model with the cryo-EM map is also provided. **g,** Focused 3D classification and local refinement with a mask around Snu114 and Ntr1 CTD led to an improved resolution for Ntr1 CTD. Superposition of Snu114 and Ntr1 CTD with the *sp*B$^{d}$-II and *sp*B$^{d}$-I cryo-EM maps is shown along with FSC curve (0.143 cut-off) and resolution estimates.

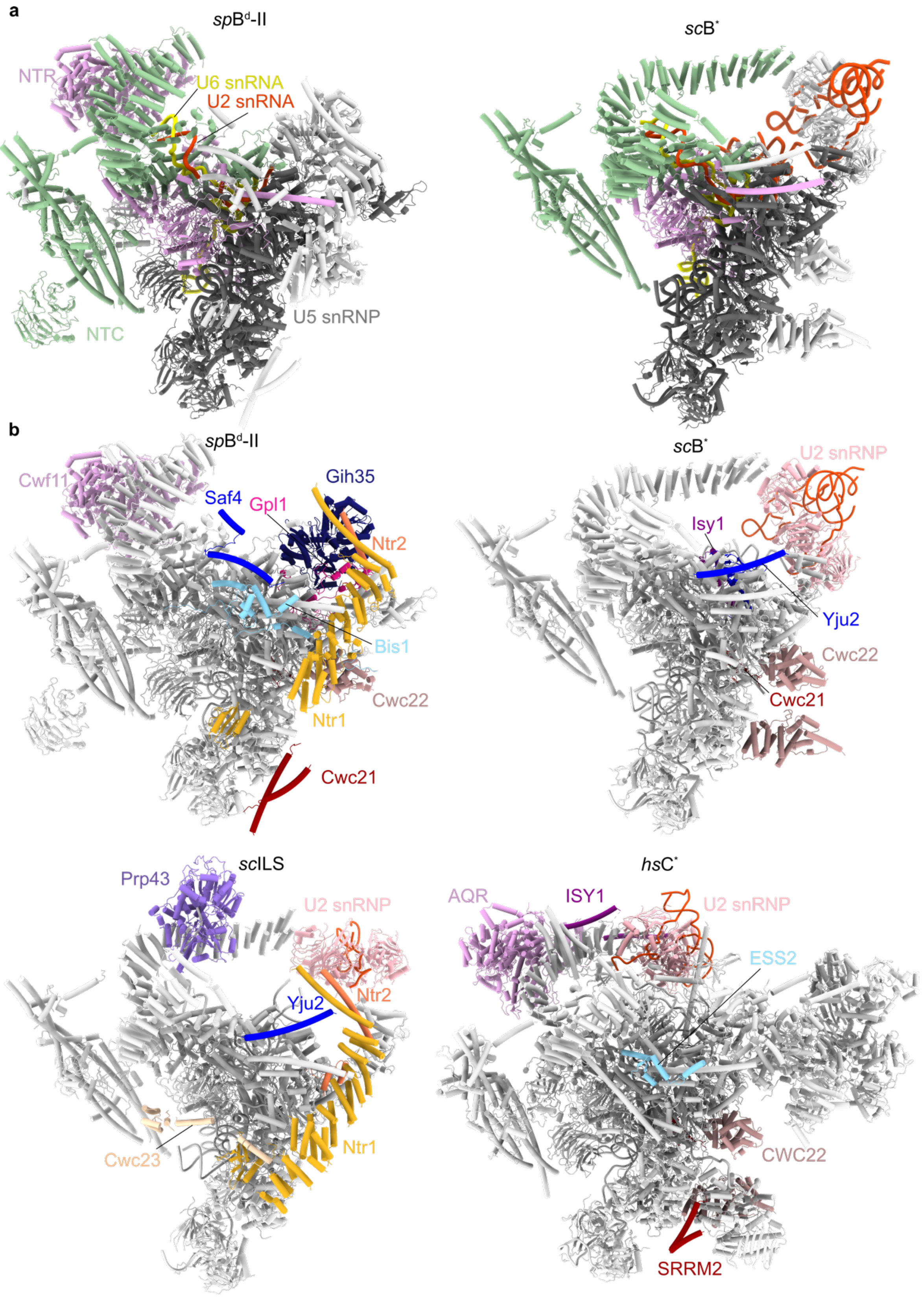


**Extended Data Fig. 3: Comparison of *sp*B$^{d}$-II state with other spliceosome complexes**
**a,** Comparison of overall structures of the *sp*B$^{d}$-II state and *sc*B* (PDB:6J6Q[1]) shows the stable core of the spliceosome that remains largely unchanged. **b,** Comparison of overall structures of the *sp*B$^{d}$-II state and *sc*B* (PDB:6J6Q[1]), *sc*ILS (PDB:5Y88[2]) and *hs*C* (PDB:8C6J[3]) complexes. Components differing in the four states are shown in color.

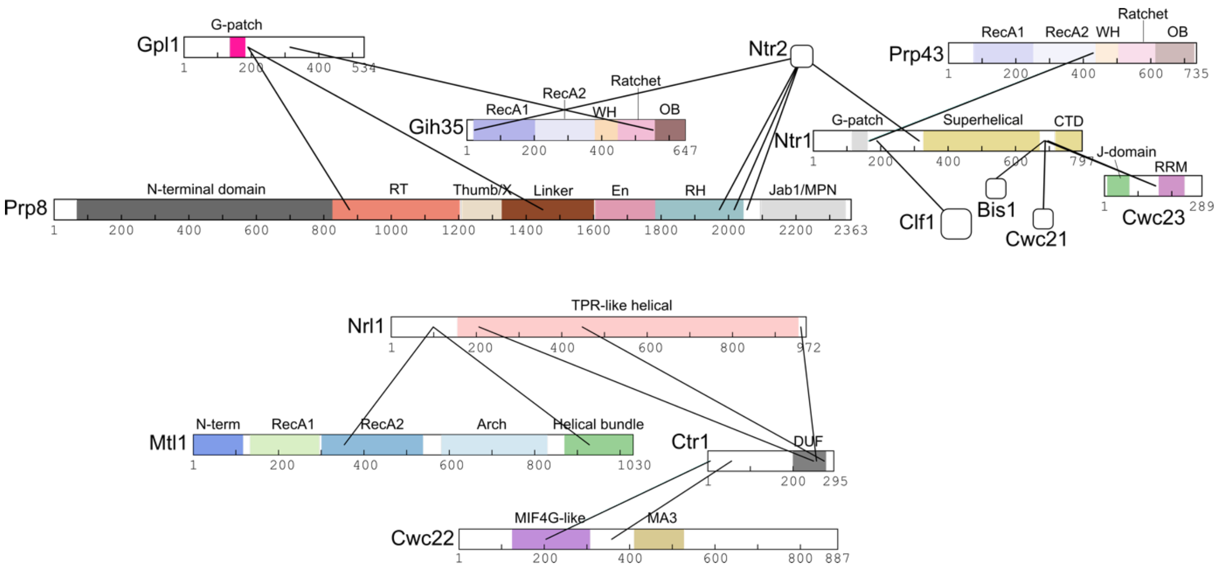


**Extended Data Fig. 4: Inter-protein crosslinks identified within the *sp*B$^{d}$ complex.**

Schematic representation of a subset of inter-protein crosslinks originating from Gpl1, Gih35, Ntr1 complex (Ntr1, Ntr2, Prp43, Cwc23) and the CNM complex (Ctr1, Nrl1, Mtl1) obtained using the entire *S. pombe* proteome as a search database. See Supplementary Table 1 for a complete list of all intra-and inter-protein crosslinks identified. Domain annotations in the proteins are shown. Gpl1 contains a G-patch domain while Gih35 contains RecA1, RecA2, WH (winged-helix), Ratchet and OB-fold (oligonucleotide/oligosaccharide-binding fold) domains. Prp43 contains RecA1, RecA2, WH, Ratchet and OB-fold domains while Ntr1 contains a G-patch domain in addition to a superhelical domain and a CTD (C-terminal domain). Prp8 contains an N-terminal domain, RT (reverse transcriptase) fingers/palm domain, Thumb/X, Linker, En (endonuclease), RH (RNaseH-like) and Jab1/MPN domains. Cwc23 contains a J-domain and an RRM (RNA recognition motif) domain. Nrl1 contains a TPR-like (tetratricopeptide repeat-like) helical domain while Mtl1 contains an N-terminal domain, RecA1, RecA2, Arch and a helical bundle. Ctr1 contains a DUF domain (domain of unknown function). Finally, Cwc22 contains an MIF4G-like domain and an MA3 domain.

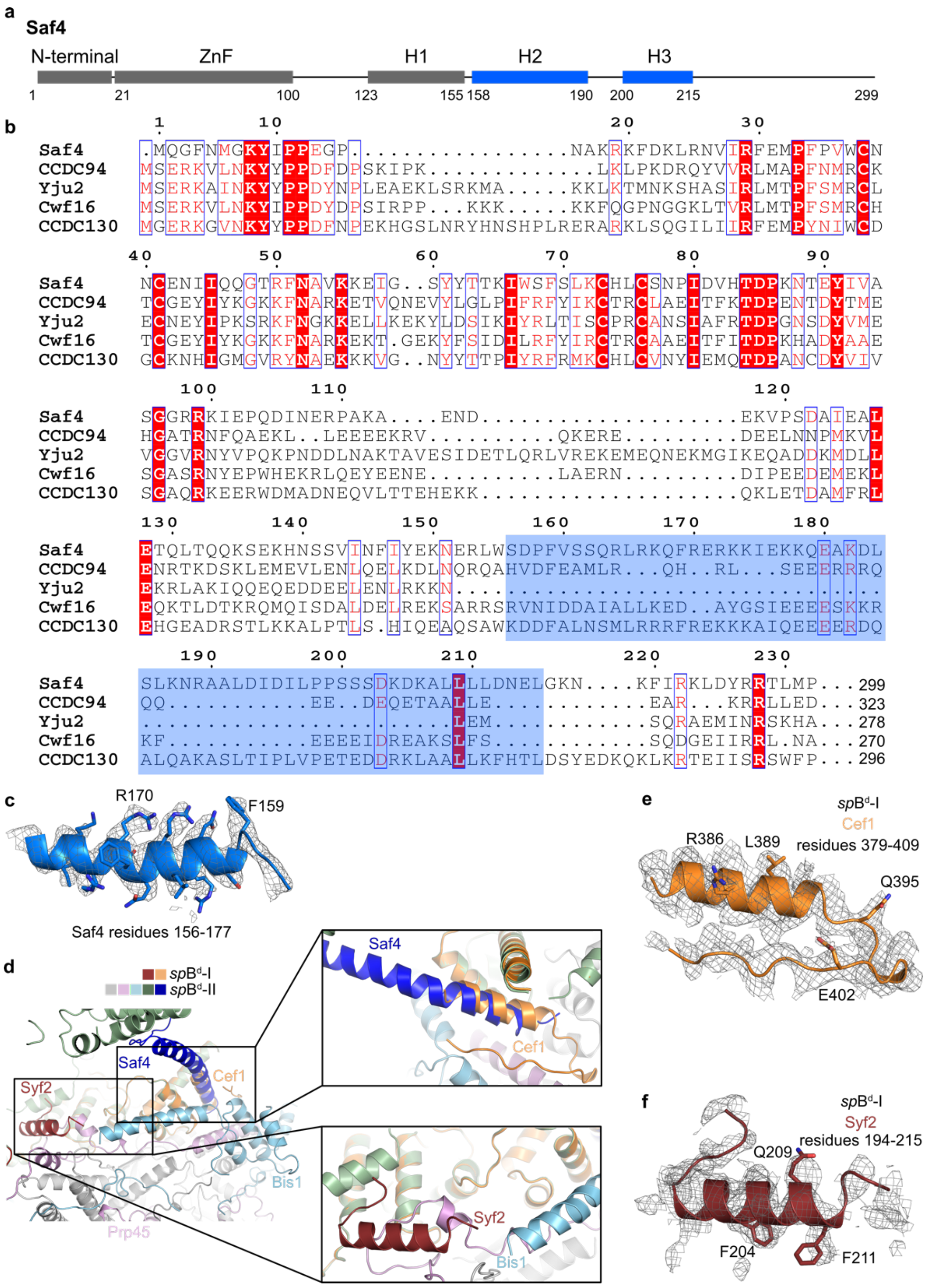


**Extended Data Fig. 5: Saf4 and Bis1 are incorporated in the *sp*B$^{d}$-II state**

**a,** Domain architecture of Saf4. **b,** Homologs of *sc*Yju2. Saf4 and Cwf16 in *S. pombe*, CCDC94 and CCDC130 in humans share sequence homology with *sc*Yju2. The two α-helices of Saf4 visible in the *sp*B$^{d}$-II state are highlighted in blue. **c,** Superposition of Saf4 residues 156-177 with the *sp*B$^{d}$-II cryo-EM map. **d,** Structure superposition of *sp*B$^{d}$-I and -II structures. Color scheme of *sp*B$^{d}$-II state follows Fig. 1. From the *sp*B$^{d}$-I state, only Syf2 and Cef1 are shown in dark red and orange, respectively. Insets show zoomed-in views of structural differences between the two states. Cef1 residues 379-409 in *sp*B$^{d}$-I state are displaced by Saf4 in the *sp*B$^{d}$-II state. Syf2 residues 194-215 are destabilized by the presence of Bis1 in *sp*B$^{d}$-II state. **e,** Superposition of Cef1 residues 379-409 with the *sp*B$^{d}$-I cryo-EM map. **f,** Superposition of Syf2 residues 194-215 with the *sp*B$^{d}$-I cryo-EM map.

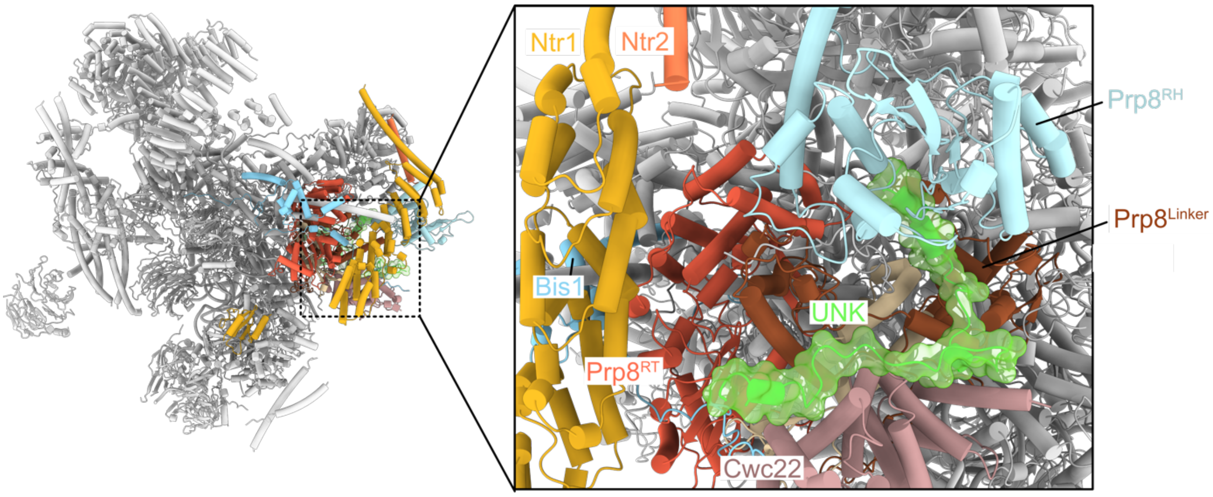


**Extended Data Fig. 6: Interactions of UNK within the *sp*B$^{d}$-II state.**
Additional cryo-EM density exists near Prp8, Cwc22 and the Ntr1 complex, which could not be reliably assigned and is referred to as unknown (UNK). It should be noted that fragments of different proteins could possibly contribute to UNK.

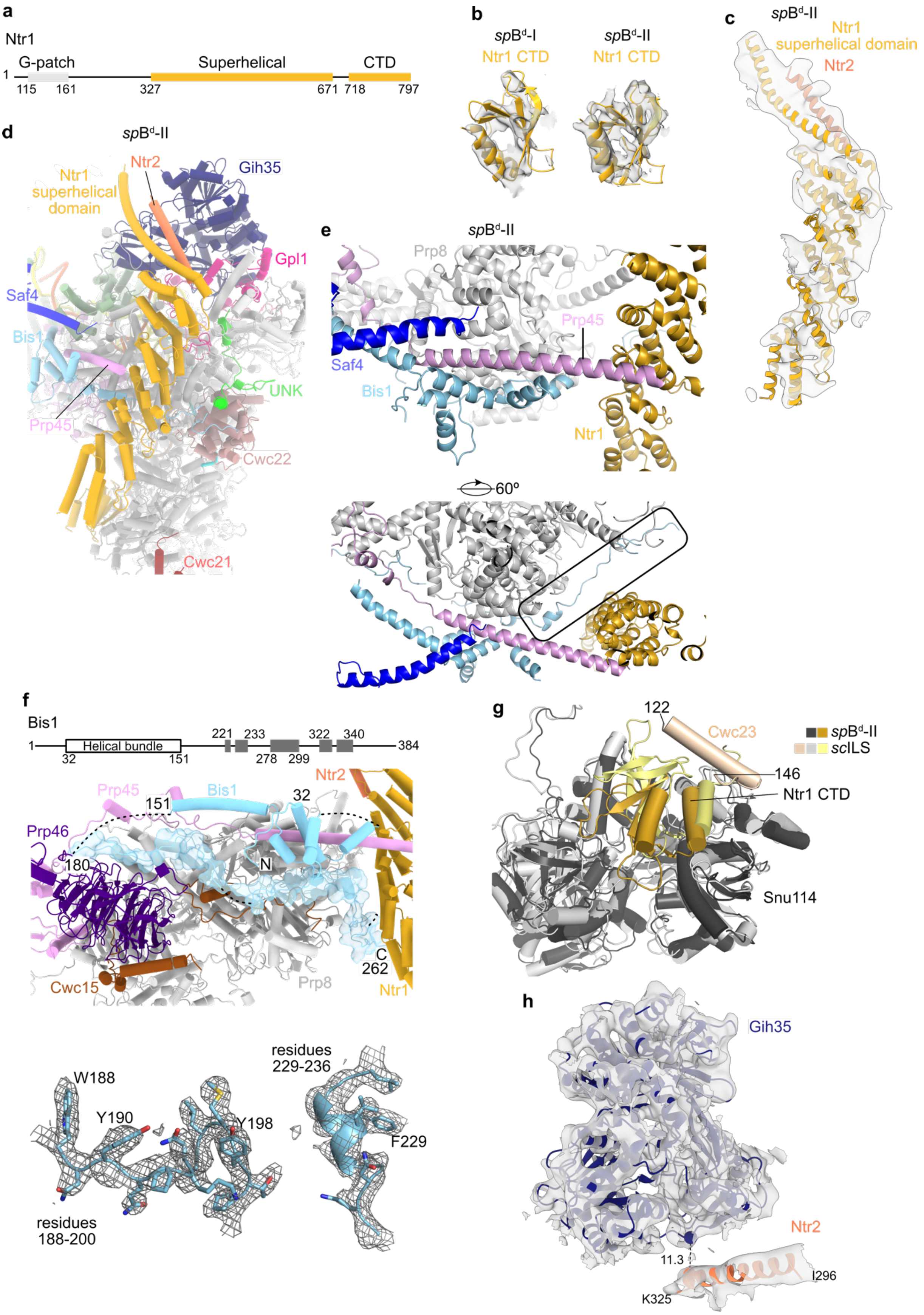


**Extended Data Fig. 7: The Ntr1 complex is stabilized in the *sp*B$^{d}$-II state.**

**a,** Schematic representation of the domain architecture of Ntr1. **b,** Superimposition of Ntr1 CTD from *sp*B$^{d}$-I and -II states with their respective cryo-EM maps. **c,** Superimposition of Ntr1 superhelical domain and the Ntr2 helix (residues 296-325) with the *sp*$B^d$-II cryo-EM map. **d,** The superhelical domain of Ntr1 contacts Ntr2, Prp45 and Bis1 and is positioned close to Gih35 in the *sp*$B^d$-II complex. **e,** Bis1 (sky blue) and Saf4 (dark blue) stabilize the long α-helix of Prp45 interacting with Ntr1 superhelical domain. The stretch of Bis1 directly binding to Ntr1 is boxed. **f,** Domain architecture of Bis1 is shown with the helical bundle marked and other helices shown as gray boxes in the top panel. The middle panel shows residues 22-262 (including gaps) of Bis1 that can be traced in the *sp*$B^d$-II state. Apart from the helical bundle, ~80 residues of Bis1 can be traced (shown also as surface). Superposition of Bis1 residues 188-200 and 229-236 with the *sp*$B^d$-II cryo-EM map is provided in the bottom panel. **g,** The Ntr1 CTD is anchored on Snu114. In the *sc*ILS complex, this association is stabilized by an α-helix of Cwc23 (residues 122-146). **h,** Superimposition of Gih35 and Ntr2 with their respective focused cryo-EM maps. The distance (11.3 Å) between Cα atoms of closest residues Pro 206 (Gih35) and Ile 323 (Ntr2) is shown with a dotted line.

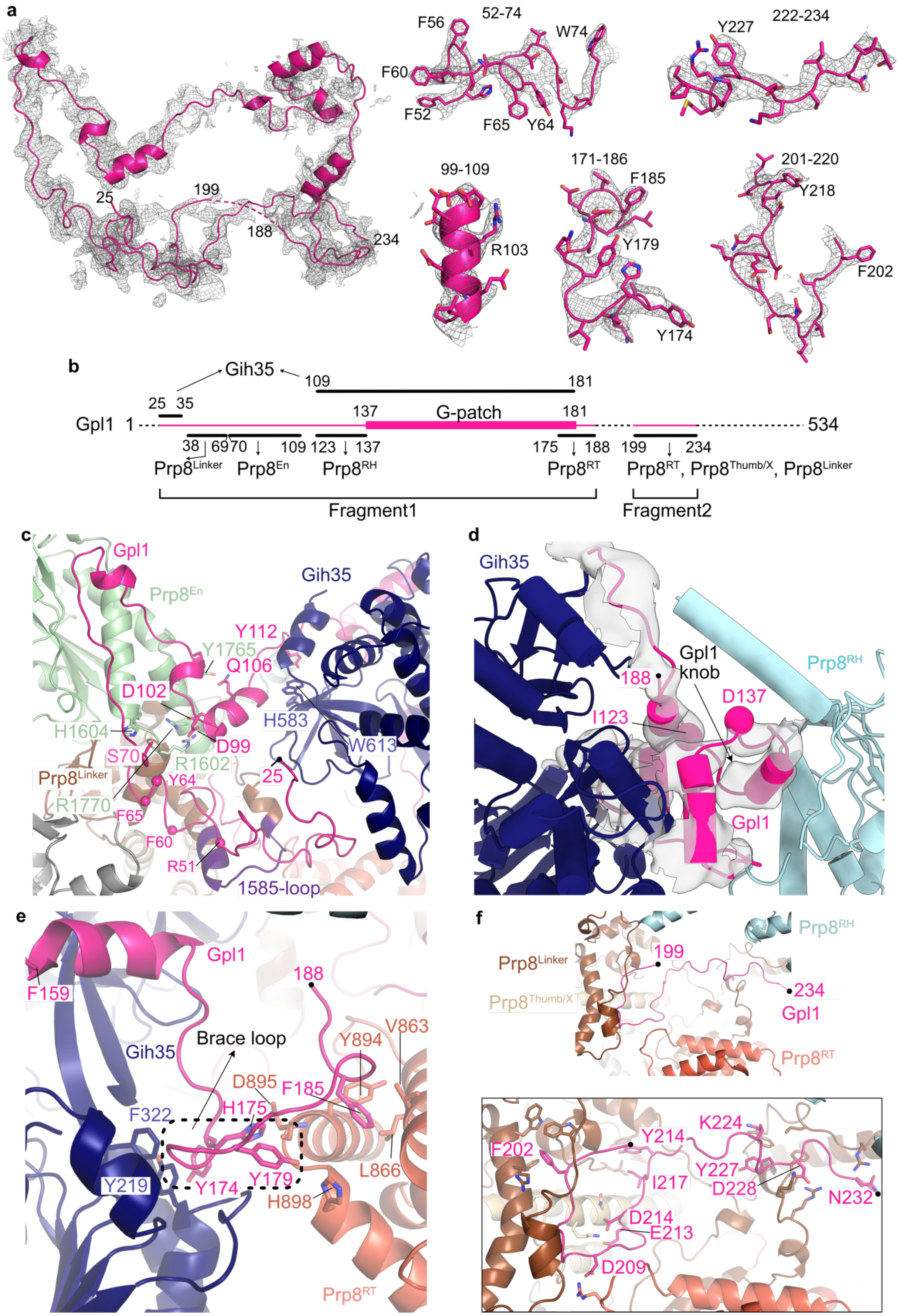
a
52-74
F56
W74
F60
F52
F65
Y64
222-234
Y227
25
199
188
234
99-109
R103
171-186
F185
Y179
Y174
201-220
Y218
F202
b
Gih35
109
181
25 35
137
G-patch
181
Gpl1 1
534
38 69 70 109 123 137
175 188 199 234
Prp8Linker Prp8En Prp8RH
Prp8RT Prp8RT, Prp8Thumb/X, Prp8Linker
Fragment1
Fragment2
c
Gpl1
Prp8En
Gih35
Y112
Y1765
Q106
D102
H1604
H583
D99
W613
S70
R1602
Prp8Linker
Y64
25
R1770
F65
F60
R51
1585-loop
d
Gih35
Gpl1 knob
Prp8RH
188
D137
I123
Gpl1
e
Gpl1
F159
188
Gih35
Brace loop
V863
Y894
D895
F185
F322
H175
L866
Y219
Y174
Y179
H898
Prp8RT
f
Prp8RH
199
Prp8Linker
234
Prp8Thumb/X
Gpl1
Prp8RT
F202
Y214
K224
Y227
I217
D228
N232
D214
E213
D209

**Extended Data Fig. 8: Gpl1 interacts with Prp8.**

**a,** Cryo-EM map for the entire Gpl1 trace visible in the *sp*B$^{d}$-II complex (residues 25-188, 199-234). Zoomed-in views of representative regions of Gpl1 for *sp*B$^{d}$ interaction are shown as insets. **b,** Gpl1 forms a bridge between Prp8 and Gih35. Regions largely adjacent to the G-patch domain contribute to the interaction. The boundaries of different regions of Gpl1 interacting with the helicase Gih35 and distinct domains of Prp8 are shown (detailed in panels c to f where N- and C-termini of the two Gpl1 fragments visible in the *sp*B$^{d}$ complex are marked with black dots). **c,** Interactions between the Gpl1 N-terminal region and Prp8$^{En}$ and Prp8$^{Linker}$ (including the 1585-loop). the The C$\alpha$ atoms of residues contacting the pre-mRNA in the active site of the spliceosome are shown as spheres. **d,** Gpl1 residues 123-137 form a knob and insert into Prp8$^{RH}$, thus sequestering the Prp8 domain in this extreme position. An overlay of the cryo-EM map with Gpl1 residues 109-188 is provided, and the C$\alpha$ atoms of Gpl1 knob residues I123 and D137 are shown as spheres. **e,** Interactions between the Gpl1 region including the brace loop and Prp8$^{RT}$ (see also Extended Data Fig. 12). **f,** Interactions between Gpl1 residues 199-234 and an interface formed by the Prp8$^{RT}$, Prp8$^{Thumb/X}$ and Prp8$^{Linker}$ domains. Inset shows a zoom-in of the interactions. For simplicity, only residues from Gpl1 are labeled.

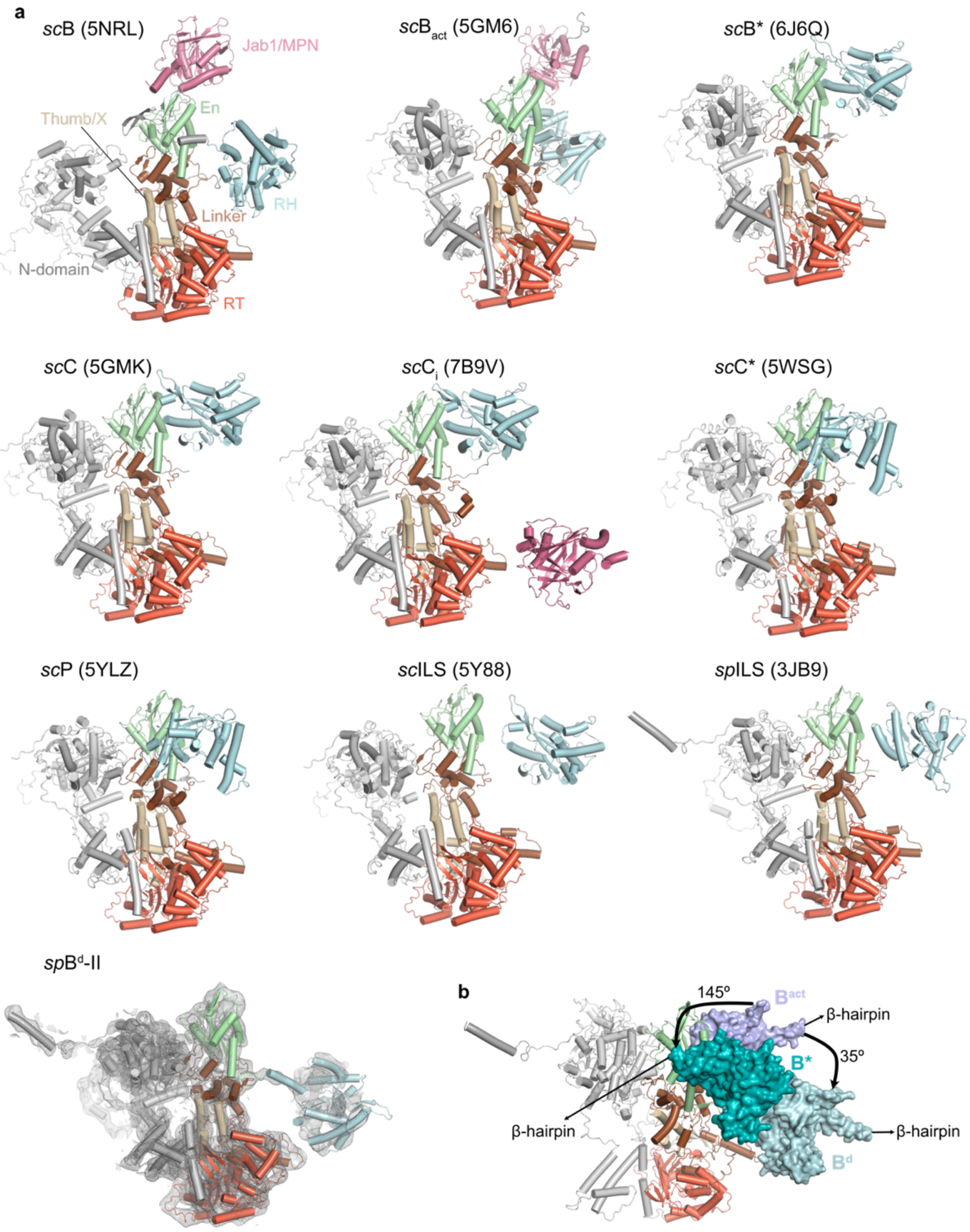


**Extended Data Fig. 9: Conformation sampling by the Prp8$^{RH}$.**

**a,** Arrangement of the N-domain (gray), RT (orange), Thumb/X (wheat), Linker (brown), En (green), RH (pale blue) and Jab1/MPN (pink) domains of Prp8 from the *sc*B-*sc*/*sp*ILS and the *sp*B$^{d}$-II state is shown. For *sp*B$^{d}$-II Prp8, an overlay of the cryo-EM map is provided. The respective PDB IDs are indicated. **b,** Superposition of Prp8 from the *sc*B$^{act}$, *sc*B*, and the *sp*B$^{d}$-II complexes shows that Prp8$^{RH}$ undergoes a 145° rotation from the *sc*B$^{act}$ to *sc*B* state, while it undergoes a -35° rotation in the opposite direction from the *sc*B$^{act}$ to the *sp*B$^{d}$-II state. This position of the Prp8$^{RH}$ in the *sp*B$^{d}$-II complex is fixed by the Gpl1 knob (Extended Data Fig. 8d). For simplicity, the Prp8$^{N\text{-}domain}$, Prp8$^{RT}$, Prp8$^{Thumb/X}$, Prp8$^{Linker}$ and Prp8$^{En}$ domains of the *sp*B$^{d}$ complex are shown in cartoon representation, while Prp8$^{RH}$ from *sc*B$^{act}$ (purple), Prp8$^{RH}$ from *sc*B* (teal blue) and Prp8$^{RH}$ from *sp*B$^{d}$-II complex (pale blue) are shown as surfaces. Structure alignment is performed on the Prp8 Large domain comprising the N-domain till the En domain.

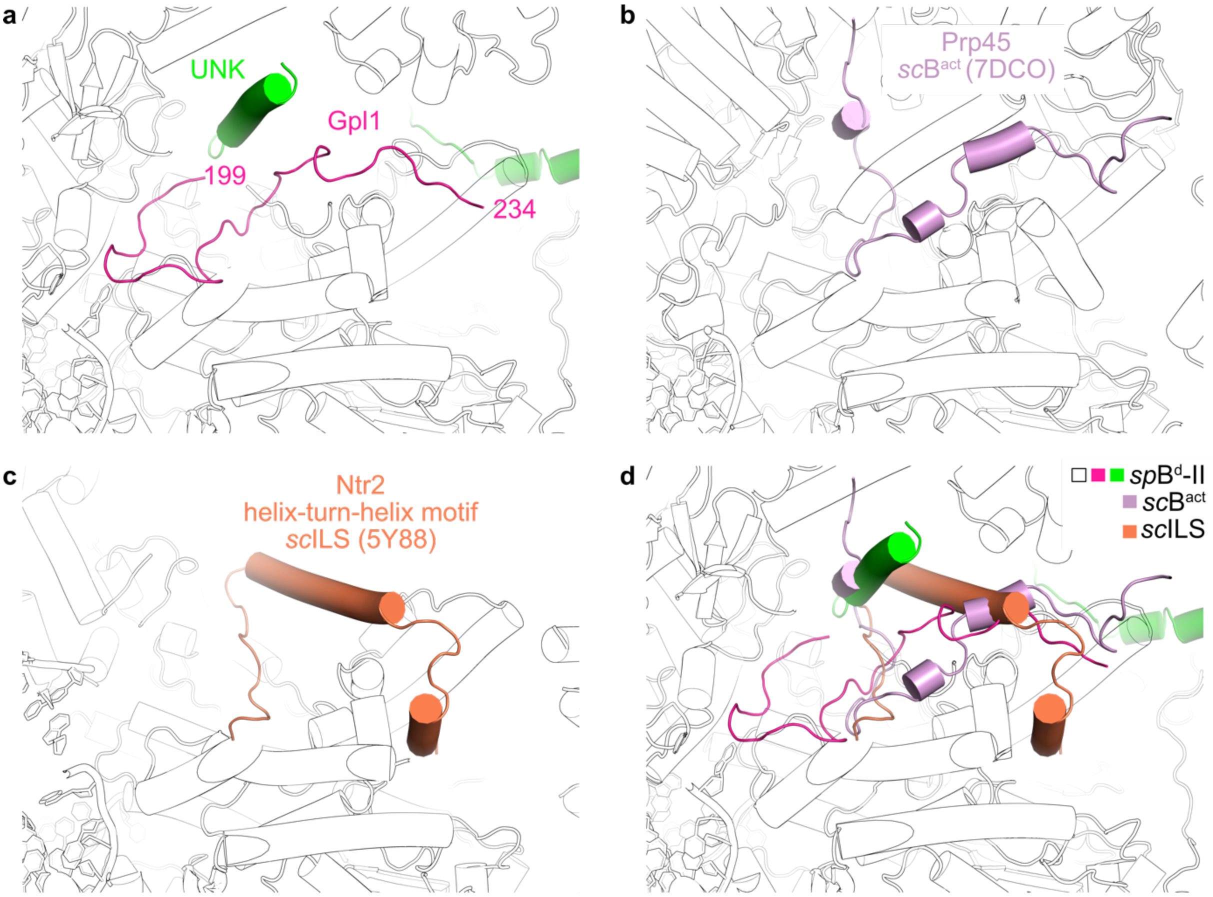


**Extended Data Fig. 10: Comparison of Gpl1 (residues 199-234) in the *sp*B$^{d}$-II complex with Prp45 and Ntr2 as part of the *sc*B$^{act}$ and *sc*ILS complexes, respectively, indicates an overlapping binding site on Prp8.**

**a,** Positions of Gpl1 residues 199-234 (pink) and UNK (green) in the *sp*B$^{d}$-II complex are shown. **b,** Position of Prp45 (light magenta) in the *sc*B$^{act}$ (PDB: 7DCO) complex is shown in the same view. **c,** Position of the helix-turn-helix motif of Ntr2 (orange) in the *sc*ILS (PDB: 5Y88) complex is shown. **d,** A superposition of the *sp*B$^{d}$-II, *sc*B$^{act}$ and *sc*ILS complexes show that Gpl1 residues 199-234 and UNK in *sp*B$^{d}$-II occupy the same location as stretches of Prp45 and Ntr2 in the *sc*B$^{act}$ and *sc*ILS complexes, respectively. For simplicity, only the *sp*B$^{d}$-II complex is shown in black-and-white outline representation.

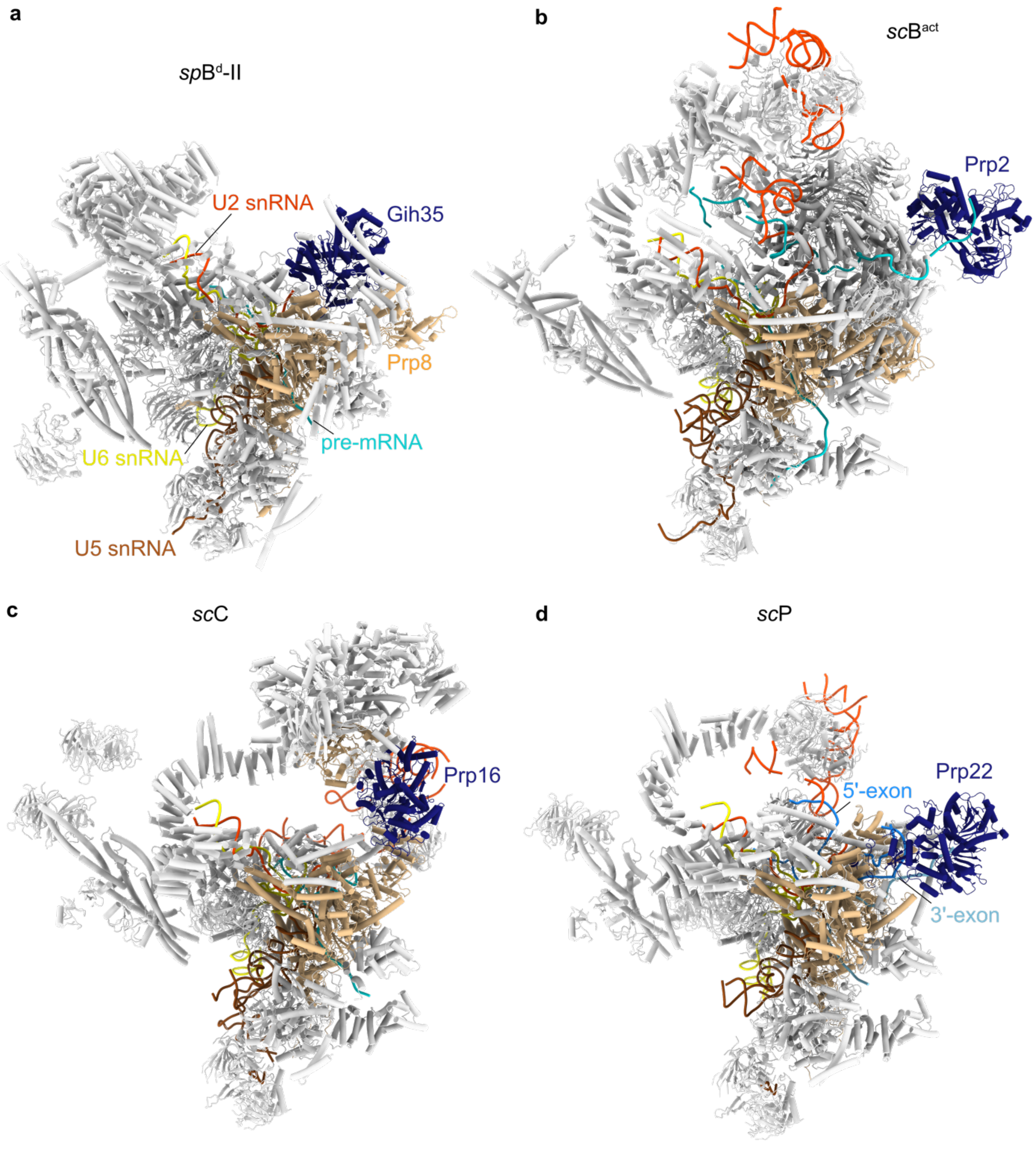


**Extended Data Fig. 11: RNA helicases Gih35, Prp2, Prp16 and Prp22 bind at similar positions at the periphery of the spliceosome.**

Positions of Gih35, Prp2, Prp16 and Prp22 (dark blue) at the periphery of the *sp*B$^{d}$-II complex, the *sc*B$^{act}$ complex (PDB: 7DCO), the *sc*C complex (PDB: 5LJ5), and the *sc*P complex (PDB: 6BK8) are shown in panels **a, b, c and d,** respectively. For orientation, the different RNA species are shown in color and Prp8 is shown in wheat, while the bulk of the spliceosome is shown in gray.

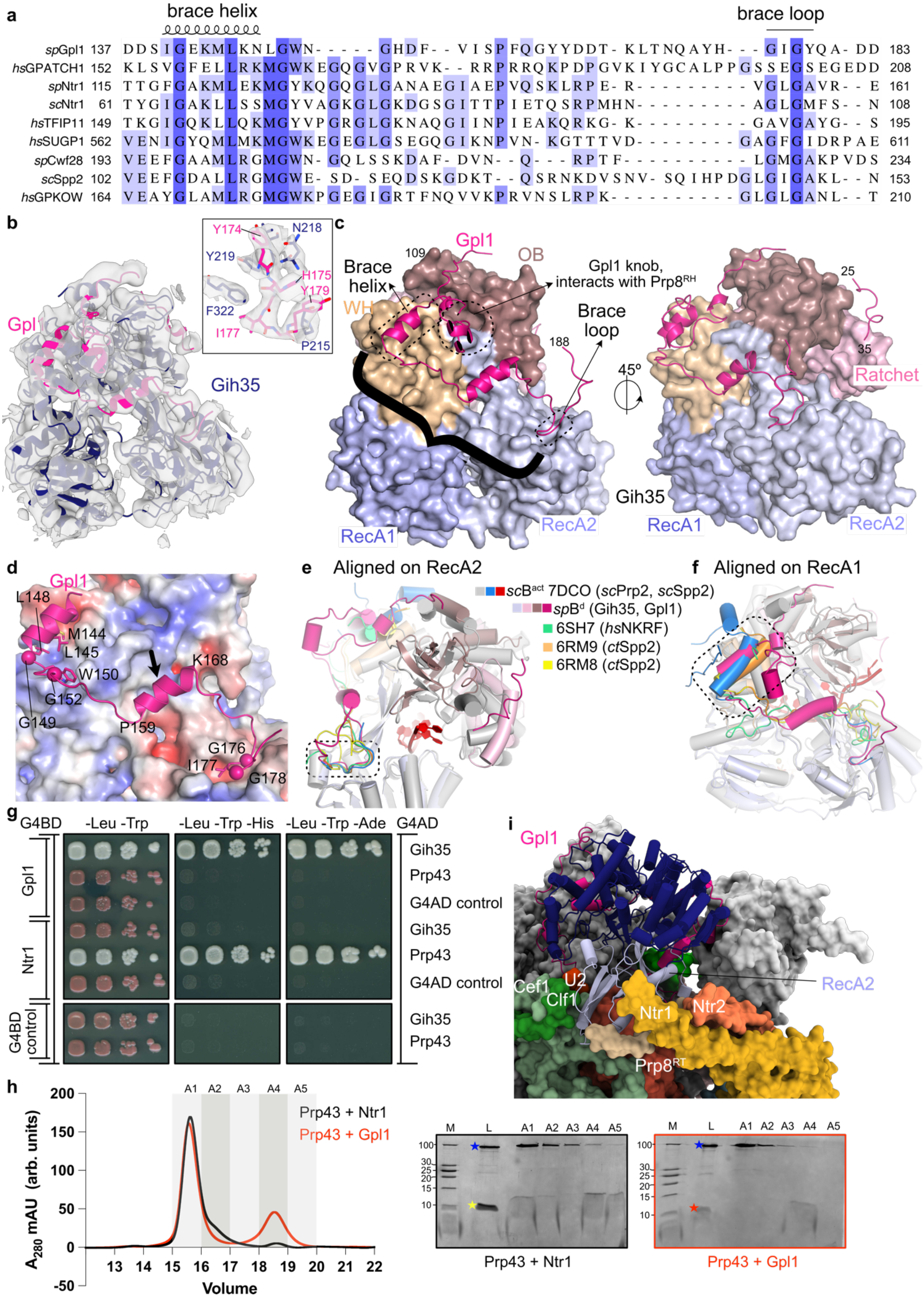


**Extended Data Fig. 12: Gih35-Gpl1 is a helicase – G-patch protein pair.**

**a,** Sequence alignment of G-patch proteins from *S. pombe*, *S. cerevisiae* and *H. sapiens* known or predicted as co-activators of Gih35, Prp43 and Prp2 (or DHX35, DHX15 or DHX16 in humans, respectively). Only the G-patch region of proteins involved in splicing are listed. **b,**

Superimposition of Gih35-Gpl1 (this study) with the cryo-EM map. Inset shows a zoom-in with well-defined side chain densities of Gih35 and the region of Gpl1 encompassing the brace loop. **c,** Overview of the Gih35-Gpl1 complex with the brace helix and brace loop marked with dotted boxes. The Gpl1 knob that interacts with $Prp8^{RH}$ in the *sp*$B^{d}$-II state is marked by a dotted circle. A brace encompassing the brace helix and brace loop is shown in black. **d,** The electrostatic surface potential (±5 kT; red, negative; blue, positive) of Gih35 is plotted. Conserved hydrophobic residues of the Gpl1 G-patch are marked and the Cα atoms of the conserved glycine residues are shown as spheres. The central α-helix in the brace linker is marked with an arrow. **e** and **f,** Comparison of the G-patch interactions. Superposition of the Gih35-Gpl1 complex (*sp*$B^{d}$) with that of *S. cerevisiae* Prp2-Spp2 (*sc*$B^{act}$, PDB: 7DCO), two structures of the *C. thermophilum* Prp2-Spp2 (PDB: 6RM9, 6RM8) and the human DHX15-NKRF complex (PDB: 6SH7). The structures are aligned on the RecA2 domain of the respective helicase in panel (e) and RecA1 domain in panel (f) showing structural conservation in the brace helix and brace loop regions, respectively. For simplicity, only the helicases Gih35 (color coded similar to panel (c)) and *sc*Prp2 (gray) are shown. **g,** Y2H experiments show that Gpl1 interacts with Gih35 but not with Prp43, and Ntr1 interacts with Prp43 but not with Gih35. Full-length Gpl1 or Ntr1 constructs were fused to the Gal4 DNA binding domain (G4BD) while full-length Gih35 and Prp43 were fused to the Gal4 activation domain (G4AD). Auto-activation controls are provided. Serial dilutions of equivalent amounts of yeast were plated on double (-Leu-Trp) and triple dropout media (-Leu-Trp-His, -Leu-Trp-Ade), with growth on triple dropout media indicating an interaction between the tested proteins. Uncropped images are provided as a source data file. **h,** Size-exclusion chromatography profiles of Prp43+Ntr1 G-patch domain (black) and Prp43+Gpl1 G-patch domain (red) are shown (molar ratio Prp43: Ntr1/Gpl1 1:2). Fractions loaded on SDS-PAGE gels are marked. 'arb. units' represent arbitrary units. 1% of sample mixture before loading on size-exclusion column (L) and size-exclusion fractions A1-A5 were separated on 19% SDS-PAGE gels. Positions of Prp43 (blue), Ntr1 G-patch domain (yellow) and Gpl1 G-patch domain (red) are marked with stars. While Prp43 and Ntr1 G-patch domain co-elute in fractions A1-A2, Prp43 and Gpl1 G-patch domain elute separately in fractions A1 and A4, respectively. Excess of Ntr1 G-patch domain elutes in fractions A4-A5. Uncropped gel images are provided as a source data file. **i,** In the *sp*$B^{d}$ -II state, the Gih35 RecA2 domain (light purple) is anchored in a pocket formed by $Prp8^{RT}$, Ntr1, Ntr2, NTC proteins Cef1 and Clf1 and a helix of an unknown protein (light brown).

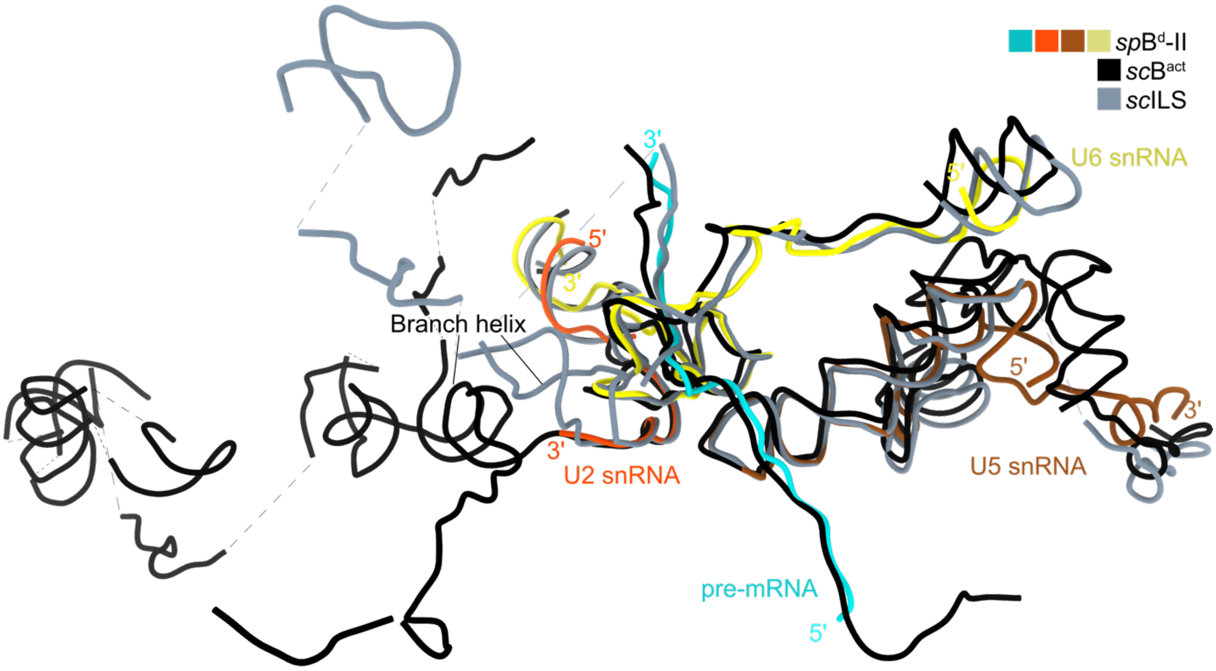


**Extended Data Fig. 13: Comparison of RNA networks between the *sp*B^d^, *sc*B^act^ and *sc*ILS complexes.**

A comparison of the architecture of U2 snRNA, U5 snRNA, U6 snRNA and the pre-mRNA shows overall similarities of the *sp*B$^{d}$ complex with the *sc*B$^{act}$ (PDB: 7DCO) and *sc*ILS (PDB:5Y88) states. The structures are aligned on U6 snRNA.

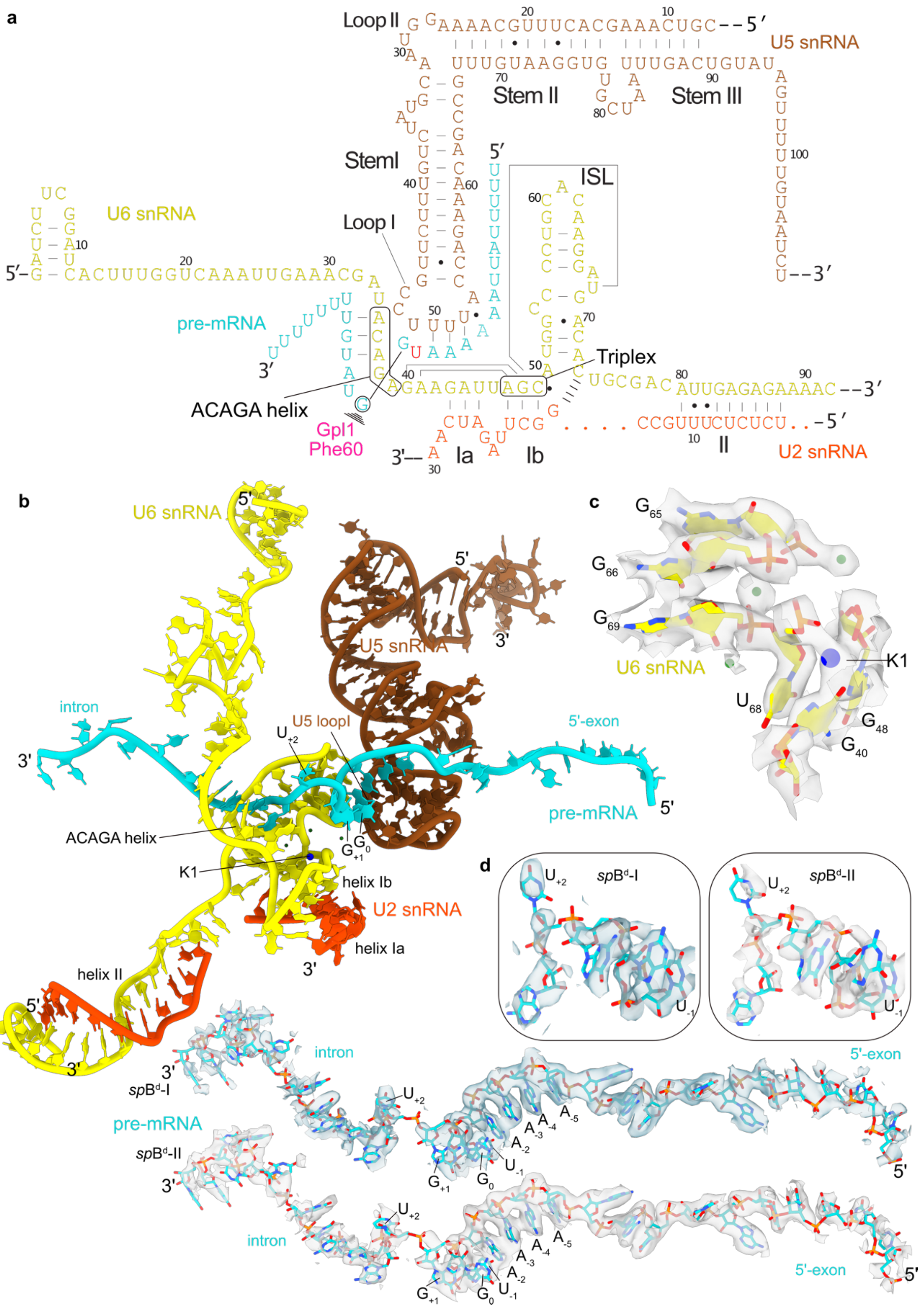


**Extended Data Fig. 14: RNA structure in the *sp*B$^{d}$ complex.**

**a,** Schematic RNA representation for the *sp*B$^{d}$ complex. Gpl1 Phe60, which stacks onto $G_{+1}$ of the pre-mRNA, is shown. **b,** RNA structures in the *sp*B$^{d}$ complex. For orientation, the positions of the U6 ACAGA helix, U5 loop I, U2/U6 helix Ia/Ib, the 5’ss and the catalytic $K^{+}$ ion at the K1 site are marked. **c,** Cryo-EM map (*sp*B$^{d}$-II) shows signals for the $K^{+}$ ion at the K1 site and three other structural $Mg^{2+}$ ions. **d,** Overlays of *sp*B$^{d}$-I and *sp*B$^{d}$-II cryo-EM maps with the pre-mRNA.  Insets show zoom-ins of nucleotides between positions -1 and +3 of the pre-mRNA.

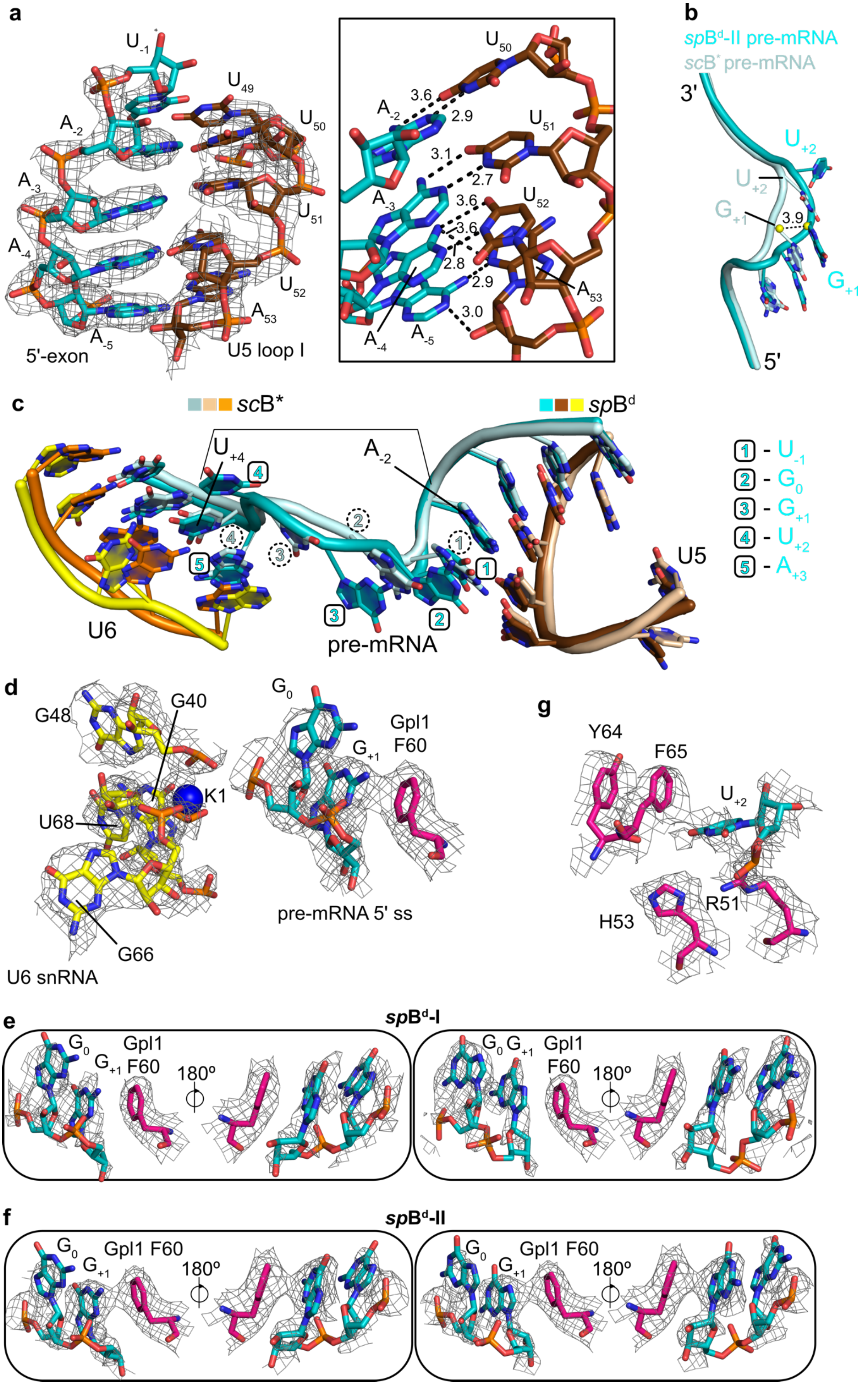

a
$U_{-1}$
$A_{-2}$
$A_{-3}$
$A_{-4}$
$A_{-5}$
5'-exon
$U_{49}$
$U_{50}$
$U_{51}$
$U_{52}$
$A_{53}$
U5 loop I
3.6
2.9
3.1
2.7
3.6
3.6
2.8
2.9
3.0
b
$spB^{d}$-II pre-mRNA
$scB^{*}$ pre-mRNA
3'
5'
$U_{+2}$
$G_{+1}$
3.9
c
$scB^{*}$
$spB^{d}$
$U_{+4}$
$A_{-2}$
U6
pre-mRNA
U5
1 - $U_{-1}$
2 - $G_{0}$
3 - $G_{+1}$
4 - $U_{+2}$
5 - $A_{+3}$
d
G48
G40
K1
U68
G66
U6 snRNA
$G_{0}$
$G_{+1}$
Gpl1
F60
pre-mRNA 5' ss
g
Y64
F65
$U_{+2}$
R51
H53
e
$spB^{d}$-I
$G_{0}$
$G_{+1}$
Gpl1
F60
180º
f
$spB^{d}$-II
Gpl1 F60
180º

**Extended Data Fig. 15: Active site of the *sp*B$^d$ complex.**

**a,** The *sp*B$^d$ cryo-EM map shows that a tight RNA duplex is formed between U5 loop I and 5'-exon nucleotides $A_{-2}A_{-3}A_{-4}A_{-5}$. Inset shows a zoom-in of interactions between U5 loop I and 5'-exon. Hydrogen bonds are shown as dotted lines and distances are indicated in Å. **b,** Insertion of an extra nucleotide at the active site introduces a bend in the pre-mRNA conformation (related to Fig. 3a). Yellow spheres represent centers of backbone sugar rings of *sc*B* pre-mRNA $G_{+1}$ (PDB: 6J6Q) and *sp*B$^d$-II pre-mRNA $G_{+1}$ and the dotted line marks the distance between them as 3.9 Å. **c,** Related to Fig. 3a-c. Superposition of RNA elements (pre-mRNA, U5 and U6) at the active site of *sp*B$^d$ and canonical *sc*B* (PDB:6J6Q) complexes. The bracket marks the positions of pre-mRNA bases $A_{-2}$ and $U_{+4}$ which is covered by 4 nucleotides in *sc*B* (marked with dashed circles) and 5 nucleotides in *sp*B$^d$ (boxed in squares). Using the information from 5'ss sequence logo generated from CNM-sensitive transcripts, the identity of the 5 nucleotides boxed in squares for the *sp*B$^d$ complex is presented. **d,** Gpl1 Phe60 stacks on top of the 5'ss. The cryo-EM map shows density for the $K^+$ ion at the K1 site. **e** and **f,** *sp*B$^d$-I and -II cryo-EM maps superimposed with the 5'ss $G_0$ and $G_{+1}$ and the Gpl1 Phe60, respectively. Since the signal of the pre-mRNA at the active site is weak, alternative conformations (180° flipped) of the $G_0$ and $G_{+1}$ can be modeled. However, in both *sp*B$^d$-I and *sp*B$^d$-II states, both the alternative conformations of $G_0$ and $G_{+1}$ stack well with Gpl1 Phe60. **g,** $U_{+2}$ of the intron is sequestered into a pocket formed by Gpl1.

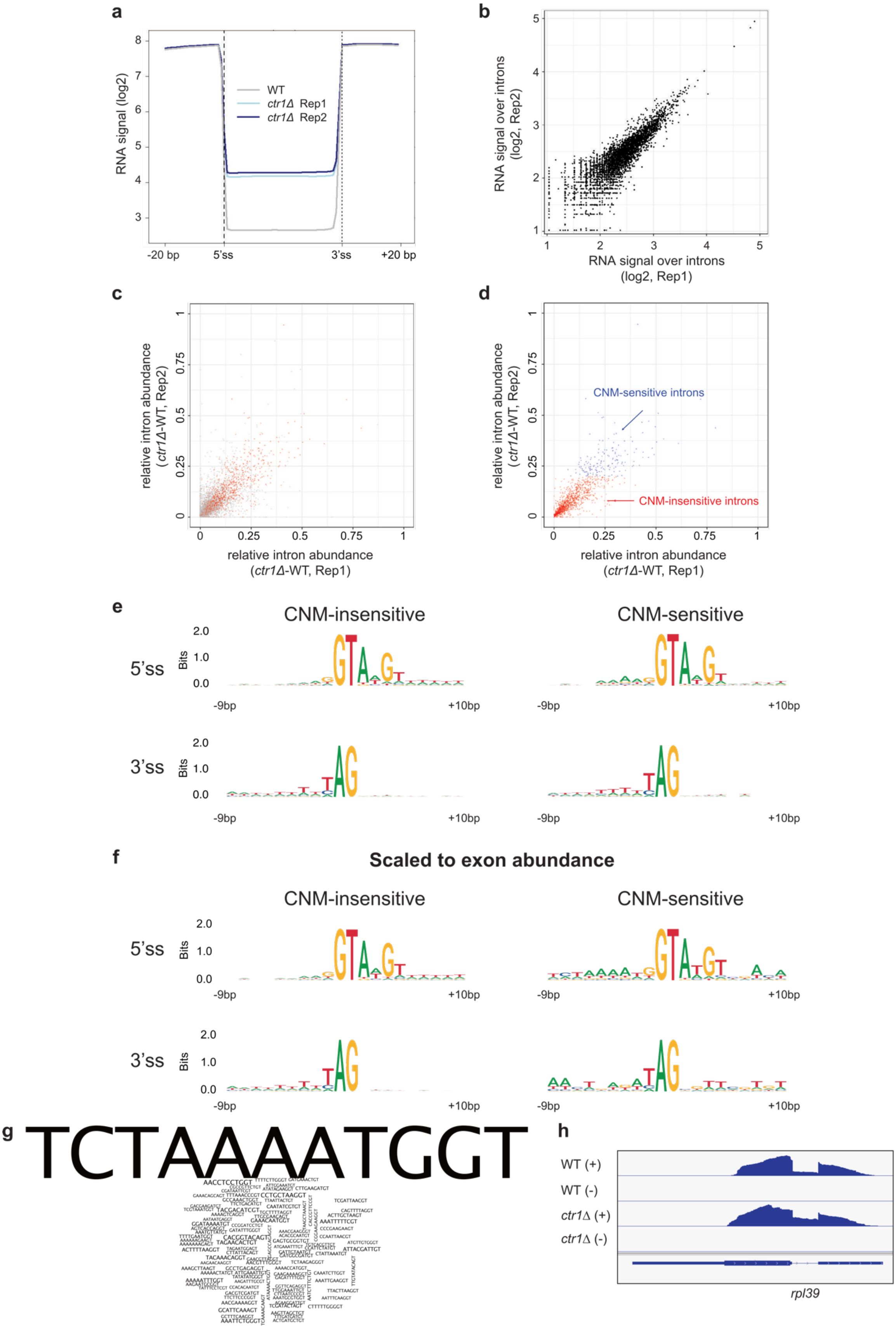


**Extended Data Fig. 16: Bioinformatics analysis of CNM-sensitive and CNM-insensitive introns.**

**a,** Metagene profile of sense RNA expression levels in the indicated strains (WT, *ctr1*Δ) for all annotated *S. pombe* introns. The geometric average of RNA expression is shown from 20 bp upstream to 20 bp downstream of 5' ss and 3' ss of the introns. **b,** Scatter plot showing average RNA coverage of all annotated introns of *S. pombe* genes, based on RNA signal of two biological replicates of the *ctr1*Δ strain. **c-d,** Scatter plots representing relative intron abundance in *ctr1*Δ compared to WT strains. **c,** Introns with reproducible RNA coverage (> 30 normalized read count in both biological replicates) are shown in red. **d,** Reproducible introns classified into CNM-sensitive sites and CNM-insensitive sites. CNM-sensitive sites are defined as introns having at least 20% increase in intron abundance in *ctr1*Δ relative to WT. CNM-insensitive and CNM-sensitive introns are shown in red and blue, respectively. Intron abundance is expressed as the ratio of the RNA coverage over the intron and the arithmetic average of those of the neighboring exons. Complete lists of all CNM-sensitive and CNM-insensitive sites are provided in Supplementary Table 2. **e-f,** Sequence logos of 5'ss and 3'ss of CNM-sensitive and CNM-insensitive sites shown with a 10 bp upstream and downstream flanking regions. Logos are derived based on unscaled ("per gene") occurrences (**e**) or being scaled to be proportional to the normalized coverage of the upstream exon (**f**). At each position, the total height of the nucleotides expresses the information content in bits. **g,** Word cloud showing the 5'ss of CNM-sensitive introns together with 9 bp upstream of the 5'ss. The height of the letters is proportional to normalized coverage of the upstream exon. **h,** IGV[4] snapshot of strand-specific RNA-seq coverage of the *rpl39* gene, the most highly expressed mRNA with a CNM-sensitive intron, in the indicated strains.

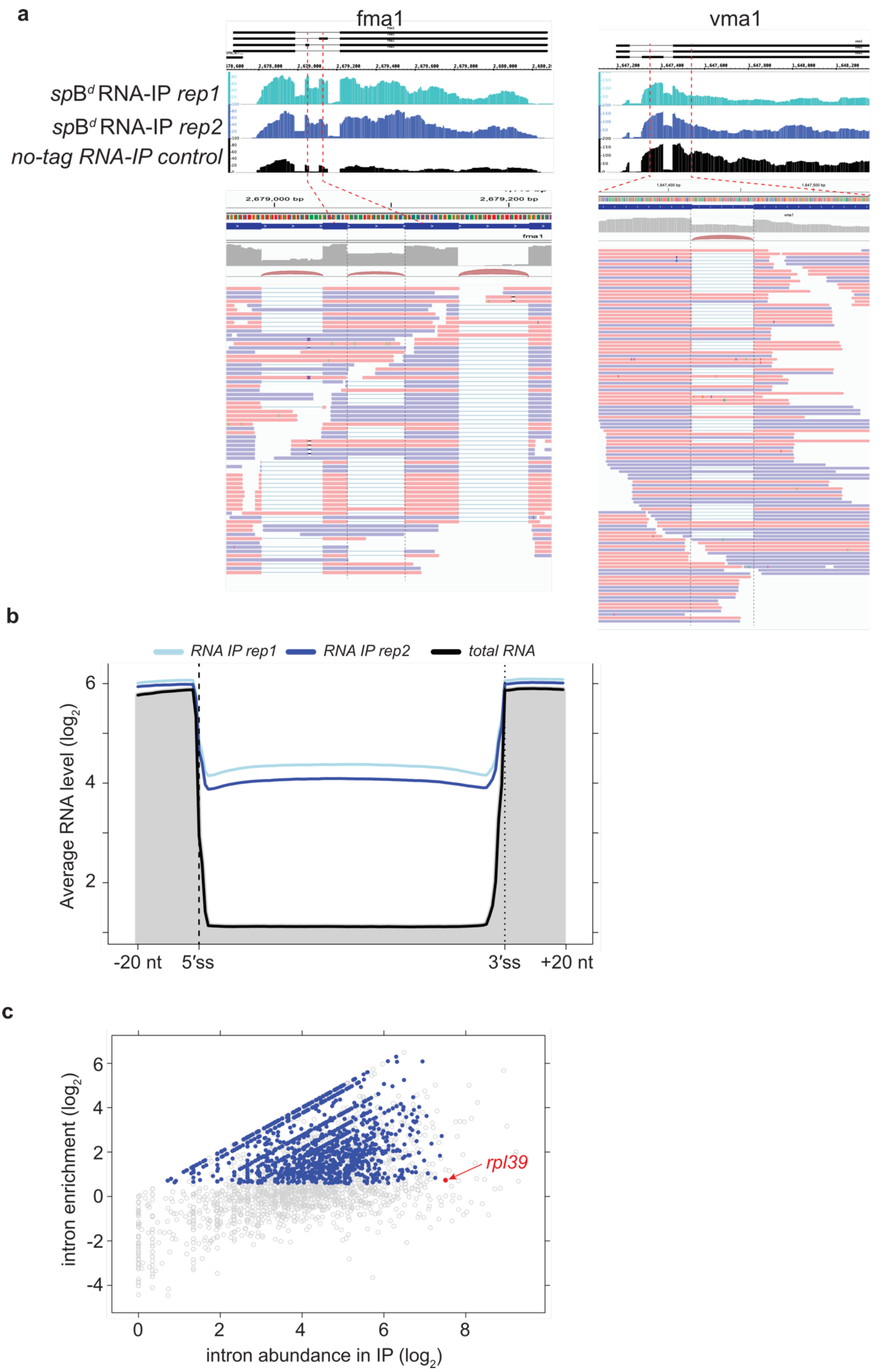
a
fma1
vma1
spB^d RNA-IP rep1
spB^d RNA-IP rep2
no-tag RNA-IP control
2,679,000 bp
2,679,200 bp
fma1
b
RNA IP rep1
RNA IP rep2
total RNA
Average RNA level (log2)
6
4
2
-20 nt
5′ss
3′ss
+20 nt
c
rpl39
intron enrichment (log2)
6
4
2
0
-2
-4
0
2
4
6
8
intron abundance in IP (log2)

**Extended Data Fig. 17**: ***sp*B$^d$ spliceosome RNA immunoprecipitation (RNA-IP).**

**a,** Gene browser view of strand-specific RNA levels at the *fma1* and *vma1* genes show representative examples of retained introns in the *sp*B$^d$ spliceosome RNA-IP samples (light blue and dark blue tracks represent biological replicates 1 and 2, respectively). Black track shows the no-tag control for the RNA-IP, representing the background signal. All datasets are coverage normalised. Intronic reads at affected introns are strongly enriched in the *sp*B$^d$ spliceosome purification compared to no-tag RNA-IP background. The lower panel shows the magnified view of individual strand-specific paired-end reads in the *sp*B$^d$ RNA-IP *rep1* sample. Pink lines represent read 1 (forward read), blue lines represent read 2 (reverse reads). These representative examples show that most intronic reads in the affected introns run across exon-intron boundaries, confirming that pre-mRNA transcripts are uncleaved in *sp*B$^d$ spliceosome preparations. **b,** Metagene profile of sense RNA levels in the *sp*B$^d$ spliceosome purification (light blue and dark blue lines represent biological replicates 1 and 2, respectively) and in total RNA sequencing of wild-type *S. pombe* strain (black line and grey shaded area), for all annotated *S. pombe* introns. The geometric average of RNA levels is shown from 20 nt upstream to 20 nt downstream of 5’ ss and 3’ ss of introns. To allow better comparison of intron coverage, datasets were normalized for the total exon coverage of all annotated ORFs in *S. pombe*. **c,** Scatter plot of $\log_2$ intron coverage (X-axis) versus $\log_2$ intron enrichment, compared to no-tag RNA-IP background (Y-axis). Dots represent all individual introns annotated in the *S. pombe* genome. Blue dots represent enriched introns in the RNA-IP (min 1.5x enriched compared to background) that undergo genuine splicing (contain at least 25% spliced reads) and coverage at the 3’ end of the intron is at least 50% of its 5’ coverage (to filter out alternatively spliced and mis-annotated introns). Red dot represents intron 1 in *rpl39* gene. Note that RNA-IP dataset is not ideal to determine absolute abundance values, due to the high level of contaminating RNAs and the relatively high coverage values of mis-annotated, alternatively spliced or retained introns, compared to genuinely spliced introns.

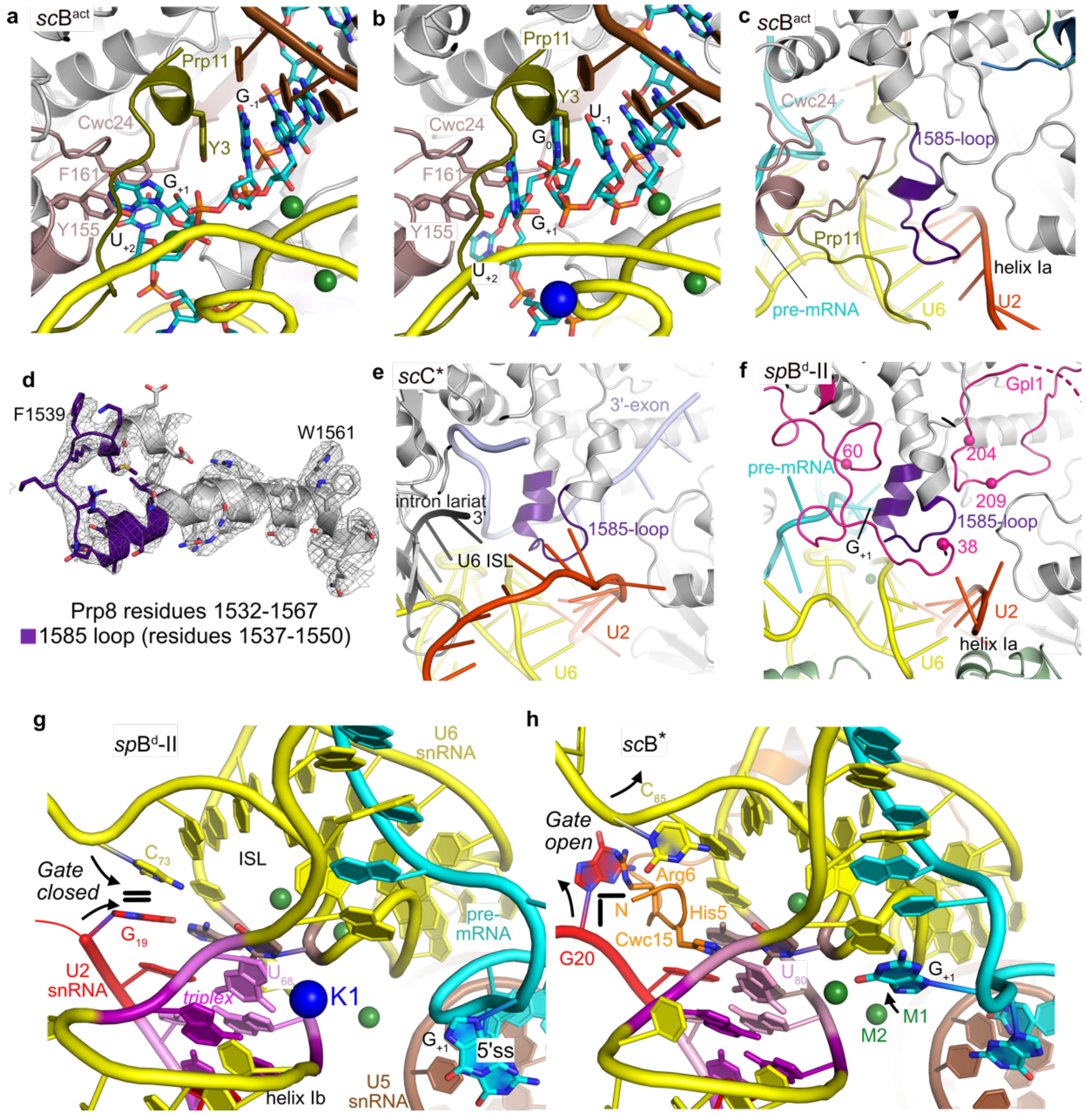


**Extended Data Fig. 18: Comparisons of *sp*B$^{d}$ active site with that of *sc*B$^{act}$, scB* and scC*.** **a,** Prp11 and Cwc24 are bound to the active site in the *sc*B$^{act}$ complex (PDB: 7DCO). Prp11 Tyr3 stacks with $G_{-1}$ and Tyr155, and Phe161 of Cwc24 stack with $G_{+1}$ and $U_{+2}$. **b,** Superposition of Prp11 and Cwc24 from the *sc*B$^{act}$ complex onto the *sp*B$^{d}$ complex shows steric clashes between the pre-mRNA and Prp11/Cwc24. For simplicity, Gpl1 is not shown. **c,** The 1585-loop of Prp8 interacts with Prp11, Cwc24 and the U2/U6 duplex in the *sc*B$^{act}$ complex (PDB: 7DCO). **d,** The 1585-loop of Prp8 comprising residues 1537-1550 (purple) in *S. pombe* is structured in the *sp*B$^{d}$ complex as seen by the overlay with the cryo-EM map. **e,** Position of the 1585-loop in the *sc*C* complex (PDB: 5WSG) is similar to that of the *sp*B$^{d}$ complex. **f,** In the *sp*B$^{d}$ complex, the 1585-loop is held in position by Gpl1 residues 38-60 on one side and residues 204-209 on the other. It also directly interacts with U2/U6 helix Ia. **g** and **h,** A gating mechanism for entry of the N-terminus of Cwc15 into the active site. **g,** In the *sp*B$^{d}$ complex, $C_{73}$ of the U6 snRNA stacks onto the $G_{19}$ of U2 snRNA (part of U2/U6 helix Ib). Due to the stacking, the entrance 'gate' for the N-terminus of Cwc15 into the active site (characterized by three conserved base triples: triplex; including the uridine ($U_{68}$) coordinating the K1 site) is blocked. **h,** In the *sc*B* complex (PDB: 6J6H), the 'gate' is open and the N-terminus of Cwc15 can enter into the active site where it stabilizes the catalytic triplex. The positions of the catalytic metal ions M1 (behind $G_{+1}$, indicated with an arrow) and M2 are marked.

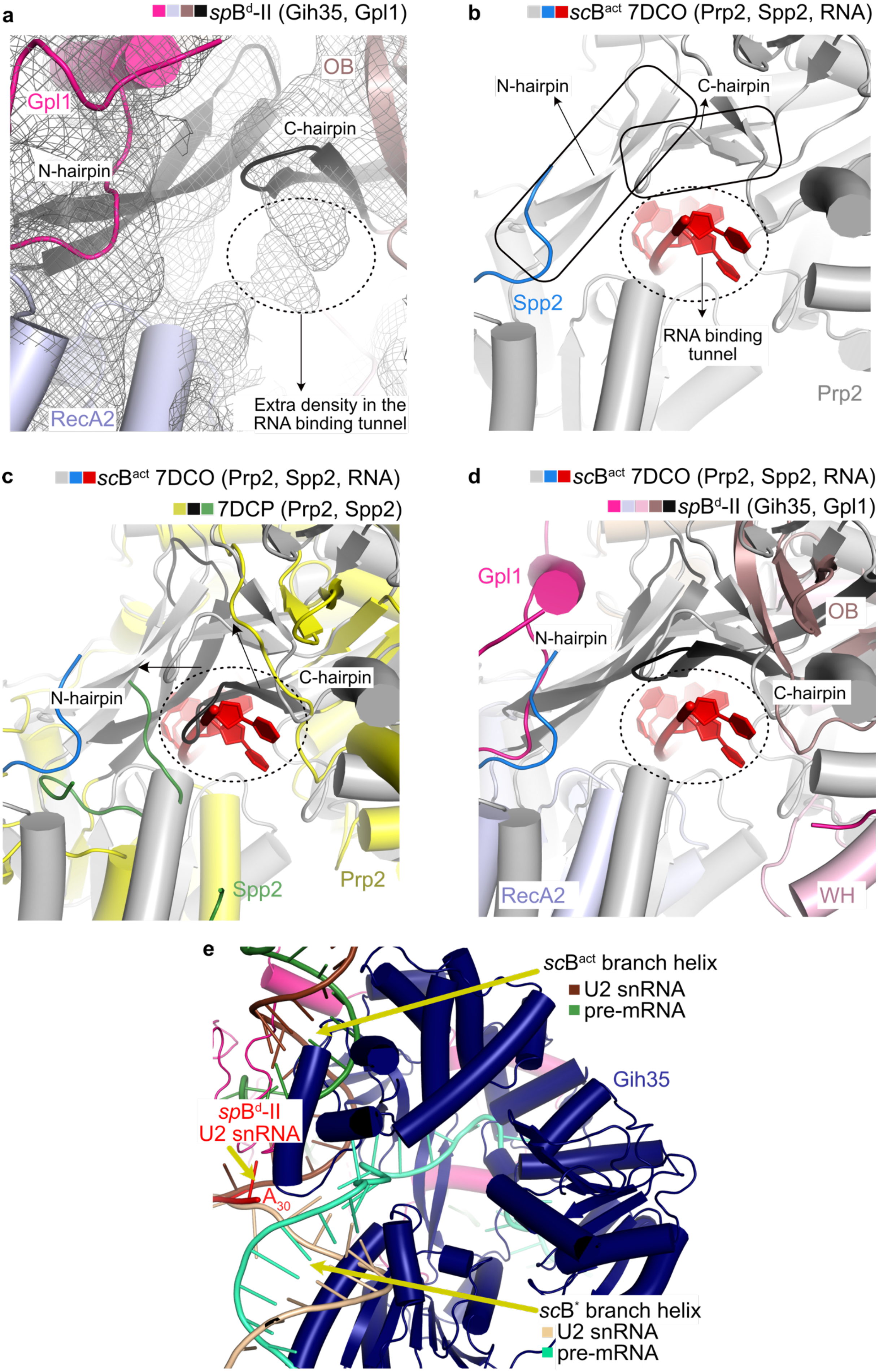
a
spBd-II (Gih35, Gpl1)
Gpl1
OB
C-hairpin
N-hairpin
RecA2
Extra density in the RNA binding tunnel
b
scBact 7DCO (Prp2, Spp2, RNA)
N-hairpin
C-hairpin
Spp2
RNA binding tunnel
Prp2
c
scBact 7DCO (Prp2, Spp2, RNA)
7DCP (Prp2, Spp2)
C-hairpin
N-hairpin
Spp2
Prp2
d
scBact 7DCO (Prp2, Spp2, RNA)
spBd-II (Gih35, Gpl1)
Gpl1
N-hairpin
OB
C-hairpin
RecA2
WH
e
scBact branch helix
U2 snRNA
pre-mRNA
Gih35
spBd-II
U2 snRNA
A30
scB* branch helix
U2 snRNA
pre-mRNA

**Extended Data Fig. 19: RNA binding to Prp2 in the *sc*B$^{act}$ complex and to Gih35 in the *sp*B$^{d}$ complex.**

**a,** *sp*B$^{d}$-II cryo-EM map shows density likely corresponding to an RNA substrate bound to Gih35. **b,** RNA-binding to Prp2. The N-and C-hairpins of Prp2 prevent the backsliding of pre-mRNA ([5], PDB: 7DCO). The RNA binding tunnel is outlined with dashes. **c,** Structure superposition of Spp2-Prp2 in the absence (PDB: 7DCP) and presence of RNA (PDB: 7DCO) show that the N- and C-hairpins undergo large movements to accommodate the RNA. The structures are aligned on the RecA1 domains. **d,** Structure superposition of the Gih35-Gpl1 complex with Prp2-Spp2-RNA shows that the position of the N- and C-hairpins of Gih35 in the *sp*B$^{d}$ complex are compatible with RNA binding. **e,** A close-up view of the last nucleotide of U2 snRNA visible in the *sp*B$^{d}$ complex ($A_{30}$) positioned close to the RNA binding tunnel of Gih35. Superpositions with the *sc*B$^{act}$ complex (PDB: 7DCO) and the *sc*B* complex (PDB: 6J6H) show that the branch helix from either state would clash with Gih35. Given that Gih35 is bound to an RNA substrate and the branch helix is invisible in our structure, it is likely that Gih35 acts on the branch helix.

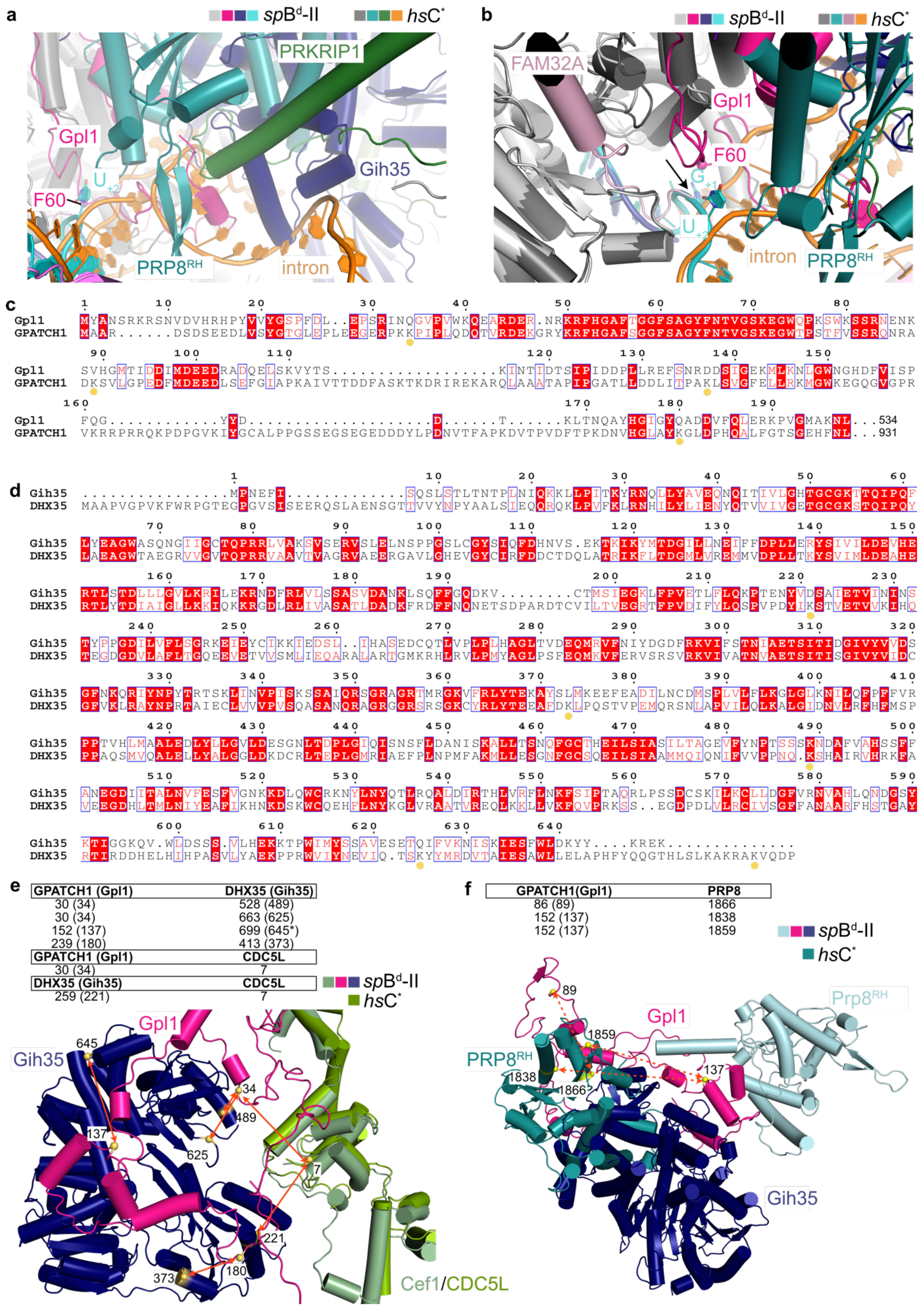


**Extended Data Fig. 20: Comparison between Gih35-Gpl1 in the *sp*B$^{d}$ complex and DHX35-GPATCH1 in the *hs*C* complex**

**a,** Superposition of *sp*B$^{d}$-II state with *hs*C* complex. For simplicity, only PRKRIP1 (green), PRP8$^{RH}$ (teal) and intron (orange) from *hs*C* complex are shown in color. Gpl1 and Gih35 from *sp*B$^d$ complex clash with PRKRIP1 and Prp8$^{RH}$ from *hs*C* complex. For orientation, pre-mRNA $U_{+2}$ and Gpl1 Phe60 (Cα atom as sphere) are marked. **b,** FAM32A is at the catalytic center in *hs*C* (marked by a black arrow), which is incompatible with the pre-mRNA 5'ss and Gpl1 in the *sp*B$^d$ complex. **c and d,** Sequence alignments of Gpl1 with GPATCH1 and Gih35 with DHX35, respectively. Yellow dots mark the lysine residues of GPATCH1/DHX35 which crosslink to other proteins as listed in panels (e) and (f). **e,** All crosslinks between GPATCH1 and DHX35 reported for *hs*C* complex[3] are satisfied (distance between Cα atoms of crosslinked residues < 24 Å) in the *sp*B$^d$ complex using corresponding amino acid positions from *S. pombe* orthologues (listed in brackets, shown as red arrows). Cef1 and CDC5L superimpose well and the crosslinks from CDC5L to Gpl1/Gih35 are also satisfied. Cα atoms for the corresponding residues from Gpl1, Gih35 and CDC5L are shown as yellow spheres. **f,** All crosslinks between GPATCH1 and Prp8$^{RH}$ reported for *hs*C* complex[3] are unsatisfied (distance between Cα atoms of crosslinked residues > 24 Å, marked by dashed red arrows) in the *sp*B$^d$-II complex due to the differences in location of Prp8$^{RH}$ between *sp*B$^d$-II complex and *hs*C* complex.

### Extended Data Table 1: Summary of model building for the *sp*B$^d$-I (PDB ID: 9ESH) and *sp*B$^d$-II (PDB ID: 9ESI) complexes

| | **Protein/RNA component (*S. pombe*/ *S. cerevisiae/H. sapiens*)** | **Total length** | **Modeling template** | **Modeled region in *sp*B$^d$-I (including gaps)** | **Modeled region in *sp*B$^d$-II (including gaps)** | ***sp*B$^d$-I Chain ID** | ***sp*B$^d$-II Chain ID** |
|---|---|---|---|---|---|---|---|
| **Pre-mRNA** | Pre-mRNA | - | - | 29 (5'exon and intron) | 29 (5'exon and intron) | 1 | 1 |
| **U5 snRNP** | U5 snRNA | 120 nts | 3JB9 | 7-108 | 7-108 | 5 | 5 |
| | Spp42/Prp8/PRP8 | 2363 | | 45-1781 | 45-2030 | A | A |
| | Cwf10/Snu114/SNU114 | 984 | | 67-984 | 67-984 | B | B |
| | Cwf17/-/SNRNP40 | 340 | | 36-338 | 36-338 | C | C |
| | SmD3 | 97 | | 2-97 | 2-97 | D | D |
| | SmB | 147 | | 3-118 | 3-118 | E | E |
| | SmD1 | 117 | | 2-82 | 2-82 | F | F |
| | SmD2 | 115 | | 5-115 | 5-115 | G | G |
| | SmE | 84 | | 5-84 | 5-84 | H | H |
| | SmF | 78 | | 3-75 | 3-75 | I | I |
| | SmG | 77 | | 3-75 | 3-75 | J | J |
| **U6 snRNP** | U6 snRNA | 99 nts | 3JB9 | 1-92 | 1-92 | 6 | 6 |
| **U2 snRNP** | U2 snRNA | 186 nts | 3JB9 | 3-30 | 3-30 | 2 | 2 |
| **NTC core** | Cwf4/Clf1/SYF3 | 674 | 3JB9 + AF* | 11-620 | 11-620 | R | R |
| | Prp19/Prp19/PRP19 | 488 | | 1-132,1-134,1-135,1-131 | 1-132,1-134,1-135,1-131 | S,T,U,V | S,T,U,V |
| | Cdc5/Cef1/CDC5L | 757 | | 4-757 | 4-757 | W | W |
| | Cwf3/Syf1/SYF1 | 790 | | 63-735 | 63-735 | X | X |
| | Cwf7/Snt309/SPF27 | 187 | | 13-185 | 13-185 | Z | Z |
| | Syf2/Syf2/SYF2 | 229 | 6J6Q | 83-215 | 83-184 | Y | Y |
| **NTC related** | Prp5/Prp46/PRL1 | 473 | 3JB9 | 83-473 | 83-473 | K | K |
| | Prp45/Prp45/SKIP | 557 | | 82-332 | 82-332 | L | L |
| | Cwf5/Ecm2/RBM22 | 354 | | 16-305 | 16-305 | M | M |
| | Cwf11/-/Aquarius | 1284 | AF | 1-1284 | 1-1284 | N | N |
| | Cwf14/Bud31/G10 | 146 | 3JB9 | 3-146 | 3-146 | O | O |
| | Cwf2/Cwc2/RBM22 | 388 | | 47-328 | 47-328 | P | P |
| | Cwf15/Cwc15/CWC15 | 265 | | 24-265 | 24-265 | Q | Q |
| **Splicing factors** | Prp17/Prp17/CDC40 | 558 | 3JB9 | 9-160 | 9-160 | a | a |
| | Cwf21/Cwc21/SRRM2 | 293 | 5LJ3 + AF | 2-145 | 2-145 | b | b |
| | Cwf22/Cwc22/CWC22 | 887 | 5MQF + AF | 404-607 | 404-607 | c | c |
| | Cyp1/-/PPIL1 | 155 | 3JB9 | 2-155 | 2-155 | d | d |
| | Bis1/-/ESS2 | 384 | AF | - | 22-262 | - | e |
| | Saf4/Yju2/CCDC194 | 299 | AF | - | 156-215 | - | p |
| **Ntr1 complex** | Ntr1/Ntr1/TFIP11 | 797 | 5Y88 + AF | 717-797 | 264-797 | m | m |
| | Ntr2/Ntr2/- | 361 | 5Y88 + AF | - | 296-325 | - | n |
| **Gih35-Gpl1** | Gih35/-/DHX35 | 647 | AF | 20-645 | 20-645 | z | z |
| | Gpl1/-/GPATCH1 | 534 | *de novo* + 6HS6 + 7DCP | 25-234 | 25-234 | y | y |
| **Unknown** | UNK | - | - | - | - | - | q |
| | Unknown (NTC) | - | - | - | - | r | r |
| | Unknown (around Gih35) | - | - | - | - | f | f |

*AF: Alphafold model

**Extended Data Table 2: Cryo-EM data collection, refinement and validation statistics**

| | *sp*B$^{d}$-I complex (EMDB: EMD-19941; PDB: 9ESH) | *sp*B$^{d}$-II complex (EMDB: EMD-19942; PDB: 9ESI) |
|---|---|---|
| **Data Collection and Processing** | | |
| Detector | K3 | |
| Magnification | 105,000 | |
| Voltage (kV) | 300 | |
| Electron exposure (e-/Å$^2$) | 49.4 | |
| Defocus range (µm) | -0.6 to -1.8 | |
| Pixel size (Å) | 0.822 | |
| Symmetry imposed | C1 | |
| Initial partical images (no.) | 929,930 | |
| Final particle images (no.) | 61,423 | 72,631 |
| Map resolution (Å) | 3.2 | 3.1 |
| FSC threshold | 0.143 | 0.143 |
| Map resolution range (Å) | 2.8-23 | 2.7-24 |
| **Refinement** | | |
| Initial model used (PDB code) | n/a | n/a |
| Model resolution (Å) | 3.1 | 3.1 |
| FSC threshold | 0.143 | 0.143 |
| Model resolution range (Å) | - | - |
| Model-map scores | | |
| $CC_{mask}$ | 0.81 | 0.82 |
| $d_{FSC\ model}$ (0.143/0.5) (masked) | 3.1/3.4 | 3.1/3.3 |
| Map sharpening *B* factor (Å$^2$) | 48.1 | 43.8 |
| **Model Composition** | | |
| Non-hydrogen atoms | 90,868 | 98,220 |
| Protein residues | 10,603 | 11,468 |
| RNA/DNA | 247 | 247 |
| Ligands | 13 | 13 |
| ***B* factors (Å$^2$)** | | |
| Protein | 162 | 165 |
| RNA/DNA | 147 | 164 |
| Ligand | 183 | 165 |
| **RMSDs** | | |
| Bond lengths (Å) | 0.004 | 0.007 |
| Bond angles (°) | 0.779 | 0.799 |
| **Validation** | | |
| Molprobity score | 1.99 | 1.98 |
| Clash score | 13.53 | 14.32 |
| Poor rotamers (%) | 0.01 | 0.01 |
| **Ramachandran plot** | | |
| Favored (%) | 94.88 | 95.40 |
| Allowed (%) | 5.00 | 4.42 |
| Disallowed (%) | 0.11 | 0.18 |

### References


1 Wan, R., Bai, R., Yan, C., Lei, J. & Shi, Y. Structures of the Catalytically Activated Yeast Spliceosome Reveal the Mechanism of Branching. *Cell* 177, 339-351 e313 (2019). https://doi.org:10.1016/j.cell.2019.02.006

2 Wan, R., Yan, C., Bai, R., Lei, J. & Shi, Y. Structure of an Intron Lariat Spliceosome from Saccharomyces cerevisiae. *Cell* 171, 120-132 e112 (2017). https://doi.org:10.1016/j.cell.2017.08.029

3 Dybkov, O. *et al.* Regulation of 3' splice site selection after step 1 of splicing by spliceosomal C* proteins. *Sci Adv* 9, eadf1785 (2023). https://doi.org:10.1126/sciadv.adf1785

4 Thorvaldsdóttir, H., Robinson, J. T. & Mesirov, J. P. Integrative Genomics Viewer (IGV): high-performance genomics data visualization and exploration. *Briefings in bioinformatics* 14, 178-192 (2013).

5 Bai, R. *et al.* Mechanism of spliceosome remodeling by the ATPase/helicase Prp2 and its coactivator Spp2. *Science* 371 (2021). https://doi.org:10.1126/science.abe8863
